# An ancestral mitochondrial DNA insertion disrupts RNAi and enables persistence of a novel mycovirus in *Cryptococcus neoformans*

**DOI:** 10.64898/2026.09.15.751489

**Authors:** Jun Huang, Connor J. Larmore, Timothy C. Davenport, Anna Floyd Averette, Yeseul Choi, Humberto Debat, Purav Gupta, Artem Babaian, Marc D. Meneghini, Sheng Sun, Joseph Heitman

## Abstract

RNA interference (RNAi) is a widely conserved genome-defense mechanism that protects eukaryotes against viruses and transposable elements. We previously showed that RNAi loss can lead to hypermutation and antifungal drug resistance in the human fungal pathogen *Cryptococcus neoformans*, illuminating the potential clinical relevance of this pathway. In this study, we identified another function of RNAi in *C. neoformans*: mycovirus restriction. By screening known RNAi-deficient *C. neoformans* isolates, we identified a novel dsRNA mycovirus of the *Orthototiviridae* family, which we named CnTV1. We subsequently detected CnTV1 in three additional RNAi-deficient isolates. All CnTV1-positive isolates shared an ancestral nuclear mitochondrial DNA segment (NUMT) insertion that disrupts the gene encoding Argonaute (Ago1). Restoration of RNAi in sexually produced zygotes efficiently prevented CnTV1 inheritance by progeny. RNAi rescue by CRISPR-Cas9-mediated allele exchange eliminated CnTV1 during vegetative growth, further demonstrating RNAi is sufficient for mycoviral control. Loss of the RNA helicase Ski2 or the exoribonuclease Xrn1 increased CnTV1 abundance, revealing RNAi-independent restriction of the virus. Restoration of RNAi cleared the virus even in the absence of Ski2 or Xrn1, indicating a dominant role for RNAi in antiviral defense. To investigate the biological implications of CnTV1 infection, we developed a cytoplasmic-mixing approach and generated isogenic strain pairs in an RNAi-deficient background that differ only in viral infection status. Leveraging these strains, we show that CnTV1 infection leads to coordinated, low-magnitude transcriptomic changes and that strains lacking the mycovirus were moderately less virulent in a murine infection model. Our findings reveal that RNAi serves as a dominant antiviral defense system in *C. neoformans* and suggest that naturally occurring RNAi deficiency may be more prevalent than previously appreciated. This work highlights mycovirus persistence as an important consequence of RNAi loss in this WHO-designated critical priority fungal pathogen.

**Significance:** Despite its broad conservation, RNAi has been repeatedly lost across diverse fungal lineages, yet the biological consequences of RNAi loss remain poorly understood. We previously found that RNAi loss leads to hypermutation and antimicrobial drug resistance in the human fungal pathogen *Cryptococcus neoformans*. Here, we show that an ancestral mitochondrial DNA insertion disrupts RNAi and thereby enables the persistence of a novel dsRNA mycovirus in natural *C. neoformans* isolates. RNAi and cytoplasmic RNA decay pathways both restrict the mycovirus, with RNAi providing the dominant antiviral defense. Mycovirus infection introduces modest but coordinated transcriptional changes and moderately impacts virulence in mice. Together, these findings reveal mycovirus persistence as a consequence of natural RNAi loss and establish RNAi as a central antiviral defense system in *C. neoformans*.

## Introduction

RNA interference (RNAi) is an evolutionarily conserved gene-silencing mechanism that plays diverse roles in genome defense, gene regulation, and antiviral immunity across eukaryotes (1–6). Double-stranded RNA recognition and cleavage by Dicer, amplification by RNA-dependent RNA polymerase (RdRp), and sequence-specific targeting by Argonaute are key components of the RNAi machinery (3). In fungi, RNAi protects genome integrity by suppressing transposable elements and silencing foreign nucleic acids (5–7). Despite these functions, RNAi has been repeatedly and independently lost across diverse fungal lineages, presenting an evolutionary paradox (8–10). For example, the model budding yeast *Saccharomyces cerevisiae* lacks key RNAi components, including Dicer and Argonaute, yet introduction of functional Dicer and Argonaute proteins can reconstitute RNAi activity in this species (11). Notably, reconstituting RNAi also drives loss of the resident L-A totivirus and its associated M satellite, which together support the killer phenotype in *S. cerevisiae* (9). This incompatibility between RNAi and the ecologically beneficial L-A/M killer virus system has been proposed as one selective force favoring RNAi loss in fungi (9). Similarly, RNAi loss has been documented in the phytopathogenic fungus *Ustilago maydis* and the human fungal pathogens *Cryptococcus deuterogattii* and *Malassezia* (12–14). In *Candida albicans*, the widely used reference strain SC5314 is RNAi-deficient due to a missense mutation in Argonaute, although most clinical isolates are predicted to be RNAi-proficient (15).

*Cryptococcus neoformans* is a major human fungal pathogen, responsible for approximately 112,000 deaths annually and accounting for ∼19% of all HIV/AIDS-related mortality worldwide (16). Owing to its substantial disease burden and limited treatment options, the World Health Organization (WHO) has designated *C. neoformans* as a critical-priority fungal pathogen (17). Beyond its clinical importance, *C. neoformans* has also emerged as a powerful model for studying fungal genetics and epigenetics because of its genetic tractability and well-developed molecular toolkits (18–21). Notably, the *C. neoformans* reference strain (i.e., H99) retains a broad repertoire of epigenetic mechanisms, including H3K9 and H3K27 methylation, DNA methylation, and RNAi (5, 6, 22–25). Many of these pathways have been lost during the evolution of classical model yeasts, with all four absent from *S. cerevisiae* and only a subset retained in *Schizosaccharomyces pombe* (26).

Despite the conservation of a functional RNAi pathway in the reference strain, natural loss of RNAi has been observed in a subset of isolates from the global *C. neoformans* population. In our previous study, we identified ∼2% of isolates in a global strain diversity collection of clinical and environmental strains as RNAi-deficient due to naturally occurring mutations in RNAi components (8). Some RNAi-deficient isolates have accumulated extremely high transposon burdens and exhibit a hypermutator phenotype in response to antifungal drug selection, whereas others maintain or reduce their transposon copy number and remain non-hypermutators under altered environmental conditions (8, 10). These findings raise a broader question: what additional biological consequences arise from natural RNAi loss in this major human fungal pathogen?

Similar to animals and plants, fungi can also be infected by viruses, collectively known as mycoviruses (27, 28). Mycoviruses are widespread across the fungal kingdom. For example, a large-scale survey found that approximately 21% of tested early-diverging fungal species harbored RNA mycoviruses (29). Although many mycovirus infections are cryptic with no obvious effects on their fungal hosts, others can substantially alter fungal physiology, fitness, and virulence (28). In the chestnut blight fungus *Cryphonectria parasitica*, mycovirus infection, exemplified by Cryphonectria hypovirus 1 (CHV1), causes hypovirulence and has been successfully exploited for biological control (30, 31). Although mycoviruses have been most extensively studied in model yeasts and plant pathogenic fungi, emerging evidence indicates that they are also widespread in human fungal pathogens and can modulate fitness and virulence in these organisms (32). For example, curing *Aspergillus fumigatus* polymycovirus-1M (AfuPmV-1M) has been shown to significantly reduce fungal fitness and virulence (33). Additionally, mycoviruses from the human fungal pathogens *Talaromyces marneffei* and *Malassezia sympodialis* have been linked to altered host immune responses and transcriptional rewiring (34–36). Surprisingly, mycoviruses have yet to be identified in some of the most well-studied human fungal pathogens, including *C. neoformans* and *C. albicans* (28), underscoring the possibility that biologically important mycoviruses remain widely overlooked among major human fungal pathogens.

RNAi constitutes a major antiviral defense against mycoviruses in fungi (37). The RNAi machinery can recognize and process viral double-stranded RNA into virus-derived small interfering RNAs (vsiRNAs), which guide sequence-specific degradation of viral RNA (37). Consistent with this antiviral role, the natural absence of RNAi has been associated with mycovirus presence (9, 34), while experimental disruption of RNAi can increase viral accumulation or susceptibility to mycovirus infection (38, 39). Together, these observations show that RNAi attenuates viral replication and can even cure the host of these viruses. However, RNAi is not the only antiviral defense in fungi. In *S. cerevisiae*, which lacks RNAi, the cytoplasmic Ski complex and the 5′-3′ exonuclease Xrn1 restrict replication of the L-A totivirus and its M satellite, and disruption of these antiviral pathways produces the classic “superkiller” phenotype (40–42). Whether naturally occurring RNAi loss promotes mycovirus persistence in human fungal pathogens, and how RNAi and other antiviral pathways cooperate to restrict these infections, remain poorly understood.

Here, we investigated the biological consequences of natural RNAi loss in *C. neoformans*. Total-RNA sequencing of naturally RNAi-deficient isolates led to the discovery of a novel dsRNA mycovirus, which we named Cryptococcus neoformans totivirus 1 (CnTV1), and expanded screening identified CnTV1 in additional RNAi-deficient isolates. We found that CnTV1-infected isolates share an ancestral mitochondrial DNA insertion disrupting *AGO1* and used genetic restoration approaches to directly test the relationship between RNAi deficiency and viral persistence. We further examined RNAi-independent antiviral pathways and developed a cytoplasmic-mixing approach to generate genetically matched CnTV1-infected and virus-cured strains, enabling the biological consequences of viral infection to be assessed. Together, this work establishes a direct mechanistic link between naturally occurring RNAi deficiency and persistent mycovirus infection and provides a framework for dissecting the biological consequences of mycovirus infection in *C. neoformans*.

## Results

### Identification and phylogenetic characterization of a novel mycovirus in RNAi-deficient *C. neoformans* isolates

We recently discovered that RNAi loss occurs in ∼2% of the natural *C. neoformans* population (8). To test the hypothesis that RNAi-deficient isolates serve as candidate hosts for mycoviruses, we performed rRNA-depleted total RNA sequencing (total RNA-seq) to search for mycoviruses in seven known RNAi-deficient natural isolates (Bt65, Bt81, Bt210, LP-RSA2296, NRHc5028, D17-1 and A2-102-5) (8, 10). Sequencing reads from total RNA-seq were mapped to corresponding genome assemblies. The Bt65, Bt81, Bt210, and LP-RSA2296 genome assemblies were obtained from our previous studies (8, 10), while telomere-to-telomere (T2T) assemblies for NRHc5028, D17-1, and A2-102-5 were generated in this study. Unmapped reads were assembled *de novo* to identify potential viral genomes. By this approach, we recovered a ∼4.6 kb viral contig from the clinical isolate NRHc5028 (SI Appendix, Fig. S1). This viral genome contains two predicted open reading frames encoding a capsid protein (Gag) and an RNA-dependent RNA polymerase (RdRp, also known as Pol) (Fig. 1A). We further confirmed the presence of this novel mycovirus by two independent approaches: 1) total RNA extraction followed by dsRNA enrichment and electrophoretic analysis with agarose gels; 2) RT-PCR with primers specific to the viral Gag and Pol sequences. Positive signals were detected only in NRHc5028 and not in the other RNAi-deficient isolates or the reference strain H99 (Fig. 1B).

**Figure 1.**
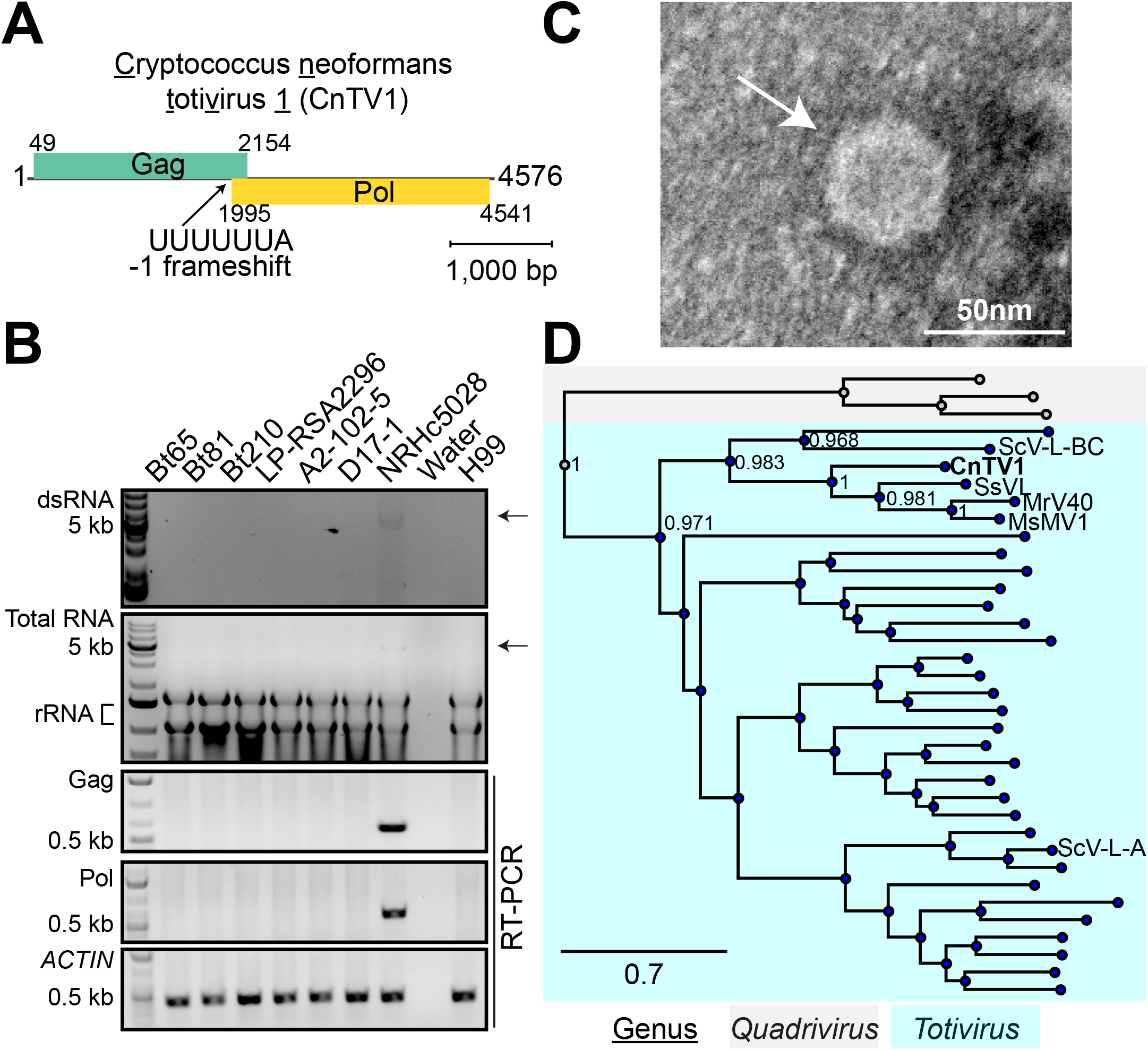
Identification of CnTV1 in an RNAi-deficient *C. neoformans* clinical isolate. (A) Schematic representation of the CnTV1 genome organization. The 4,576 bp dsRNA genome contains two UTRs and two partially overlapping ORFs encoding the capsid protein (Gag) and RNA-dependent RNA polymerase (Pol). A putative heptanucleotide slippery sequence (UUUUUUA) is located within the overlapping region of the two ORFs. (B) Detection of CnTV1 in purified dsRNA, total RNA, and by RT-PCR amplification from the indicated strains. Arrows indicate the presence of the dsRNA band of CnTV1 in strain NRHc5028. The reference strain H99 was included as a negative control for virus detection. The *ACTIN* gene served as a positive control for RT-PCR. (C) TEM image of a CnTV1 viral particle (white arrow). (D) RdRp-based phylogenetic analysis placed CnTV1 within the genus *Totivirus*. The totiviruses L-BC and L-A from *S. cerevisiae* are included for reference. Numbers at nodes indicate Shimodaira–Hasegawa branch support values, and the unit of the scale bar is amino acid substitutions per site. Only virus names and support values for the CnTV1-containing clade are shown. The complete phylogeny is presented in SI Appendix, Fig. S4.

To further characterize the virus, viral particles were purified from NRHc5028 and visualized by transmission electron microscopy (TEM), revealing isometric virus-like particles ∼45 nm in diameter (Fig. 1C). RNA extracted from purified virions revealed substantial enrichment of the ∼4.6 kb RNA species relative to total cellular RNA preparations (SI Appendix, Fig. S2). Enzymatic treatment of the purified viral nucleic acid demonstrated resistance to DNase I and S1 nuclease but sensitivity to RNase III, consistent with a double-stranded RNA genome (SI Appendix, Fig. S3). 5′ and 3′ rapid amplification of cDNA ends (5’/3’ RACE) extended the assembled contig and resolved the terminal sequences, yielding a complete 4,576 bp viral genome containing two predicted open reading frames encoding Gag and Pol, flanked by 48 bp and 35 bp 5′ and 3′ untranslated regions (UTRs), respectively (Fig. 1A).

Phylogenetic analysis of the predicted RdRp placed the virus within the genus *Totivirus* (order *Ghabrivirales*, family *Orthototiviridae)*, where it formed a distinct lineage, most closely related to Scheffersomyces segobiensis virus L (SsVL), Malassezia sympodialis mycovirus (MsMV), and Malassezia restricta virus (MrV), and more distantly related to the well-characterized Saccharomyces cerevisiae virus L-BC (Fig. 1D and SI Appendix, Fig. S4). Consistent with its placement within the genus *Totivirus*, the viral genome contains two partially overlapping ORFs encoding Gag and Pol, separated by a heptanucleotide slippery sequence (UUUUUUA) and a predicted downstream pseudoknot structure (Fig. 1A and SI Appendix, Fig. S5). These features suggest expression of a Gag-Pol fusion protein through programmed −1 ribosomal frameshifting, a well-established translational strategy among totiviruses (43). RdRp amino acid identities between the novel *C. neoformans* mycovirus and its closest relatives ranged from 45% to 48%, well below the 70% species demarcation threshold established for the order *Ghabrivirales* (44). We therefore designated this virus <u>C</u>ryptococcus <u>n</u>eoformans toti<u>v</u>irus <u>1</u> (CnTV1), representing a novel species within the genus *Totivirus*. To our knowledge, CnTV1 is the first mycovirus identified in *C. neoformans*.

### A NUMT insertion in *AGO1* underlies RNAi deficiency in CnTV1-infected isolates

To further assess CnTV1 prevalence, we utilized a publicly available rRNA-depleted total RNA-seq dataset (45) comprising 31 *C. neoformans* isolates representing three lineages (VNI, VNBI, and VNBII). The dataset included RNA-seq libraries generated under one or more growth conditions, including YPD, artificial cerebrospinal fluid (CSF), human CSF, and rabbit CSF. Importantly, the dataset also included the virus-free strain H99 and the CnTV1-infected strain NRHc5028. We screened all the isolates for evidence of CnTV1 by mapping RNA-seq reads to the full-length viral genome.

CnTV1-mapping reads were detected in 10 of the 31 isolates (∼32%), including the known CnTV1-infected strain NRHc5028. Among the screened isolates, NRHc5010 and NRHc5028 were the only strains that consistently exhibited high-confidence evidence of CnTV1 infection, with >80% genome coverage and a mean viral genome sequencing depth exceeding 10x across multiple independent RNA-seq libraries (SI Appendix, Fig. S6). dsRNA enrichment and RT-PCR analysis of NRHc5010 grown on YPD medium confirmed the presence of CnTV1 (Fig. 2A and SI Appendix, Fig. S7). We then assembled the complete viral genome from the NRHc5010 RNA-seq dataset and compared it with the previously identified CnTV1 genome from NRHc5028. The two viral genomes shared approximately 97% nucleotide identity, with 98% and 99% amino acid identity in Gag and Pol, respectively, indicating that they represent closely related variants of the same virus (SI Appendix, Fig. S5C and SI Dataset 1). Accordingly, we designated the virus identified in NRHc5028 as CnTV1A, and the virus identified in NRHc5010 as CnTV1B. Notably, NRHc5010 exhibited exceptionally high viral abundance, particularly in human CSF, where the mean depth of coverage across the CnTV1 genome reached 128,792x, suggesting robust viral replication under host-mimicking conditions. Eight additional isolates each contained detectable CnTV1-mapped reads in only one RNA-seq library, with mean coverage depths ranging from 11x to 1,575x (SI Appendix, Fig. S6). Five were classified as high-confidence based on a mean CnTV1 coverage depth of ≥50x (NRHc5027, NRHc5030, PMHc1033, PMHc1063, and PMHc1065), whereas three were classified as medium-confidence based on a mean depth of <50x after passing the preliminary threshold of ≥100 mapped reads (PMHc1051, PMHc1040, and NRHc5045). However, total RNA gel electrophoresis and RT-PCR analysis of YPD-grown cultures of all five additional high-confidence isolates and one medium-confidence isolate (PMHc1040) failed to detect CnTV1 (SI Appendix, Fig. S8). We therefore considered these isolates CnTV1 false positives and excluded them from subsequent analyses.

**Figure 2.**
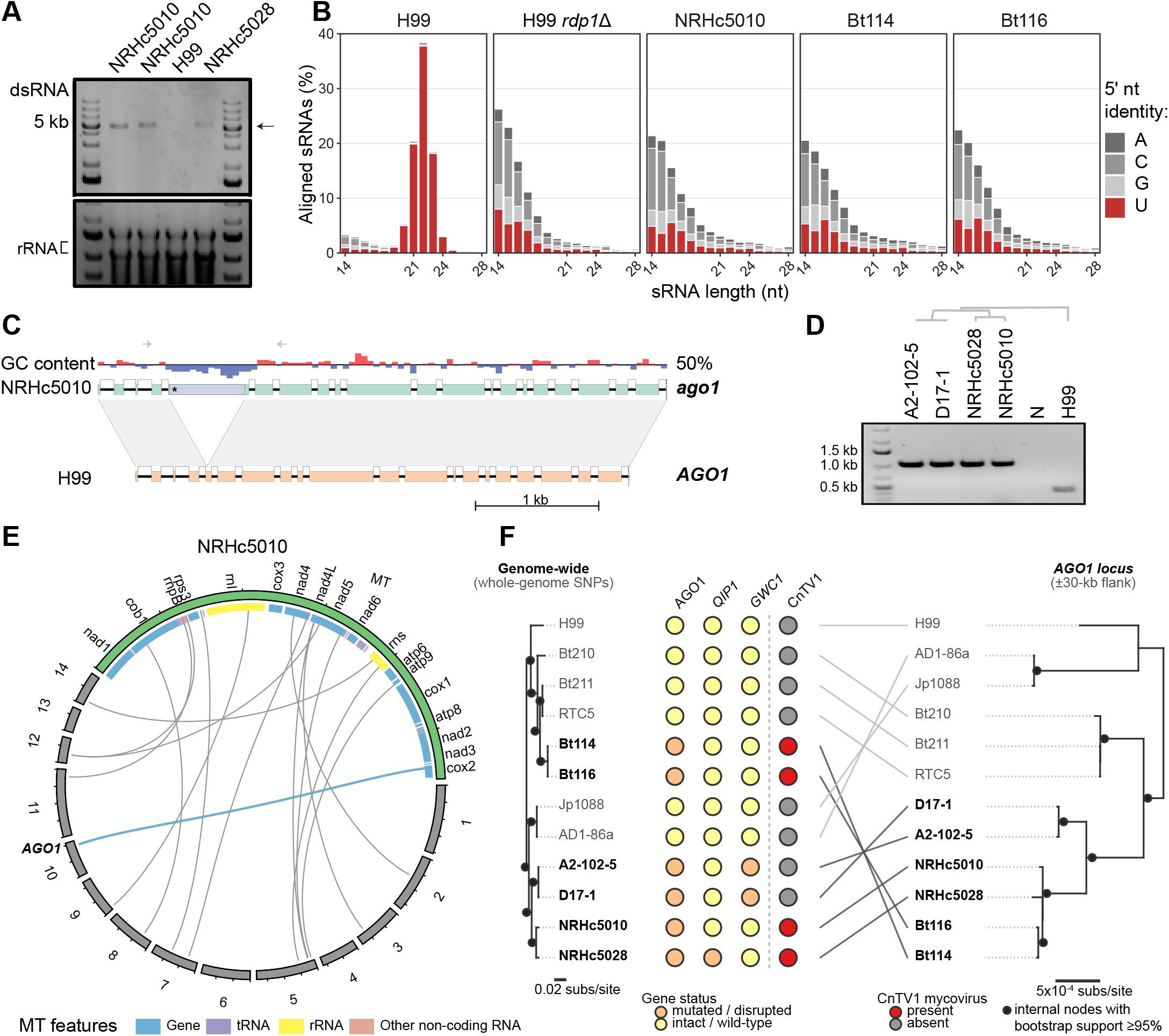
Newly identified CnTV1-infected isolates contain a novel mitochondrial DNA insertion disrupting *AGO1*. (A) Detection of CnTV1 in the clinical isolate NRHc5010. Enriched dsRNA from purified total RNA of NRHc5010 was separated by agarose gel electrophoresis, with rRNA shown as a loading control. Two independent biological replicates of NRHc5010 were analyzed. The arrow indicates the dsRNA band of CnTV1. RNA extracted from H99 and NRHc5028 served as negative and positive controls, respectively, for CnTV1 detection. (B) Length and 5′-nucleotide composition of genome-aligned sRNAs from wild-type H99, the RNAi-deficient control H99 *rdp1*Δ, NRHc5010, Bt114, and Bt116. Bars show the percentage of aligned sRNAs at each length, colored according to the 5′ nucleotide. (C) Comparison of the *AGO1* locus in NRHc5010 and H99. Gene models and local GC content are shown, with homologous regions connected by gray shading. The asterisk marks the predicted premature stop codon caused by the mitochondrial DNA insertion that disrupts *AGO1* in NRHc5010. Arrows indicate the primer positions used for PCR validation of the insertion. (D) PCR validation of the *AGO1* insertion in the indicated isolates. The insertion-containing allele produces an approximately 1 kb amplicon, whereas the wild-type H99 allele produces an approximately 0.5 kb amplicon. N, no-template control (water). The dendrogram above the gel indicates the genome-wide relationships among the tested insertion-positive isolates. (E) Circos representation of sequence similarity between the NRHc5010 nuclear genome and mitochondrial genome. Nuclear chromosomes are shown in gray and the mitochondrial genome in green, with mitochondrial features indicated by the inner colored track. The blue link connects the mitochondrial source region and the insertion within *AGO1* on chromosome 10, while the gray links represent additional nuclear regions with homologs located in the mitochondrial genome. Tick marks represent 0.5 Mb on the nuclear chromosomes and 5 kb on the mitochondrial genome to improve visualization across different genome scales. (F) Comparison of the phylogenetic relationships among 12 isolates based on SNPs across the whole genome (left) and within a 60,027 bp region spanning the *AGO1* insertion site (right). Lines connect corresponding isolates between the two phylogenies. Circles indicate the genotypes at the *AGO1*, *QIP1*, and *GWC1* loci, and the presence/absence of the CnTV1 mycovirus. Isolates carrying the *AGO1* insertion are highlighted in bold. Scale bars indicate substitutions per site.

CnTV1-infected strains NRHc5010 and NRHc5028 are both clinical isolates recovered from patients with HIV/AIDS in Botswana (46). Interestingly, the two isolates are clustered closely together in a previous population genomic study of *C. neoformans* (47). Although NRHc5028 had previously been identified as an RNAi-deficient isolate (8), it remained unknown whether NRHc5010 also lacked RNAi. To determine the RNAi status of NRHc5010, we performed small RNA sequencing (sRNA-seq). Unlike the RNAi-proficient strain H99, NRHc5010 lacked the canonical hallmarks of RNAi-mediated silencing, including enrichment of 21–24-nt siRNAs and the characteristic 5′ uridine bias (Fig. 2B). These findings establish CnTV1-infected NRHc5010 as a novel RNAi-deficient isolate. This prompted us to investigate the genetic basis underlying its RNAi deficiency. Notably, NRHc5010 lacked the previously reported *QIP1* mutation responsible for RNAi deficiency in NRHc5028 and the *GWC1* mutation from the closely related strains D17-1 and A2-102-5 (8). Furthermore, our bioinformatic pipeline, designed to identify SNP- and small indel-based loss-of-function mutations in known RNAi components, failed to detect any candidate mutation in NRHc5010 (8). We therefore performed long-read Nanopore sequencing of NRHc5010 to identify structural variants that may underlie its RNAi-deficient phenotype. This effort yielded a near-complete, chromosome-level genome assembly comprising all 14 chromosomes, with telomeric repeat sequences absent from several chromosome ends. Whole-genome synteny analysis further revealed a large reciprocal translocation between Chr 1 and Chr 3 in NRHc5010 that was not observed in NRHc5028, A2-102-5, or D17-1, supporting genomic divergence among these closely related RNAi-deficient isolates (SI Appendix, Fig. S9). Interestingly, comparison of the NRHc5010 genome assembly with the H99 reference genome identified a 609 bp, low-GC-containing DNA insertion within the coding region of the canonical RNAi gene *AGO1* (Fig. 2C). This insertion is predicted to introduce a premature stop codon, resulting in a truncated Ago1 protein. PCR validation confirmed the presence of this insertion in NRHc5010, as well as in the closely related RNAi-deficient isolates NRHc5028, A2-102-5, and D17-1 (Fig. 2D). Unexpectedly, BLAST analysis revealed that the inserted sequence is 100% identical to a mitochondrial DNA fragment immediately adjacent to the *COX2* locus, demonstrating that the insertion represents a nuclear mitochondrial DNA segment (NUMT) integration event (Fig. 2E). The 5′ insertion junction carries a 4 bp microhomology (CATT), whereas the 3′ junction carries only a 2 bp microhomology (TT). Both are shared between the ancestral *AGO1* recipient locus and the mitochondrial-derived insert, consistent with microhomology-mediated end joining (MMEJ) repair (48, 49), with more extensive microhomology at the 5′ break. Together, these analyses identified a shared mitochondrial-derived insertion disrupting *AGO1* in multiple closely related RNAi-deficient isolates, suggesting that *AGO1* disruption by NUMT integration arose from a single ancestral integration event before the emergence of additional loss-of-function mutations in other RNAi components, including *QIP1* and *GWC1* (8).

### A rare ancestral *AGO1* NUMT insertion is shared across divergent VNI isolates

The presence of an identical NUMT insertion in *AGO1* in four closely related isolates prompted us to investigate whether this insertion is restricted to this small RNAi-deficient group or is more broadly distributed across the *C. neoformans* population. We first screened a collection of 387 globally distributed isolates spanning all four major lineages (VNI, VNII, VNBI, and VNBII) for which Illumina whole-genome sequencing data are available (47). We mapped reads to a targeted reference and scored them based on whether they span the insertion junction and whether read pairs bridge *AGO1* to the mitochondrial donor sequence (see Extended Methods). We detected the insertion in six isolates, all belonging to VNI: the four previously validated isolates (NRHc5010, NRHc5028, A2-102-5, and D17-1) and two additional isolates (Bt114 and Bt116). This insertion was not identified in any non-VNI isolates (0/16 VNII, 0/122 VNBI, 0/64 VNBII), resulting in a significant association with the VNI lineage within this isolate collection (6/185 VNI versus 0/202 non-VNI; Fisher’s exact test, *p* value = 0.011). Even within VNI, however, the carrier frequency was low (∼3%), indicating that the insertion is present in a small fraction of the lineage rather than being a general VNI feature (Fig. 2F). To test this directly, we screened a second, independent collection of 677 VNI isolates (50). The NUMT insertion in *AGO1* was absent from all 677, confirming that the insertion is present in a small subset of VNI rather than most of the lineage.

Like the original four *AGO1*-NUMT carriers, Bt114 and Bt116 are clinical isolates recovered from HIV/AIDS patients in Botswana. PCR validation and amplicon sequencing further confirmed that the same NUMT insertion in *AGO1* is present in isolates Bt114 and Bt116. Total RNA and RT-PCR analyses additionally confirmed the presence of CnTV1 in both isolates (SI Appendix, Fig. S10). The partial Gag and Pol sequences recovered by RT-PCR from Bt114 and Bt116 share 100% sequence identity with each other and are >96% identical to those recovered from NRHc5010 and NRHc5028 (SI Dataset 1). Additionally, sRNA-seq analysis demonstrated a marked depletion of 21–24-nt siRNAs and a depletion of 5′ uridine bias, confirming that Bt114 and Bt116 are RNAi-deficient (Fig. 2B).

The identical NUMT insertions at the same position in the *AGO1* gene across all six strains strongly suggest that they descend from a single ancestral integration event. However, a phylogeny based on genome-wide polymorphisms showed that they belong to two distinct clades, one containing Bt114 and Bt116, which belong to the VNIc lineage, and the other containing the other four strains (NRHc5010, NRHc5028, A2-102-5 and D17-1), which belong to the VNIa lineage (47, 50). Notably, each clade also contains strains that do not contain the NUMT insertion and have wild-type *AGO1* alleles (e.g., RTC5, Bt211, Jp1088, and AD1-86a) (Fig. 2F, left). Thus, there are two possibilities. First, the identical NUMT insertion event occurred repeatedly and independently in strains belonging to different clades. In this case, the sequences adjacent to the insertion should show phylogenetic relationships generally consistent with the genome-wide phylogeny. The second possibility is that the NUMT insertion in all six strains descended from a single event. In this case, sequences adjacent to the NUMT insertion would be expected to be more similar among the NUMT-carrying strains than to non-carrying strains, and their phylogeny would disagree with the one based on genome-wide polymorphisms, reflecting shared local ancestry around the insertion.

To test this, we constructed a phylogeny of the six NUMT-carrying strains, together with a collection of their closely related strains based on whole-genome polymorphisms, using only the ∼60 kb genomic region that encompasses the *AGO1* locus (∼30 kb on either side) to balance phylogenetic resolution with local ancestry (Fig. 2F, right). Interestingly, this phylogeny supported a very different strain relationship. Specifically, the six strains carrying the NUMT insertion now clustered together and formed a well-supported monophyletic clade. Additionally, Bt114 and Bt116 were deeply embedded in this clade and were more closely related to strains NRHc5010 and NRHc5028. Furthermore, within this 60 kb region, the six strains showed very high sequence similarity (mean pairwise nucleotide p-distance = 0.054%). This divergence was substantially lower than that observed between the NUMT-carrying strains and closely related non-NUMT strains identified from the whole-genome phylogeny (mean pairwise nucleotide p-distance = 0.195%) and was also lower than that between the NUMT-carrying strains and more distantly related wild-type reference strains, such as H99 and Bt210 (mean pairwise nucleotide p-distance = 0.190%). Our results thus support a model in which the NUMT insertion in these six strains originated from a single ancestral event and the surrounding genomic region retained shared local ancestry despite the genome-wide divergence of these strains. Together with the additional, lineage-specific lesions identified in other RNAi components (Fig. 2F, left), including *QIP1* and *GWC1* (8), these findings suggest that the ancestral *AGO1* disruption may have preceded the accumulation of additional loss-of-function mutations in other RNAi components.

### RNAi controls CnTV1 during sexual reproduction and vegetative growth

Because all CnTV1-positive isolates carry one or more mutations affecting the RNAi pathway, we hypothesized that loss of RNAi permits viral persistence, and that the rescue of RNAi would restrict CnTV1. To test this hypothesis, we crossed the CnTV1A-positive isolate NRHc5028 (*MAT*α *ago1 qip1*) with the CnTV1-negative laboratory strain KN99**a** (*MAT***a** *AGO1 QIP1*) and recovered 29 F1 progeny (Fig. 3A). We selected representative progeny carrying all four combinations of the *AGO1* and *QIP1* alleles and analyzed their small RNA profiles by sRNA-seq. By mapping to the H99 nuclear genome, progeny carrying a mutation in either *AGO1* or *QIP1* lacked the characteristic 21–24-nt siRNA population and 5′ uridine bias, whereas progeny carrying wild-type alleles at both loci displayed a functional RNAi profile (Fig. 3B). These results demonstrate that *AGO1* and *QIP1* are each necessary for RNAi activity in *C. neoformans*. Interestingly, only two of the 29 progeny retained CnTV1, as confirmed by total RNA gel electrophoresis and CnTV1-specific RT-PCR. Both progeny were RNAi-deficient, carrying a mutation in either *AGO1* (JHG40) or *QIP1* (JHG35) (Fig. 3A). This pattern was inconsistent with passive, unrestricted cytoplasmic inheritance and suggested that RNAi may restrict CnTV1 transmission during sexual reproduction. We therefore hypothesized that RNAi activity in the dikaryotic zygote limits CnTV1 inheritance by the sexual progeny. To test this hypothesis, we crossed NRHc5028 with either KN99**a** *ago1*Δ or KN99**a** *qip1*Δ, generating dikaryotic zygotes lacking a functional copy of *AGO1* or *QIP1*, respectively. Strikingly, 100% of the recovered progeny inherited CnTV1 in both crosses (13/13 and 16/16, respectively). We next repaired *QIP1* in NRHc5028 and crossed the resulting strain with KN99**a** *qip1*Δ. Although each parental strain was individually RNAi-deficient, the dikaryotic zygote received functional copies of both *AGO1* and *QIP1* through complementation between the two parental nuclei and was therefore expected to be RNAi-proficient. Under this condition, 0% (0/15) of the recovered progeny inherited CnTV1 (Fig. 3A). Together, these results indicate that the reconstitution of a functional RNAi pathway in the dikaryotic zygote restricts CnTV1 transmission during sexual reproduction, preventing detectable inheritance of CnTV1 in the progeny.

**Figure 3.**
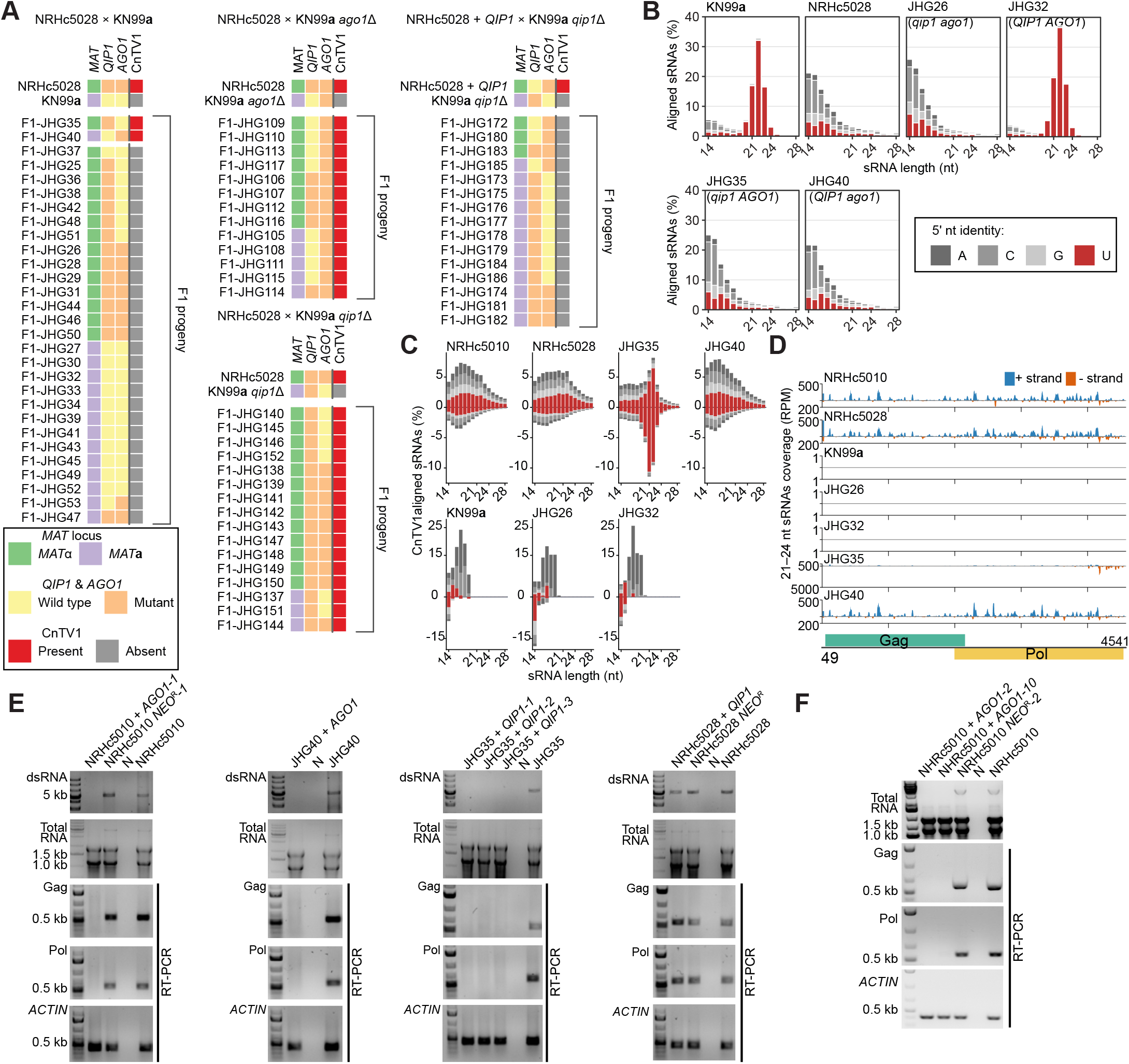
RNAi restricts CnTV1 during sexual reproduction and vegetative growth. (A) Segregation of *MAT*, *QIP1*, *AGO1*, and CnTV1 among F1 progeny from the indicated crosses. (B) Size distribution and 5’ nucleotide identity of nuclear genome-aligned small RNAs from parental strains and representative F1 progeny. (C) Size and strand distributions of CnTV1-derived sRNAs in the indicated strains. Bars show the percentage of CnTV1-aligned sRNAs at each length, with reads mapping to the plus and minus strands plotted above and below the x-axis, respectively. Strains shown in the top panels are CnTV1-positive, whereas those in the bottom panels are CnTV1-negative. Bars are colored according to the 5’ nucleotide identity using the same color scheme as in (B). (D) Genome-wide distribution of 21–24-nt CnTV1-derived sRNAs in the indicated strains. Plus- and minus-strand reads are plotted above and below the baseline, respectively, as reads per million (RPM). Note the different y-axis scales used for CnTV1-positive and CnTV1-negative strains. The positions of the predicted Gag and Pol coding regions are shown at the bottom. (E) CRISPR-mediated repair of *AGO1* or *QIP1* in CnTV1-positive strains using H99-derived repair donors. Viral RNA was assessed by dsRNA gel electrophoresis and targeted RT-PCR; *ACTIN* served as a control. (F) CRISPR-mediated repair of *AGO1* in NRHc5010 using an NRHc5010-derived repair donor eliminated CnTV1, as assessed by total RNA gel electrophoresis and targeted RT-PCR.

Additionally, we mapped the sRNA-seq reads to a modified H99 genome carrying the CnTV1 sequence as an added contig to characterize CnTV1-derived viral sRNAs (vsRNAs) across different genetic backgrounds. Among the CnTV1-positive strains, only JHG35, which retained a functional *AGO1* allele and a *qip1* mutation, accumulated a distinct 21–22-nt vsRNA peak (Fig. 3C). This size-specific enrichment was absent from NRHc5010, NRHc5028, and JHG40, all of which carried *ago1* mutations, despite abundant CnTV1-derived sRNA reads in these strains overall (Fig. 3C). CnTV1-negative strains (JHG26, JHG32 and KN99**a**) showed only background-level mapping to the viral genome (∼200 reads per sample, compared with >150,000 reads in CnTV1-positive strains) and no enrichment of 21–24-nt species (Fig. 3C). Further examination of the 21–24-nt vsRNA pool revealed that JHG35 displayed an approximately balanced distribution between the plus and minus strands across the CnTV1 genome, together with a prominent antisense hotspot near the 3′ end of Pol (Fig. 3C and 3D). In contrast, the other CnTV1-positive strains showed a strongly plus-strand-biased 21–24-nt population that was broadly distributed across the viral genome. No comparable 21–24-nt vsRNA population was detected in the CnTV1-negative strains (Fig. 3D). Together, these findings associate functional *AGO1* with the accumulation of a discrete 21–22-nt CnTV1-derived sRNA population, consistent with antiviral siRNAs. However, CnTV1 persisted in the absence of functional *QIP1* despite these siRNAs, indicating that siRNA accumulation alone is insufficient for viral elimination.

Beyond its role during sexual reproduction, we next tested whether a functional RNAi pathway could eliminate CnTV1 from infected strains. We restored RNAi function with CRISPR-mediated allele exchange in multiple independent CnTV1-positive genetic backgrounds with donor DNA from the H99 background. In NRHc5010, replacement of the NUMT-disrupted *AGO1* allele with a functional *AGO1* allele resulted in complete loss of detectable CnTV1, as confirmed by dsRNA enrichment gel electrophoresis and RT-PCR (Fig. 3E). By contrast, CnTV1 was retained in an NRHc5010 control strain carrying only the *NEO* resistance cassette, demonstrating that the CRISPR transformation and selection procedures had no effect on mycovirus persistence (Fig. 3E). Likewise, restoration of *AGO1* in the CnTV1-positive F1 progeny JHG40 (*ago1 QIP1*) also resulted in complete loss of detectable CnTV1 (Fig. 3E). Similarly, repair of *QIP1* in another CnTV1-positive F1 progeny, JHG35 (*AGO1 qip1*), eliminated detectable CnTV1 in all three independently generated repair strains (Fig. 3E). In contrast, repair of *QIP1* in NRHc5028 (*ago1 qip1*), in which *AGO1* remained nonfunctional, failed to eliminate CnTV1 (Fig. 3E). Introduction of the *NEO* resistance cassette alone into the same genetic background likewise had no effect on viral persistence (Fig. 3E). Considering the additional *AGO1* sequence variation that distinguishes H99 from NRHc5010 beyond the NUMT insertion, we generated an NRHc5010-derived repair donor in which only the NUMT insertion was removed, retaining the remaining NRHc5010-specific *AGO1* sequence. Two independently repaired strains generated with this donor also lost CnTV1, whereas control transformants carrying the *NEO* marker alone remained CnTV1-positive (Fig. 3F). Together, these results demonstrate that the NUMT insertion disrupts Ago1-dependent RNAi and that restoration of functional RNAi is sufficient to eliminate CnTV1.

### RNAi-independent mechanisms suppress CnTV1

In addition to RNAi, we investigated whether RNAi-independent mechanisms restrict CnTV1 proliferation in *C. neoformans*. We focused on four genes with established roles in yeast antiviral defense, particularly against L-A and related dsRNA viruses: *NUC1*, encoding a mitochondrial endonuclease; *REX2*, encoding an RNA exonuclease; *SKI2*, encoding an RNA helicase and core component of the cytoplasmic SKI RNA decay complex; and *XRN1*, encoding a 5′–3′ exoribonuclease (40–42, 51). To assess whether any of these genes have antiviral activity against CnTV1, we generated deletion mutants in the CnTV1-infected isolates NRHc5028 (CnTV1A) and NRHc5010 (CnTV1B) (SI Appendix, Fig. S11) and quantified CnTV1 RNA with RT-qPCR. Unlike what is observed for L-A totivirus in *S. cerevisiae* (40), deleting *NUC1* and/or *REX2* had no effect on viral RNA levels for either CnTV1A or CnTV1B. In contrast, deleting *SKI2* or *XRN1* significantly elevated CnTV1 RNA levels in both strains, suggesting that these genes contribute to limiting CnTV1 accumulation in *C. neoformans* (Fig. 4A-B). Additionally, the *nuc1*Δ *ski2*Δ double mutant showed a similar increase in viral RNA levels to the *ski2*Δ single mutant, indicating no synthetic effect by deleting both genes, in contrast to that observed in *S. cerevisiae* (40) (Fig. 4A-B).

**Figure 4.**
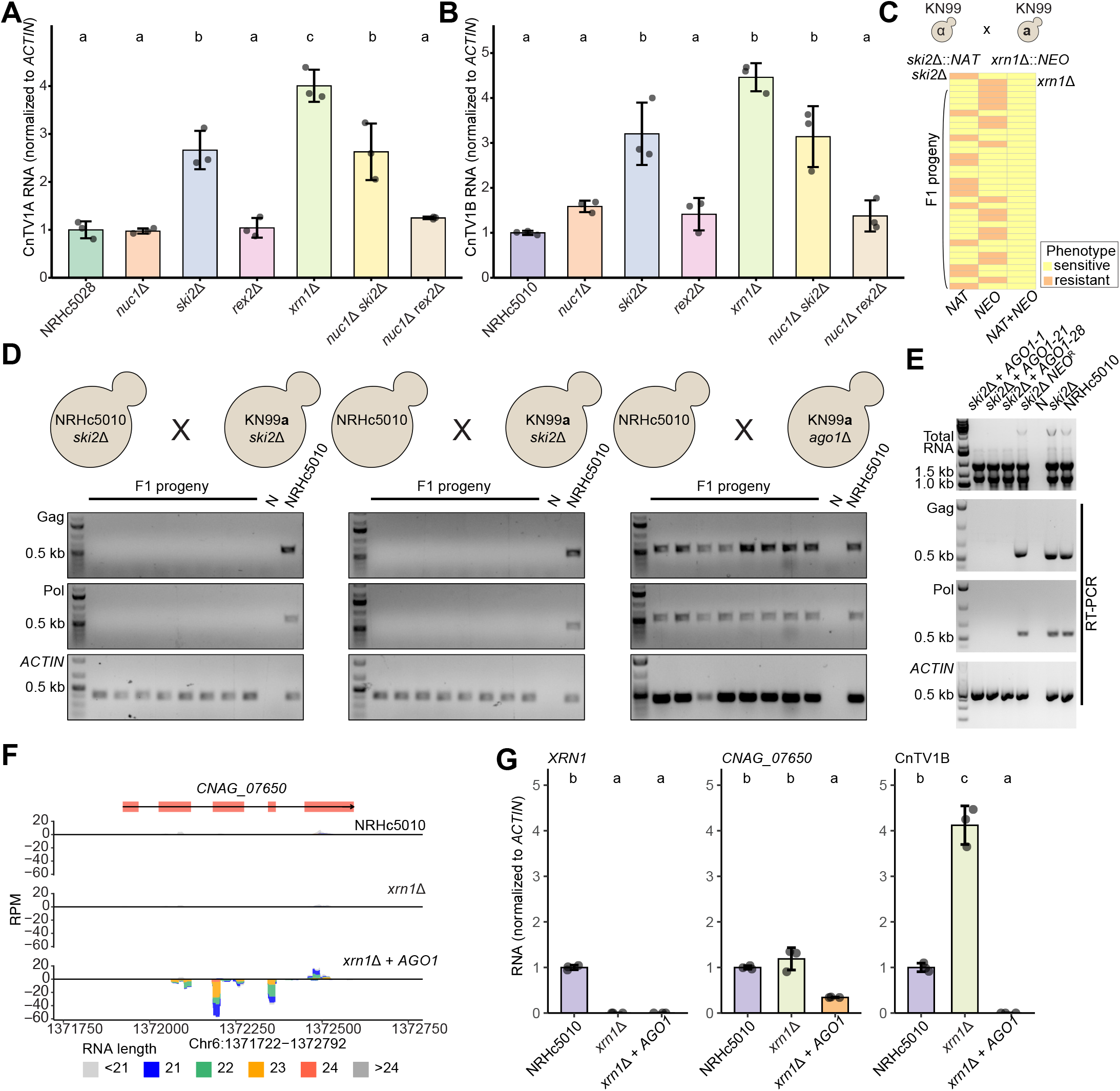
RNAi-dependent and RNAi-independent mechanisms restrict CnTV1. (A and B) CnTV1 RNA levels in the indicated mutant backgrounds derived from NRHc5028 carrying CnTV1A (A) and NRHc5010 carrying CnTV1B (B), quantified by RT-qPCR and normalized to *ACTIN*. Points represent biological replicates, and bars show mean ± SD. Different letters indicate statistically significant differences between groups. (C) Segregation of *SKI2* and *XRN1* alleles among 33 F1 progeny from a cross between KN99α *ski2*Δ::*NAT* and KN99**a** *xrn1*Δ::*NEO*. Resistance to NAT, NEO, and combined NAT+NEO selection is shown for each progeny. (D) RNAi in the dikaryotic zygote is sufficient to eliminate CnTV1 despite the absence of *SKI2*. Viral RNA was assessed in F1 progeny derived from the indicated crosses by RT-PCR; *ACTIN* served as a control. (E) CRISPR-mediated repair of *AGO1* in NRHc5010 *ski2*Δ. Viral RNA was assessed by total RNA gel electrophoresis and targeted RT-PCR; *ACTIN* served as a control. (F) sRNA-seq profiles across the predicted noncoding RNA *CNAG_07650* in NRHc5010, NRHc5010 *xrn1*Δ, and NRHc5010 *xrn1*Δ + *AGO1*. Reads per million (RPM) are shown across the indicated genomic region. (G) RT-qPCR quantification of *XRN1*, *CNAG_07650*, and CnTV1B RNA in NRHc5010, NRHc5010 *xrn1*Δ, and NRHc5010 *xrn1*Δ + *AGO1*. Transcript levels were normalized to *ACTIN*. Points represent biological replicates, and bars show mean ± SD. Different letters indicate statistically significant differences between groups. For A, B, and G, statistical analyses were performed on ΔCt values using one-way ANOVA followed by Tukey’s honestly significant difference (HSD) post hoc test.

To assess whether combined disruption of *SKI2* and *XRN1* would further increase CnTV1 proliferation, we attempted to generate *ski2*Δ *xrn1*Δ double mutants in NRHc5028 and NRHc5010 but were unsuccessful (SI Appendix, Fig. S12). Interestingly, these genes have been shown to exhibit synthetic lethality in *S. cerevisiae* independent of viral infection (52). To test whether this interaction is conserved in *C. neoformans*, we crossed KN99α *ski2*Δ::*NAT* with KN99**a** *xrn1*Δ::*NEO* (both are CnTV1-free) and phenotyped the resulting F1 progeny for NAT and NEO resistance. We recovered no progeny resistant to both NAT and NEO, indicating that *SKI2* and *XRN1* are synthetically lethal in *C. neoformans*, independently of CnTV1 infection (chi-square test, *p* value = 0.0011) (Fig. 4C).

Given that loss of *SKI2* or *XRN1* increased CnTV1 levels in RNAi-deficient backgrounds, we hypothesized that disruption of either pathway might be sufficient to permit CnTV1 persistence even in the presence of functional RNAi. To test this hypothesis,

we crossed NRHc5010 (*MAT*α *ago1*) *ski2*Δ with KN99**a** *ski2*Δ and analyzed CnTV1 segregation in F1 progeny by RT-PCR. None of the progeny inherited CnTV1 (Fig. 4D, left). We obtained the same results by crossing NRHc5010 (*MAT*α *ago1*) with KN99**a** *ski2*Δ (Fig. 4D, middle). These observations are consistent with RNAi-mediated viral restriction in the dikaryotic zygote, despite the absence of *SKI2* in one or both parental strains. In contrast, when we crossed NRHc5010 (*MAT*α *ago1*) with KN99**a** *ago1*Δ, all progeny inherited CnTV1, consistent with efficient viral transmission through an RNAi-deficient dikaryotic zygote (Fig. 4D, right). Similarly, restoration of a functional *AGO1* allele in NRHc5010 *ski2*Δ was sufficient to eliminate CnTV1, as confirmed by total RNA gel electrophoresis and RT-PCR (Fig. 4E).

We also attempted similar mating-based segregation assays with NRHc5010 *xrn1*Δ; however, severe mating defects precluded recovery of sufficient progeny for analysis, consistent with the critical role of *XRN1* in *C. neoformans* sexual reproduction reported previously (53). Despite this, we restored a functional *AGO1* allele in NRHc5010 *xrn1*Δ via CRISPR-mediated allele exchange. Consistent with restoration of RNAi activity in NRHc5010 *xrn1*Δ + *AGO1*, we observed enrichment of 21–24-nt sRNAs mapping to the predicted noncoding RNA *CNAG_07650* by sRNA-seq, accompanied by reduced *CNAG_07650* transcript abundance (Fig. 4F-G). Furthermore, RT-qPCR quantification of CnTV1 RNA showed that restoration of *AGO1* in NRHc5010 *xrn1*Δ eliminated detectable viral RNA (Fig. 4G). Together, these results do not support the hypothesis that the loss of *SKI2* or *XRN1* is sufficient to permit CnTV1 persistence in the presence of functional RNAi. Instead, RNAi is the dominant antiviral mechanism and is sufficient to eliminate CnTV1.

### CnTV1 is stably maintained in NRHc5010 during prolonged *in vitro* growth and *in vivo* infection

To assess the biological consequences of CnTV1 infection, we generated five independent CnTV1B-cured derivatives of NRHc5010, primarily through cycloheximide treatment and serial passage (see Extended Methods), with viral clearance confirmed by RT-PCR and total RNA gel electrophoresis (SI Appendix, Fig. S13). Whole-genome sequencing followed by mapping to the H99 reference and subtraction of variants present in the parental strain NRHc5010 identified an average of ∼131 candidate variants per cured strain (range, 108–179). Approximately 78% were shared among multiple derivatives, including 31 present in all five strains. Read-level inspection showed that 97% of shared SNVs were already present in parental NRHc5010 at high allele frequencies (median Variant Allele Frequency ∼0.95), indicating that they largely represented pre-existing background variation. On average, ∼29 candidate private variants remained per strain, nearly all with predicted low impact. Only one private high-impact mutation was identified across the five CnTV1B-cured derivatives (SI Dataset 2). These residual genomic differences indicated that the chemically cured derivatives were not strictly isogenic to the parental strain and could confound downstream phenotypic analyses.

Despite this, we evaluated the virulence of the parental CnTV1B-infected NRHc5010 strain, two independently derived chemically cured strains (TCD3 and JHG225), and one genetically cured derivative (JHG84, *AGO1*) in a murine intranasal infection model. Survival was monitored for 60 days. All strains in the NRHc5010 background showed significantly reduced virulence relative to H99 (H99 vs NRHc5010, Gehan-Breslow-Wilcoxon test, *p* value <0.0001), whereas survival differences among NRHc5010 and its derivatives were not significant (SI Appendix, Fig. S14).

We also examined whether the CnTV1 infection status remained stable during prolonged growth *in vitro* and whether CnTV1 persisted during infection *in vivo*. Following five consecutive *in vitro* passages (on solid YPD medium at 2-day intervals), viral Gag and Pol RNA remained readily detectable in NRHc5010, whereas the CnTV1B-cured derivatives remained virus-free, indicating that both the infected and cured states were stable during vegetative growth (SI Appendix, Fig. S15A). CnTV1B was also retained following prolonged murine infection (60 days), with viral RNA detected in fungal isolates recovered from both murine brain and lung tissues (SI Appendix, Fig. S15B).

### BLOSSOM enables generation of isogenic strain pairs differing only in mycovirus infection status

Given the low virulence of the NRHc5010 background and the presence of genetic differences between NRHc5010 and the chemically cured CnTV1B-free isolates, we sought to generate isogenic *C. neoformans* strain pairs in a highly virulent genetic background that differ only in their CnTV1 infection status. To this end, we developed a cytoplasmic mixing method that allows the virus to be introduced or removed from a specific genetic background, which we term **<u>Bl</u>**astosp**<u>o</u>**re-mediated **<u>s</u>**preading and **<u>s</u>**ilencing **<u>o</u>**f **<u>m</u>**ycovirus (BLOSSOM) (Fig. 5A). To introduce CnTV1, matings were set up between two RNAi-deficient strains of opposite mating types, of which one harbors the mycovirus (donor) and the other does not (recipient). Blastospores (mitotically produced yeast cells that bud from dikaryotic hyphae prior to nuclear fusion and meiosis) (5) emerging along the dikaryotic hyphae were subsequently isolated and screened for cells carrying the recipient genome and the presence of the mycovirus (Fig. 5A, left). To remove the mycovirus, matings were set up between one strain that is RNAi-deficient and virus-infected (target) and another that is RNAi-proficient and virus-free. Blastospores were then isolated and screened for those containing the target genome, and in which the virus has been removed due to the RNAi-proficient nature of the zygote and dikaryotic hyphae (Fig. 5A, right).

**Figure 5.**
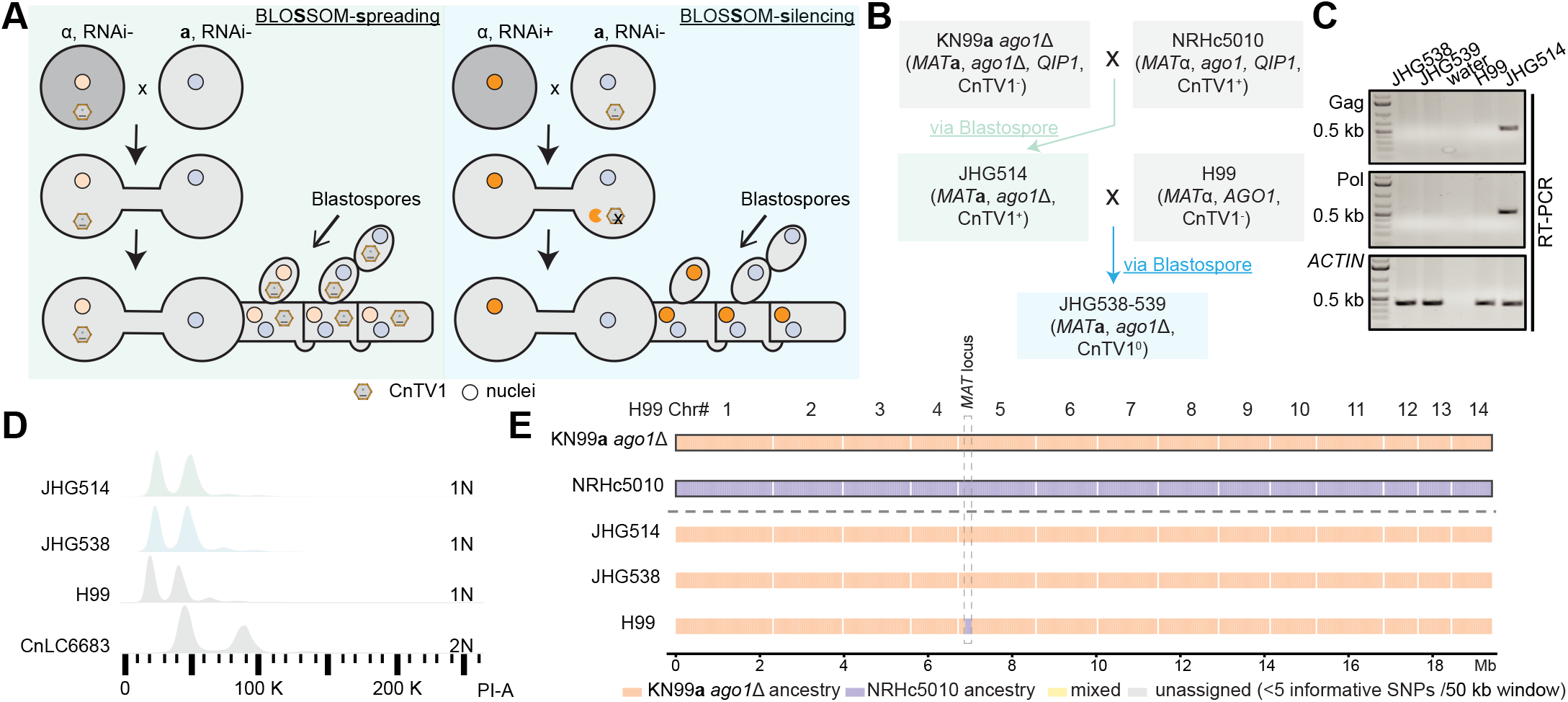
BLOSSOM enables generation of isogenic CnTV1-infected and -cured strains. (A) Schematic of the Blastospore-mediated Spreading and Silencing of Mycovirus (BLOSSOM) method. For BLOSSOM-spreading (left), an RNAi-deficient, CnTV1-infected donor is crossed with an RNAi-deficient, virus-free recipient, allowing CnTV1 to spread through the shared cytoplasm and into blastospores carrying the recipient nucleus. For BLOSSOM-silencing (right), an RNAi-deficient, CnTV1-infected target strain is crossed with an RNAi-proficient, virus-free strain, allowing RNAi-mediated elimination of CnTV1 from blastospores carrying the target nucleus. Gray and white cell shading indicates the mitochondrial genotype inherited from the corresponding parental strains and illustrates uniparental mitochondrial inheritance from the *MAT***a** parent. (B) Strain genealogy showing the generation of the CnTV1B-infected strain JHG514 and CnTV1B-cured strains JHG538 and JHG539 using BLOSSOM. Blue and green arrows indicate progeny isolated via blastospore dissection. Genotypes, mating types, and CnTV1 infection status are indicated. (C) CnTV1 infection status of the indicated strains, as assessed by RT-PCR using primers targeting the viral Gag and Pol regions. *ACTIN* served as a control. (D) Ploidy analysis of JHG514 and JHG538 by FACS. H99 and CnLC6683 served as haploid (1N) and diploid (2N) controls, respectively. (E) Genome-wide ancestry analysis based on informative polymorphisms distinguishing the KN99**a** *ago1*Δ and NRHc5010 parental backgrounds. Colors indicate KN99**a** *ago1*Δ ancestry, NRHc5010 ancestry, mixed ancestry, and regions with insufficient informative SNPs for ancestry assignment. The *MAT* locus is indicated by dashed vertical lines.

Using this method, we first crossed KN99**a** *ago1*Δ (recipient) with NRHc5010 (*MAT*α *ago1*; donor), generating strain JHG514, which carries the KN99**a** *ago1*Δ genetic background and is CnTV1B-infected (CnTV1B^+^). We next crossed JHG514 with H99 (*MAT*α) and recovered two blastospores, JHG538 and JHG539, that carry the KN99**a** *ago1*Δ genetic background and are CnTV1B-cured (CnTV1B^0^) (Fig. 5B). We also used BLOSSOM to generate additional pairs of CnTV1A-infected (CnTV1A+) and -cured (CnTV1A) strains (SI Appendix, Fig. S16). We validated the infection status of the generated strains by RT-PCR (Fig. 5C and SI Appendix, Fig. S16). Fluorescence-activated cell sorting (FACS) analysis of dissected blastospores showed that the vast majority remained haploid, although diploid profiles were occasionally observed (Fig. 5D and SI Appendix, Fig. S16). One such exception occurred when we attempted to remove CnTV1A from JHG499 (KN99**a** *ago1*Δ CnTV1A^+^) by crossing it with H99. The initial cured product, JHG530, was diploid (*MAT*α/**a**, CnTV1A). Because JHG530 carried both mating type alleles, we induced selfing and recovered the haploid *MAT***a** derivative JHG597 (*MAT***a** CnTV1A) (SI Appendix, Fig. S16).

Eventually, we successfully obtained two pairs of isogenic strains in the KN99**a** *ago1*Δ background: JHG499 (CnTV1A^+^)/JHG597 (CnTV1A^0^) and JHG514 (CnTV1B^+^)/JHG538 (CnTV1B^0^). These infected/cured strain pairs were used for subsequent analyses. Whole-genome sequencing read-depth analysis revealed no evidence of *de novo* aneuploidy in these strains (SI Appendix, Fig. S17). Genome-wide SNP analysis based on informative polymorphisms between the parents further showed that the BLOSSOM-derived strains were predominantly of KN99**a** *ago1*Δ ancestry, with no evidence of substantial NRHc5010- or NRHc5028-derived genomic regions (Fig. 5E and SI Appendix, Fig. S18). In contrast, the same analysis readily detected recombinant ancestry blocks in F1 progeny (JHG35 and JHG40) derived from meiotic basidiospores (SI Appendix, Fig. S18). Finally, serial *in vitro* passage showed that the infection status of these BLOSSOM-derived strains was stable (SI Appendix, Fig. S19), and RT-qPCR showed that CnTV1A and CnTV1B RNA levels in the BLOSSOM-derived infected strains were comparable to those of their respective parental infected strains, NRHc5028 and NRHc5010 (SI Appendix, Fig. S20). Together, these results establish BLOSSOM as a reliable approach for generating isogenic mycovirus-infected and - cured strain pairs for assessing virus-associated phenotypes.

### CnTV1 infection modestly impacts the fungal transcriptome and virulence in a murine inhalation model of cryptococcosis

Having established isogenic CnTV1-infected and -cured strain pairs in the KN99**a** *ago1*Δ background using BLOSSOM, we next investigated the biological consequences of CnTV1 infection. We first examined three major *in vitro* virulence-associated phenotypes: thermotolerance, polysaccharide capsule production, and melanin production. All strains showed similar thermotolerance profiles across the temperatures tested, including at the host-relevant temperature of 37°C both with and without 5% CO2 (Fig. 6A). Similarly, we observed no obvious differences in cell morphology, cell size, capsule production, or melanin production between the infected and cured strains (Fig. 6B-C and SI Appendix, Fig. S21).

**Figure 6.**
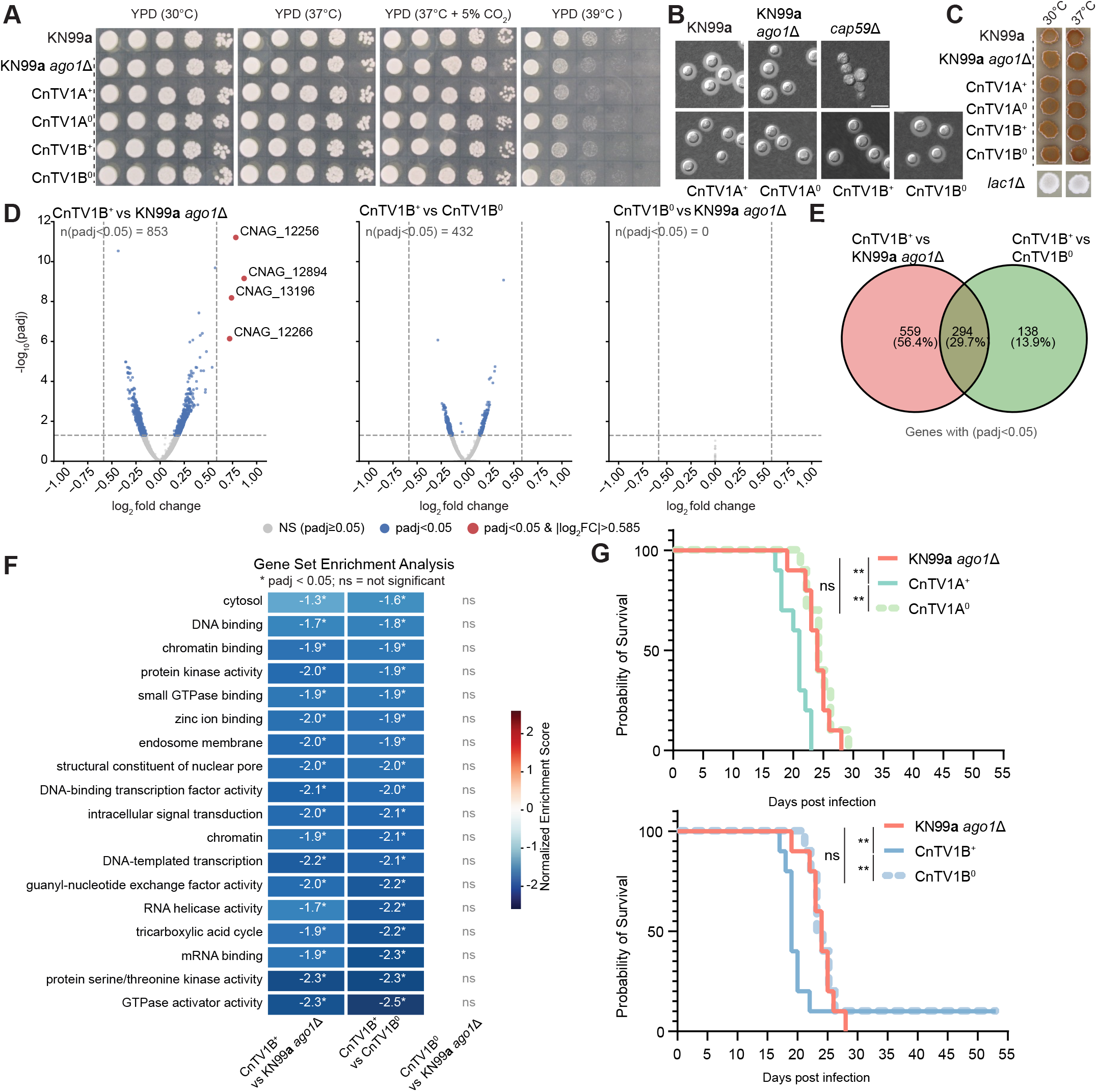
CnTV1 infection modestly impacts the fungal transcriptome and virulence in a murine inhalation model of cryptococcosis. (A) Growth of the indicated strains under different temperature and CO_2_ conditions. Serial dilutions were spotted onto YPD medium and incubated at 30°C, 37°C, 37°C with 5% CO_2_, or 39°C for 2 days before imaging. (B) CnTV1-infected and -cured strains exhibit similar cell morphologies, size, and polysaccharide capsule production. Strains were grown in liquid RPMI at 37°C for 3 days before imaging. Cells were negatively stained with India ink to visualize the capsules. Strain *cap59*Δ served as an acapsular control strain. Scale bar represents 10 µm. (C) CnTV1-infected and - cured strains exhibit similar melanin production. Saturated cultures of each strain were spotted onto Niger seed medium. Plates were incubated at the indicated temperatures for 1 week before imaging. Strain *lac1*Δ served as a melanin-deficient control strain. (D) Volcano plots showing host transcriptional differences among CnTV1B^+^, CnTV1B^0^, and the progenitor KN99**a** *ago1*Δ strain following exclusion of CnTV1-derived reads. Dashed horizontal and vertical lines indicate adjusted *p* value = 0.05 and |log_2_ fold change| = 0.585 (1.5-fold), respectively. Blue points indicate genes with an adjusted *p* value < 0.05, and red points indicate genes with an adjusted *p* value < 0.05 and |log_2_ fold change| > 0.585. The four genes exceeding the 1.5-fold threshold are indicated. (E) Overlap of host genes with an adjusted *p* value < 0.05 between the CnTV1B^+^ versus KN99**a** *ago1*Δ and CnTV1B^+^ versus CnTV1B^0^ comparisons. (F) GSEA of host transcriptional changes associated with CnTV1B infection. Heatmap shows normalized enrichment scores (NES) for functional categories significantly enriched (adjusted *p* value < 0.05) in comparisons of CnTV1B^+^ with either KN99**a** *ago1*Δ or CnTV1B^0^. The CnTV1B^0^ versus KN99**a** *ago1*Δ comparison is shown to assess reversibility following viral clearance. Asterisks indicate an adjusted *p* value < 0.05; ns, not significant. (G) Survival of A/J mice infected with the indicated CnTV1A^+^ and CnTV1A strains (top) or CnTV1B^+^ and CnTV1B strains (bottom), with KN99**a** *ago1*Δ included for comparison. All groups were tested concurrently in the same experiment, and the same KN99**a** *ago1*Δ control group is displayed in both plots to facilitate visual comparison. Statistical significance was assessed using the Gehan–Breslow–Wilcoxon test. Median survival times (95% CI) were 25 (22–28), 24 (19–25), 21 (17–22), 24 (21–26), 19 (17–20), and 23.5 (21–25) days for KN99**a**, KN99**a** *ago1*Δ, CnTV1A^+^, CnTV1A^0^, CnTV1B^+^, and CnTV1B^0^, respectively.

Given the lack of obvious differences in these *in vitro* phenotypes, we next asked whether CnTV1 infection induces more subtle changes at the transcriptional level. We therefore performed rRNA-depleted total RNA-seq on RNA extracted from overnight YPD liquid cultures of the BLOSSOM-generated CnTV1B-infected strain JHG514 (CnTV1B^+^), the corresponding cured strain JHG538 (CnTV1B^0^), and the virus-free parental KN99**a** *ago1*Δ strain. One KN99**a** *ago1*Δ replicate was identified as a technical outlier based on PCA and was excluded from subsequent analyses (SI Appendix, Fig. S22). When CnTV1-derived reads were included, PCA clearly separated CnTV1B^+^ from CnTV1B^0^ and KN99**a** *ago1*Δ along PC1, which explained 71% of the total variance (SI Appendix, Fig. S22). In contrast, excluding CnTV1-derived reads substantially diminished this separation, with PC1 explaining 35% of the variance and no clear segregation by viral infection status (SI Appendix, Fig. S22).

Consistent with the limited separation of the host transcriptomes, differential gene expression analysis revealed that although many host genes showed statistically significant changes, the magnitude of these changes was generally small. Specifically, 853 and 432 host genes had adjusted *p* value < 0.05 in the CnTV1B^+^ versus KN99**a** *ago1*Δ and CnTV1B^+^ versus CnTV1B^0^ comparisons, respectively, but none exhibited greater than two-fold changes in expression. Only four predicted non-coding RNAs (*CNAG_12256*, *CNAG_12894*, *CNAG_13196*, and *CNAG_12266*) reached 1.5-fold changes. In contrast, no genes had an adjusted *p* value < 0.05 when comparing CnTV1B and parental KN99**a** *ago1*Δ, further supporting the isogenic nature of the BLOSSOM-generated strains (Fig. 6D). Notably, 294 genes with an adjusted *p* value < 0.05 were shared between the CnTV1B^+^ versus KN99**a** *ago1*Δ and CnTV1B^+^ versus CnTV1B^0^ comparisons (Fig. 6E), supporting a reproducible but low-magnitude transcriptional response associated with CnTV1 infection. GO enrichment analysis of these overlapping genes did not identify significantly enriched functional categories after multiple-testing correction (SI Dataset 3).

Because these transcriptional changes were widespread but individually small in magnitude, we next asked whether they reflected coordinated changes in specific biological processes. We therefore performed gene set enrichment analysis (GSEA) using the ranked transcriptomic data. Comparison of CnTV1B^+^ with CnTV1B^0^ revealed coordinated downregulation of multiple functional categories, including those related to transcription, RNA helicase activity, protein kinase and GTPase-associated activities, mRNA binding, and the tricarboxylic acid cycle (Fig. 6F). Importantly, 18 functional categories showed significant enrichment in the same direction in both the CnTV1B^+^ versus KN99**a** *ago1*Δ and CnTV1B^+^ versus CnTV1B^0^ comparisons (adjusted *p* value < 0.05), whereas significant enrichment was not observed in the comparison between CnTV1B^0^ and KN99**a** *ago1*Δ (adjusted *p* value ≥ 0.05). These findings indicate that these pathway-level changes were consistently associated with CnTV1 infection and largely reversed following CnTV1 elimination (Fig. 6F). Together, these analyses indicate that in liquid YPD at 30°C, CnTV1 infection is associated with a broad but low-magnitude transcriptional response characterized by coordinated downregulation of genes involved in multiple cellular processes, much of which is reversed following viral clearance.

Finally, we compared the virulence of the infected and cured strains in a murine inhalation model of cryptococcosis. We first compared the parental KN99**a** *ago1*Δ strain with wild-type KN99**a** and observed no significant difference in murine survival (Gehan-Breslow-Wilcoxon test, *p* value = 0.1147), consistent with previous reports that RNAi loss does not affect virulence in *C. neoformans* (5) (SI Appendix, Fig. S23). The CnTV1A^+^ strain exhibited a modest increase in virulence relative to both the parental KN99**a** *ago1*Δ strain (*p* value = 0.0022) and its isogenic cured counterpart, CnTV1A^0^ (*p* value = 0.0012) (Fig. 6G, top), while no difference was observed between KN99**a** *ago1*Δ and CnTV1A^0^ (*p* value = 0.8197). Similarly, we observed a mildly increased virulence for the independently generated CnTV1B^+^ strain (Fig. 6G, bottom, *p* value = 0.0059 for CnTV1B^+^ versus KN99**a** *ago1*Δ, *p* value = 0.0016 for CnTV1B^+^ versus CnTV1B^0^, *p* value = 0.9394 for CnTV1B^0^ vs KN99**a** *ago1*Δ). Compared with their corresponding virus-cured controls, CnTV1A^+^ and CnTV1B^+^ infections reduced median mouse survival by 3 and 4.5 days, respectively (Fig. 6G). Together, these observations suggest that CnTV1 curing is associated with a modest decrease in virulence in a murine inhalation model. However, the mechanisms underlying this mycoviral effect remain to be determined.

## Discussion

In this study, we investigated the biological consequences of RNAi loss in *C. neoformans* and hypothesized that RNAi-deficient isolates are more permissive to mycovirus infection. Building on our previous identification of seven naturally occurring RNAi-deficient isolates (8, 10), we discovered a novel dsRNA totivirus CnTV1 in the clinical isolate NRHc5028 and subsequently identified CnTV1 in three additional RNAi-deficient isolates, NRHc5010, Bt114, and Bt116. During preparation of this manuscript, an independent preprint also identified CnTV1 sequences in NRHc5010, NRHc5028, and PMHc1063 through analysis of publicly available RNA-seq datasets and confirmed their presence by RT-PCR (54). However, we were unable to detect CnTV1 in PMHc1063 by either total RNA gel electrophoresis or RT-PCR (SI Appendix, Fig. S8). We further found that NRHc5010 is RNAi-deficient due to a NUMT insertion disrupting *AGO1*. The identical NUMT insertion in *AGO1* was also present in NRHc5028, Bt114, and Bt116, all of which were infected with CnTV1, as well as in the CnTV1-negative isolates A2-102-5 and D17-1. Although the NRHc5010/NRHc5028 and Bt114/Bt116 groups are divergent in genome-wide SNP-based analyses, all NUMT-carrying isolates cluster together based on *AGO1*-flanking sequences, which supports a single ancestral NUMT insertion event. Importantly, the presence of the same NUMT insertion in both CnTV1-positive and CnTV1-negative isolates indicates that *AGO1* disruption does not determine viral infection status but instead creates a background permissive for CnTV1 persistence after viral exposure. The identification of NRHc5010, Bt114 and Bt116 as additional naturally occurring RNAi-deficient isolates brings the total identified in our studies to ten, further supporting our previous suggestion that the frequency of RNAi loss in the global *C. neoformans* population may be underestimated (8).

Our identification of a NUMT insertion that disrupts RNAi and permits mycovirus persistence in *C. neoformans* highlights NUMT insertions as a potentially underappreciated source of functional genetic variation. In this case, mitochondrial DNA transferred into the nuclear genome disrupted RNAi – a conserved genome-defense pathway, thereby altering host-virus interactions. The impact of mitochondrial DNA damaging the nuclear genome is already appreciated in humans, where NUMT insertions occur spontaneously in the germline, generating NUMT diversity within human populations (55). NUMT insertions have also been associated with a variety of human diseases (56). Our findings extend the potential functional consequences of NUMT insertion to this major human fungal pathogen and suggest that they may contribute to naturally occurring genetic and phenotypic diversity. Defining the prevalence, genomic distribution, and functional consequences of NUMT insertions across *C. neoformans* populations will therefore be an important area for future investigation.

In the *Cryptococcus* pathogenic species complex, RNAi is dispensable for a variety of stress responses and for virulence (5, 8, 10, 14, 23). Nevertheless, RNAi plays a key role in silencing repetitive sequences, including transposable elements (TEs) and repetitive transgenes, during both vegetative growth and sexual reproduction (5, 8, 10, 23, 57–59). We previously showed that RNAi loss can result in TE proliferation and hypermutation driven by uncontrolled transposition (8, 10). The present study expands the biological consequences of naturally occurring RNAi loss beyond genome defense against endogenous repetitive elements to include altered interactions with mycoviruses. Whether mycovirus infection and transposon activity interact in RNAi-deficient backgrounds remains an intriguing question for future investigation.

A relationship between RNAi loss and mycovirus persistence has also been observed in other fungi and is perhaps best illustrated by the killer system of the model budding yeast *S. cerevisiae*. The yeast killer system consists of the dsRNA mycovirus L-A and its satellite dsRNA M, which encodes a toxin that enables *S. cerevisiae* to kill and outcompete neighboring cells while remaining immune to the toxin. Restoring RNAi in *S. cerevisiae* eliminates the killer system and sensitizes cells to killing by neighboring toxin-producing cells. Consistent with incompatibility between RNAi and the killer system, killer viruses are absent from RNAi-proficient species, whereas all known killer-harboring species are RNAi-deficient. Thus, RNAi loss may have provided an evolutionary advantage by permitting maintenance of the killer system in *S. cerevisiae* and related species (9, 11). Our results establish a related association between natural RNAi deficiency and mycovirus persistence in *C. neoformans*. Whether CnTV1 provides a sufficient advantage that would lead to selection for RNAi-deficient genotypes in natural populations remains unknown.

Our results suggest that RNAi is the dominant antiviral mechanism operating in *C. neoformans*. To date, CnTV1 has been identified only in RNAi-deficient isolates, and restoration of RNAi is sufficient to prevent viral inheritance during sexual reproduction and eliminate the virus during vegetative growth. Although CnTV1 persists only in RNAi-deficient isolates, these isolates retain partial pathway function: we detected 21–24-nt CnTV1-derived vsRNAs in virus-positive isolates, and an F1 progeny carrying functional *AGO1* but lacking *QIP1* showed a clear 21–22-nt peak with canonical 5′-uridine enrichment in its vsRNA population, indicating that residual RNAi machinery directly targets CnTV1. Because full RNAi restoration rapidly eliminates the virus, an inducible RNAi system could help capture and dissect this antiviral siRNA response. However, RNAi is not the only pathway that restricts CnTV1. We identified Ski2, a cytoplasmic RNA helicase and core component of the Ski RNA-degradation complex, and Xrn1, a 5’-to-3’ exoribonuclease, as additional factors that contribute to limiting CnTV1 accumulation in *C. neoformans*. Both proteins have established antiviral functions in *S. cerevisiae* (40, 51, 60). However, the precise mechanisms by which Ski2 and Xrn1 limit CnTV1 accumulation remain to be determined. Loss of either *SKI2* or *XRN1* increased CnTV1 accumulation in RNAi-deficient backgrounds. *ski2* and *xrn1* mutations are synthetically lethal in *S. cerevisiae* (52), and our inability to recover *C. neoformans ski2*Δ *xrn1*Δ double mutants, including among F1 progeny from a KN99α *ski2*Δ × KN99**a** *xrn1*Δ cross, suggests that this genetic interaction is conserved. Importantly, restoration of RNAi in either the *ski2*Δ or *xrn1*Δ background still eliminated CnTV1, demonstrating that functional RNAi is sufficient for viral clearance even in the absence of either RNA surveillance pathway. Together, these findings support a layered antiviral defense system in which RNAi provides the dominant barrier to CnTV1 persistence, while *SKI2*- and *XRN1*-dependent RNA surveillance provides additional restriction when RNAi is compromised.

Although these findings establish multiple host pathways that restrict CnTV1, an important question is whether persistent CnTV1 infection alters host biology. Assessing the phenotypic consequences of mycovirus infection can be challenging because naturally infected and virus-free isolates often differ extensively in their genetic backgrounds, while chemical or other curing approaches can introduce additional genomic changes. Indeed, our whole-genome analysis of independently chemically cured NRHc5010 derivatives identified background genetic differences that could complicate the direct attribution of phenotypes to CnTV1 infection. To overcome this limitation, we developed BLOSSOM, a cytoplasmic-mixing-based method to generate isogenic CnTV1-infected and virus-cured strains in a laboratory strain background, allowing the biological consequences of CnTV1 infection to be examined while minimizing confounding effects from background genetic variation. We anticipate that BLOSSOM will facilitate studies of additional mycoviruses that may be discovered in *C. neoformans*.

With isogenic CnTV1-infected and CnTV1-cured strains in the KN99**a** *ago1*Δ background, we found that CnTV1 infection was associated with broad but low-magnitude changes in host gene expression, with GSEA revealing an overall tendency toward reduced expression among affected pathways. Together with the absence of major differences in the *in vitro* phenotypes examined, these findings suggest that persistent CnTV1 infection can alter host transcription without broadly disrupting fungal physiology under the conditions tested. Competition assays under diverse environmental conditions may provide a more sensitive approach to detect subtle fitness effects associated with CnTV1 infection. Altered fungal gene expression affecting transcriptional and translational processes has similarly been reported during mycoviral infection in *Malassezia* (34, 36). Despite these relatively subtle effects *in vitro*, loss of CnTV1 infection was associated with a modest decrease in virulence in a murine inhalation model of cryptococcosis. By contrast, the naturally infected isolate NRHc5010 was less virulent than the H99 reference strain, and NRHc5010 showed no significant virulence difference from its chemically cured derivatives. However, these comparisons are complicated by background genomic variation among the chemically cured strains, highlighting the value of the isogenic BLOSSOM-derived strains for more directly assessing the contribution of CnTV1 to virulence. Similar observations of mycovirus-associated increases in virulence have been reported in other human fungal pathogens. Infection of *Aspergillus fumigatus* by the dsRNA virus A. fumigatus polymycovirus-1M (AfuPmV-1M) confers a fitness advantage during murine pulmonary infection, whereas virus-cured strains exhibit decreased fitness and virulence (33). Similarly, infection of *Talaromyces marneffei* with the dsRNA virus Talaromyces marneffei partitivirus 1 (TmPV1) is associated with increased virulence during murine infection relative to isogenic virus-free isolates (35).

The mechanism underlying the increased virulence associated with CnTV1 infection remains unknown. Because we did not observe obvious differences in the *in vitro* virulence-associated phenotypes examined, including thermotolerance, capsule production, and melanization, the virulence effect may involve fungal traits not captured by these assays or altered interactions with the mammalian host. For example, viral RNA from the Malassezia restricta totivirus MrV40 is recognized by Toll-like receptor 3 (*TLR3*), inducing an inflammatory response in bone marrow-derived dendritic cells (36). Thus, one possibility is that CnTV1 alters host-pathogen interactions through immune recognition or inflammatory responses, although this hypothesis remains to be tested.

These findings provide a foundation for broader investigation of the *C. neoformans* virome and its antiviral defenses. During the preparation of this manuscript, an independent preprint reported the identification of Cryptococcus gattii totivirus 1 (CgTV1), a dsRNA mycovirus in an environmental isolate of *C. gattii*, and similarly demonstrated that loss of RNAi function enables viral persistence (61). It will be important to determine whether RNAi loss is required for mycovirus persistence in *C. neoformans* or whether some viruses can evade or suppress an intact RNAi pathway. The growing availability of public sequencing datasets will facilitate the discovery of additional mycoviruses and help address this question. In other fungal systems, mycoviruses such as Cryphonectria hypovirus 1 (CHV1) and Aspergillus virus 1816 have been shown to suppress host RNA silencing (35, 62, 63). It will also be important to determine whether residual RNAi components in naturally RNAi-deficient isolates retain antiviral activity, as Dicer proteins in *C. parasitica* can restrict some viruses independent of Argonaute (64). Finally, defining how viral load and viral genotype influence fungal fitness, virulence, and host interactions may reveal new aspects of *C. neoformans* biology and potentially identify exploitable virus–fungus interactions. As a precedent, CHV1 reduces disease severity and has been successfully deployed as a biocontrol agent against chestnut blight (30, 65). Together, *C. neoformans* and CnTV1 provide a genetically tractable system for defining how natural variation in antiviral defense shapes persistent viral infection and fungal pathogenesis.

## Materials and methods

### Ethics statement

All animal experiments in this manuscript were approved by the Duke University Institutional Animal Care and Use Committee (IACUC) (protocol # A098-22-05-25). Animal care and experiments were conducted according to IACUC ethical guidelines.

### Identification and visualization of nuclear mitochondrial DNA transfers (NUMTs)

The mitochondrial genome of NRHc5010 was annotated using MFannot (https://github.com/BFL-lab/MFannot) with genetic code 4 (Mold, Protozoan, and Coelenterate Mitochondrial Code) (66). Synteny of the *AGO1* locus between NRHc5010 and the H99 reference genome was visualized using Easyfig (v2.2.5) (67). Nuclear mitochondrial DNA transfers segments (NUMTs) were identified by aligning the annotated mitochondrial genome to the NRHc5010 nuclear genome assembly using BLASTN (v 2.16.0+). Alignments were filtered to retain hits with a minimum alignment length of 100 bp and ≥90% nucleotide identity. Genome-wide NUMTs were visualized with Circos (v0.69-8) (68), and the mitochondrial genome was scaled 500-fold relative to the nuclear chromosomes to facilitate visualization. NUMTs were displayed as links connecting their mitochondrial source regions to their corresponding nuclear insertion sites.

### BLOSSOM

To introduce and remove CnTV1, crosses between recipient and donor cells (spreading) or between target and wild-type cells (silencing) were set up on MS medium and incubated at room temperature in the dark. After blastospore clusters were formed along mating hyphae, cells from individual clusters were recovered through microdissection as described previously (69). A minimum 14 cells were dissected from most of the clusters, and cells from clusters with high germination rates (>80%) were included in the subsequent genotypic analysis for the nuclear genome and presence/absence of CnTV1. Because mitochondrial inheritance in *C. neoformans* is uniparental from the *MAT***a** parent (70, 71), strains generated using KN99**a** as the *MAT***a** parent were expected to inherit the KN99**a** mitochondrial genome. Mitochondrial genome analysis confirmed no SNP differences between the CnTV1A-infected and CnTV1A-cured strains or between the CnTV1B-infected and CnTV1B-cured strains, consistent with uniparental inheritance of the *MAT***a**-derived mitochondrial genome.

## Data Availability

The *de novo* assemblies of NRHc5010, NRHc5028, A2-102-5, and D17-1 have been deposited with accession number PRJNA1524062 in the NCBI BioProject database. The raw sequence reads for Nanopore sequencing, sRNA-seq, and Illumina whole-genome sequencing have also been deposited under the same BioProject accession number. The CnTV1 genome identified and confirmed with RACE in this work has been submitted to GenBank and are available under GenBank accession number PZ955392. Key analysis scripts used in this study are available at https://github.com/Jhuang90/Huang-et-al.-2026_CnTV1-mycovirus.

## Acknowledgments

We thank all the members of the Heitman Lab for constructive suggestions. We also thank Drs. Guilhem Janbon (Institut Pasteur), Neta Shlezinger (Hebrew University), Mari L. Shinohara, Andy Alspaugh, Edward A. Miao, Jorn Coers, and Alwyn Ecker (Duke University) for helpful discussion. We thank Dr. John Perfect and the Perfect Lab (Duke University) for generously sharing strains. We thank Dr. Ruiyun Zeng (North Carolina State University) for assistance in data visualization, and Dr. Davis Ferreira (Department of Pathology, Center for Electron Microscopy & Nanoscale Technology, Duke University) for assistance with electron microscopy. Electron microscopy was performed using a JEOL JEM-2100Plus transmission electron microscope supported by NIH Shared Instrumentation Grant 1S10OD026776 (PI: Sara Miller). We also thank Dr. Bin Li (Duke Cancer Institute Flow Cytometry Core Facility) and Dr. Devi Swain Lenz (Duke Sequencing and Genomic Technologies Core Facility) for advice and expertise. We also thank Dr. Linqi Wang and Zhenghao Shen (Institute of Microbiology, Chinese Academy of Sciences) for informing us of their independent work on mycoviruses in *Cryptococcus gattii* and coordinating co-submission; no specific findings or unpublished data were discussed in detail or shared prior to co-submission of the two complementary manuscripts.

This study was supported by NIH/NIAID R01 and R21 grants AI039115-28, AI050113-20, AI133654-09, AI170543-05, AI172451-04, and AI199262-01 to J. Heitman; the Canadian Institutes of Health Research (PJT-18592) and Natural Sciences and Engineering Research Council of Canada (RGPIN-201) grants to M.D.M.; and the Canadian Institutes of Health Research (PTJ-496709) to A.B. J. Heitman is codirector and fellow of the Canadian Institute for Advanced Research (CIFAR) program Fungal Kingdom: Threats & Opportunities. A.B. is a CIFAR Global Azrieli Scholar of the CIFAR Fungal Kingdom: Threats & Opportunities program. Computing resources were provided by the Duke Compute Cluster (DCC) and the University of Toronto Cloud Research Lab at The Donnelly, powered by AWS. We thank Dr. Charles Boone (University of Toronto) for connecting J. Heitman with M.D.M. at the Asilomar CIFAR Fungal Kingdom Satellite meeting in March 2024, which resulted in this collaborative effort. We thank the Madhani laboratory (UCSF) and NIH R01 grant AI100272 for the KN99α *ski2*Δ deletion strain.

