## Supplemental Information for "An ancestral mitochondrial DNA insertion disrupts RNAi and enables persistence of a novel mycovirus in *Cryptococcus neoformans*"

**This PDF file includes:**

Supporting text
Extended Materials and Methods
Figures S1 to S23
Tables S1 to S2
Legends for Datasets S1 to S4
SI References

SI Appendix

Supporting text

Extended Materials and Methods

**Fungal strains and growth conditions**

*C. neoformans* strains used or generated in this study are listed in Supplementary Table 1. Strains were stored in 15% glycerol at −80°C for long-term storage. Unless otherwise specified, strains were streaked fresh from glycerol stocks and cultured on yeast extract peptone dextrose (YPD) agar plates at 30°C for up to three days. For liquid culture experiments, strains were inoculated into liquid YPD or Yeast Nitrogen Base (YNB) supplemented with 2% glucose and grown overnight at 30°C in a roller drum rotating at 70 rpm prior to the assay.

**Total RNA-seq analysis for detecting novel mycoviruses in *C. neoformans***

Total RNA-seq data with ribosomal RNA (rRNA) depletion were obtained for seven naturally occurring RNAi-loss strains of *C. neoformans*: Bt65, Bt81, Bt210, LP-RSA2296, A2-102-5, D17-1, and NRHc5028. These strains were previously characterized (1, 2). RNA was extracted from overnight YPD liquid cultures with the mirVana miRNA Isolation Kit (Thermo Fisher Scientific, catalog# AM1561) according to the manufacturer's instructions. Two biological replicates were included for each strain, and libraries were prepared using a total RNA protocol with rRNA reduction, yielding 14 paired-end libraries in total. Samples were then sequenced on an Illumina NovaSeq X Plus platform (2 x 150 bp) at the Duke University Sequencing and Genomic Technologies Core facility. Due to the inadvertent use of non-*Cryptococcus*-specific probes for rRNA depletion at the sequencing facility, approximately 50% of reads were derived from rRNA, although the remaining coverage exceeded 100× for each sample. Adapter and quality trimming of raw reads was performed using Trim Galore (https://github.com/FelixKrueger/TrimGalore) prior to downstream analysis.

Trimmed paired-end reads from each library were assembled independently using rnaSPAdes (v4.0.0) (3) with the –rna mode and 16 threads. Assembled transcripts from all 14 libraries were concatenated into a single reference set. Sequence statistics across the combined file were computed using SeqKit (v2.8.0) (4) . The concatenated transcript set was sorted in descending order of length using SeqKit and clustered at 90% nucleotide identity using USEARCH (v11.0.667) (5) to produce a non-redundant set of representative centroids. Cluster centroids were screened for RNA-dependent RNA polymerase (RdRp) motifs using palm_annot (palmscan2 v2.0) (6), which profiles the conserved polymerase palm domain shared by all positive-sense RNA viruses.

To enrich for non-host transcripts and improve sensitivity for low-abundance viral sequences, we performed a genome depletion pipeline for the sequenced strains. For each strain, a Bowtie2 (v2) (7) index was built from its respective nuclear genome assembly (a telomere-to-telomere or near-complete assembly was available for each of the seven strains). Trimmed paired-end reads were mapped to their respective genome index using Bowtie2 with –local –very-sensitive-local -k 1 settings. Only read pairs in which both mates failed to map (SAM flag 4) were retained. Unmapped reads then underwent a second depletion step using a mitochondrial ribosomal RNA reference to remove residual rRNA contaminants.

To quantify the abundance of CnTV1 across all seven RNAi-loss strains, rRNA- and host genome-depleted trimmed paired-end reads from each library were mapped to the full RACE-confirmed CnTV1 viral genome using Bowtie2 with –local –very-sensitive-local -k 10. Properly paired mapped reads were extracted using SAMtools (8) (excluding unmapped reads and secondary alignments; SAM flag -F 260), and mapping statistics were computed with SeqKit bam. Sorted and indexed BAM files were inspected to confirm the specificity and uniformity of coverage across the viral contig. Per-base read depth across the CnTV1 genome was computed for each library using samtools depth, from which mean coverage depth and breadth of coverage (percentage of genome positions covered at ≥1x) were calculated.

### RNA extraction, dsRNA enrichment, and PCR analyses

For RNA extraction, *C. neoformans* cells were cultured overnight in 8 mL liquid YPD at 30°C in a roller drum rotating at 70 rpm. Cells were harvested by centrifugation, lyophilized overnight, and subjected to total RNA extraction using the mirVana miRNA Isolation Kit (Thermo Fisher Scientific, catalog# AM1561) according to the manufacturer's instructions. RNA concentrations were determined using a Qubit Fluorometer with the Qubit RNA Broad Range Assay Kit (Thermo Fisher Scientific).

dsRNA enrichment was performed as previously described with minor modifications (9). Briefly, DNase I-treated (Thermo Fisher Scientific, catalog# AM1907) total RNA was subjected to selective lithium chloride precipitation (2.8 M LiCl, overnight at −20°C) to remove ssRNA. dsRNA was subsequently recovered from the supernatant by precipitation with 0.1 volume of 3 M NaCl and 2.5 volumes of 100% ethanol, resuspended in nuclease-free water, and analyzed by 1% agarose gel electrophoresis. The double-stranded nature of the enriched RNA species was further assessed by digestion with S1 Nuclease (Thermo Fisher Scientific, catalog# EN0321) and RNase III (Thermo Fisher Scientific, catalog# AM2290) according to the manufacturer's instructions.

First-strand cDNA was synthesized from 500–1,000 ng of total RNA using the Maxima H Minus cDNA Synthesis Master Mix with dsDNase (Thermo Fisher Scientific, catalog# M1681) according to the manufacturer's instructions. The resulting cDNA was diluted 1:5 and used as a template for RT–PCR and quantitative PCR (qPCR). qPCR reactions were performed using Power SYBR Green PCR Master Mix (Thermo Fisher Scientific, catalog# 4367659) on a QuantStudio 3 Real-Time PCR System (Thermo Fisher Scientific). Relative transcript abundance was normalized to *ACTIN* and calculated using the ΔΔCt method. Statistical analyses were performed on ΔCt values using one-way ANOVA followed by Tukey's honestly significant difference (HSD) post hoc test. All qPCR analyses were conducted with three biological replicates and two technical replicates per sample.

#### 5’ and 3’ rapid amplification of cDNA ends (RACE)

RACE assays were performed using RNA isolated from NRHc5028 and the SMARTer® RACE 5′/3′ Kit (Takara, catalog# 634860) according to the manufacturer’s instructions. Because CnTV1 lacks a poly(A) tail, RNA samples were treated with Poly(A) Polymerase (NEB, catalog# M0276S) prior to 3′ RACE. RACE amplicons were cloned into a pRACE vector, and plasmids containing the resulting inserts were sequenced by Plasmidsaurus plasmid sequencing. Terminal sequences were confirmed from multiple independent clones that yielded identical consensus sequences.

#### Viral particle isolation and transmission electron microscopy

Virus particle enrichment and transmission electron microscopy (TEM) were performed as previously described with modifications (10). Briefly, *C. neoformans* cultures were grown in liquid YPD at 30°C for 48 h and harvested by centrifugation. Cell pellets were resuspended in PBS containing 3% Triton X-100 and disrupted by vortexing with glass beads. Cell debris was removed by centrifugation, and the resulting supernatant was filtered through a 0.22 μm membrane. Virus-like particles were enriched by ultracentrifugation through a 20% sucrose cushion at 25,000 rpm for 2 h at 4°C. The particle-containing fraction was resuspended in PBS and maintained on ice until analysis. Samples were negatively stained and visualized by transmission electron microscopy at the Center for Electron Microscopy & Nanoscale Technology, Department of Pathology, Duke University.

#### Independent remapping of public CnTV1-positive libraries

To assess the distribution of CnTV1 among *C. neoformans* strains, we initially screened 128 publicly available paired-end RNA-seq libraries representing 34 strains from dataset GSE171092 (11). Primary reads mapping to the RACE-confirmed CnTV1 genome were counted for each library, and coverage profiles were examined to determine whether reads were distributed across the viral genome. At this preliminary screening stage, libraries containing ≥1,000 CnTV1-mapped reads were provisionally categorized as high-read-count candidates, whereas those containing 100–999 mapped reads were categorized as medium-read-count candidates. Libraries with fewer than 100 mapped reads were considered low-confidence or negative and were excluded from subsequent analysis. This screening identified 15 putative CnTV1-positive libraries, comprising 12 high-read-count and three medium-read-count candidates from 10 strains. To independently validate and visualize CnTV1 signals, we retrieved these 15 candidate-positive publicly available paired-end RNA-seq libraries from dataset GSE171092 (11) using SRA Toolkit v3.4.1. No additional *in silico* host-genome or rRNA depletion was performed before alignment. Each raw library was mapped directly to the RACE-confirmed CnTV1 genome using Bowtie2 (7) with –local –very-sensitive-local -k 10, matching the mapping parameters used for the RNAi-loss strain panel. Properly paired mapped reads were extracted with SAMtools (8) (SAM flag -F 260), sorted, and indexed. Per-base read depth was computed with samtools depth -a, mapped read counts were obtained with SeqKit bam, and total raw read counts per library were computed with SeqKit (4) stats. Mean depth, median depth, and breadth of coverage (≥1x) were calculated for each library from the resulting depth profiles. Within this preselected candidate-positive set, libraries with a mean per-base CnTV1 depth of ≥50x were classified as high-confidence, whereas those with a mean depth of <50x were classified as medium-confidence. Independent remapping classified 12 libraries as high-confidence and three as medium-confidence, consistent with their preliminary read-count categories.

**Phylogenetic analysis of CnTV1 RdRp**

To determine the phylogenetic placement of CnTV1, the predicted amino acid sequence of its RNA-dependent RNA polymerase (RdRp) was compared against representative members of all recognized species within the order *Ghabrivirales*. A dataset of 181 RdRp amino acid sequences was assembled, comprising one representative sequence per classified species from all 19 families and 23 genera across the three suborders *Alphatotivirineae* (130 sequences), *Betatotivirineae* (47 sequences), and *Gammatotivirineae* (4 sequences). Protein accession numbers and taxonomic assignments for all sequences used are provided in SI Dataset 4. Multiple sequence alignment was performed using MAFFT v7 with the L-INS-i algorithm (12), which employs an iterative refinement strategy incorporating local pairwise alignment information and is particularly suited for sequences with conserved motifs embedded within variable regions, as is the case for viral RdRp domains. The resulting alignment was manually inspected and trimmed to remove poorly aligned regions and positions dominated by gaps, yielding a final alignment of 1,427 amino acid positions. Phylogenetic inference was conducted using FastTree 2.2.0 (13) with default parameters, which employs an approximate maximum-likelihood method with the JTT (Jones–Taylor–Thornton) amino acid substitution model and a CAT approximation for rate heterogeneity across sites. Branch support values were estimated using the Shimodaira–Hasegawa (SH) test implemented in FastTree, with values ranging from 0 to 1. Pairwise amino acid sequence identities between CnTV1 and its phylogenetically closest relatives were calculated for both the RdRP domain and the complete cap–pol fusion protein (gag–pol). Species demarcation criteria followed the guidelines established in the 2023 ICTV reorganization of *Ghabrivirales*, in which viruses sharing <70% RdRp amino acid sequence identity are considered different species, irrespective of host origin. This threshold replaced the former *Totiviridae* criterion of <50% identity across all encoded proteins (14, 15). Species naming followed the ICTV-ratified binomial format, with epithets derived from Latinized Japanese numbers assigned sequentially by publication date.

**Population screen and confirmation of the *AGO1*-NUMT**

Paired-end reads from two population cohorts of *C. neoformans* were screened for the 609 bp *COX2*-derived NUMT insertion in *AGO1* (CNAG_04609; breakpoint ≈ chr10:895,524): 387 globally distributed isolates spanning VNI, VNII, VNBI and VNBII (16) and an independent 677-isolate VNI collection (17). Reads were mapped with BWA-MEM (v0.7.18) (18) to a targeted three-contig reference comprising wild-type H99 *AGO1* (REF), the NRHc5010 insertion allele (ALT), and the *COX2*-adjacent mitochondrial source sequence. Each contig was verified as single-copy against the H99 genome by

BLAST+ (v2.17.0) (19). An isolate was called a candidate carrier when it retained no clean REF-spanning read across the breakpoint together with ≥ 2 *AGO1* - mitochondrial discordant pairs or two-sided junction support, and candidates were confirmed by re-mapping to the complete H99 nuclear + mitochondrial genome, requiring ≥ 2 discordant pairs whose mitochondrial mates concentrated at the *COX2* zone (≥ 80% of mates; > 5x the genome-wide mitochondrial-homology background). The two cohorts were screened and reported separately, and lineage association was tested by Fisher's exact test on the lineage-diverse Desjardins cohort (16).

To test whether the NUMT insertion in *AGO1* arose once or recurrently, we compared each carrier's genome-wide phylogenetic position with its phylogenetic position at the *AGO1* locus. The genome-wide relationships were taken from the 388-taxon whole-genome SNP phylogeny (available at https://doi.org/10.6084/m9.figshare.7399427) (16). For the locus tree, we reconstructed phylogenetic relationships across the *AGO1* insertion locus using two 30-kb windows flanking the NUMT breakpoint (chr10:865,525–895,524 and chr10:895,525–925,524; H99 coordinates). For each strain, the corresponding flanking regions were extracted from a genome assembly. The sequences for NRHc5010, NRHc5028, A2-102-5, D17-1, wild-type H99, and the RNAi-deficient isolate Bt210 were obtained from their finished genome assemblies. Bt114 and Bt116, for which only Illumina reads were available, were *de novo* assembled with SPAdes (v4.3.0) (20). The four genome-wide nearest neighbors of the carriers, RTC5, Bt211, Jp1088, and AD1-86a, were assembled with the same SPAdes procedure. In each assembly, the *AGO1* locus was identified by BLAST analysis using a short H99 sequence spanning the breakpoint, and the 30-kb regions extending upstream and downstream of the breakpoint were extracted from the corresponding contig. All four nearest-neighbor assemblies contained the complete 30-kb windows on both sides of the breakpoint. The 5’ and 3’ flanking regions were aligned separately with MAFFT (v7.525) (21) and subsequently concatenated, producing a 60,027-bp alignment. A maximum-likelihood tree was inferred with IQ-TREE (v3.1.2) (22) under the HKY+F+I model selected by ModelFinder, with 1,000 ultrafast-bootstrap replicates. Support for the monophyly of the NUMT carriers was independently evaluated with a neighbor-joining tree based on p-distances with 1,000 bootstrap replicates. The six carriers formed a single clade with 100% ultrafast-bootstrap support.

**Small RNA sequencing and analysis**

Total RNA was extracted using the mirVana miRNA Isolation Kit (Thermo Fisher Scientific, catalog# AM1561). Small RNA sequencing libraries were generated from total RNA using the QIAseq miRNA Library Kit (Qiagen, catalog# 331502) and sequenced on the Illumina NovaSeq X Plus or NextSeq P2 platform (1 × 75 bp) at the Duke University Sequencing and Genomic Technologies Core facility.

Small RNA reads were adapter-trimmed using Cutadapt (v5.2) (23). To identify potential contaminating noncoding RNAs, rRNA and tRNA sequences were predicted from the reference genome using Barrnap (https://github.com/tseemann/barrnap) (v0.9) and tRNAscan-SE (v2.0) (24), respectively. Trimmed reads were aligned against the predicted rRNA and tRNA sequences using Bowtie (v1.3.1) (25), and matching reads were removed. The remaining reads were mapped to the reference genomes using Bowtie (v1.3.1) (25) with a 14-nt seed, one permitted seed mismatch, and random reporting of a single valid alignment for multimapping reads (-n 1 -l 14 -e 80 -M 1). Alignments were processed using SAMtools (v1.23.1) (8). Read-length distributions, 5' nucleotide frequencies, and genome-wide profiles of small RNA coverage, strandedness, and size distribution were generated from mapped reads using a modified Perl script (https://github.com/Jhuang90/smallRNA) and a modified sRNA_Viewer script (https://github.com/Jhuang90/sRNA_Viewer). Data visualization was performed in R (v4.5.0) using RStudio (v2026.05.0+218).

**Nanopore sequencing and genome assembly**

Nanopore sequencing and *de novo* genome assembly for NRHc5028, NRHc5010, D17-1 and A2-102-5 were performed as previously described (2).

### CRISPR-Cas9 mediated allele exchange and gene deletion

CRISPR–Cas9-mediated genome editing was performed as described previously (2). Briefly, for *AGO1* (CNAG_04609) or *QIP1* (CNAG_01423) allele exchange, the H99-derived donor was amplified directly from H99 genomic DNA. For the NRHc5010-derived donor, fusion PCR with primers list in Supplementary Table 2 was utilized to generate an *AGO1* donor lacking the NUMT insertion. Transformations were performed with the TRACE protocol (26, 27). Strains were transformed with a Cas9-encoding amplicon, guide RNAs targeting the intended editing site and Safe Haven 1 (SH1) (28), and donor constructs for both the intended site and SH1. Candidate transformants were screened by PCR spanning the *AGO1* or *QIP1* locus, with primers located outside the donor DNA boundaries to confirm correct integration. Complete removal of the NUMT insertion was further verified by amplicon sequencing (Plasmidsaurus). Unique sequence variants, likely introduced during transformation or genome editing, were identified in some independently generated transformants; however, these variants were not shared among independent transformants and were therefore unlikely to account for the reproducible phenotypes observed.

For targeted gene deletion of *NUC1* (CNAG_02204), *REX2* (CNAG_00288), *SKI2* (CNAG_04131), and *XRN1* (CNAG_02175), donor constructs containing 60-nt microhomology arms (27) were used. Correct replacement of each target gene was confirmed by PCR with assays targeting the internal coding region and the 5' and 3' integration junctions.

### Chemical curing of the mycovirus

NRHc5010 and NRHc5028 were cultured at 37°C in 8 mL YPD supplemented with cycloheximide at final concentrations of 50, 100, 200, or 500 ng/mL. Cultures were passaged every 48 h for a total of 10 passages. At passages 5 and 10, cultures were plated, and six independent colonies from each condition were isolated for RNA extraction followed by RT-PCR analysis to assess CnTV1 persistence.

To further promote viral curing, cultures were also grown in YPD containing 3 μg/mL ribavirin in combination with 100 ng/mL cycloheximide. Independent CnTV1-cured derivatives of NRHc5010 were recovered after five passages from cultures treated with either 50 ng/mL cycloheximide alone or 3 μg/mL ribavirin plus 100 ng/mL cycloheximide. In contrast, no CnTV1-cured isolates were identified from NRHc5028 under any treatment condition. RT-PCR analysis of all colonies isolated from passage 5 and passage 10 confirmed persistent CnTV1 RNA, with no evidence of viral clearance.

**SNP calling**

Whole-genome paired-end Illumina reads from five independent CnTV1-cured derivatives of NRHc5010 (JHG225, JHG227, JHG228, TCD1, TCD3) were aligned to the H99 reference genome and variants were called with Snippy (v4.6.0) (https://github.com/tseemann/snippy) (BWA-MEM alignment, FreeBayes variant calling) under default quality, mapping, and coverage filters. Variant functional impact was annotated with SnpEff (v5.0e) (29) with a custom database built from the H99 CNA3.31 sequence and the curated H99.10p.aATGcorrected.2019-05-15 gene models, with each variant assigned its most severe overlapping effect (impact rank HIGH > MODERATE > LOW > MODIFIER). Two independent NRHc5010 datasets (one from a previously published study (16) and the other generated in this study) served together as the parental-background control, and reads from strain H99 (2) were processed identically as a self-versus-self specificity control. For each cured strain, variants coinciding by chromosome and position with a call in either NRHc5010 control were removed, and the residual candidate variants were classified by cross-strain recurrence as sample-specific (one strain), recurrent (two to four strains), or shared (all five strains). To distinguish pre-existing parental sequence from curing-induced changes, read-level pileups (bcftools mpileup) (30) were generated at each shared site in both control alignments and the parental allele depth were recorded.

**Flow cytometry analysis**

Flow cytometric was employed to assess DNA content and ploidy was performed as previously described (31).

**Progeny ancestry analysis.**

For the progeny ancestry analysis, whole-genome paired-end Illumina reads were aligned to the H99 reference with BWA-MEM (v0.7.18) and processed with SAMtools (v1.20) (8). Variants were jointly called within each cross panel (Set 1, KN99**a** *ago1*Δ × NRHc5010; Set 2, KN99**a** *ago1*Δ × NRHc5028) with BCFtools (v1.21) (30) using bcftools mpileup | bcftools call --ploidy 1 (69), and single-nucleotide variants were retained after filtering to biallelic sites with QUAL ≥ 30 and per-sample depth DP ≥ 10 (indels excluded). Parent-informative markers were defined directly from the two cross parents' genotypes as sites where the parental alleles differed (51,708 markers for Set 1; 49,821 markers for Set 2). Each progeny genome was scored along the 14 nuclear chromosomes in non-overlapping 50-kb windows; each 50-kb window was assigned to a parent when ≥90% of its confidently genotyped informative markers (DP ≥ 10) supported that parent; windows below this threshold were scored as mixed, and windows with fewer than 5 such markers were left unassigned.

**Genome-wide coverage analysis.**

Whole-genome paired-end Illumina reads were aligned to the H99 reference genome using minimap2 (32) in short-read mode (-ax sr). Alignments were filtered to retain reads with mapping quality ≥ 30, sorted, and indexed using SAMtools (8). Genome-wide sequencing depth was then calculated using mosdepth (33) in non-overlapping 500-bp windows. For each sample, the mean coverage of each window was normalized to the median coverage across all nuclear-chromosome windows, such that a normalized coverage of 1 represented the genome-wide median. Normalized coverage profiles were visualized across the nuclear chromosomes in Rstudio.

**Spot dilution assays**

Biomass from strains grown on solid YPD medium was inoculated into 5 mL of liquid YPD, then grown overnight at 30°C in a roller drum rotating at 70 rpm. After overnight growth, the cultures were centrifuged at 3,200 rpm for 5 min to harvest the cells, then washed twice and resuspended in sterile $dH_2O$. Each culture was normalized to an $O.D._{600}$ of 0.5 and then serially diluted 5-fold in sterile $dH_2O$. 3 µL of each dilution was spotted onto solid media and allowed to air-dry under a flame. The plates were then incubated at 30°C, 37°C, 37°C+5%$CO_2$, or 39°C for 2 days prior to imaging.

**Capsule and melanin production assays**

To assess capsule production, biomass from strains grown on solid YPD medium was inoculated into liquid RPMI (RPMI 1640 with 2% dextrose) and diluted to obtain 4 mL cultures with an initial $O.D._{600}$ of 0.1. The cultures were incubated at 37°C in a roller drum rotating at 70 rpm for 3 days. The cultures were centrifuged at 3,200 rpm for 5 min to harvest the cells, washed once, and resuspended in sterile $dH_2O$. Equal volumes of resuspended culture and India ink were mixed to negatively stain the cells and visualize the capsules. Cells were imaged with a Zeiss Axioskop 2 microscope equipped with an AxioCam MRm digital camera. For quantification, at least 50 cells per culture were manually measured with Fiji (ImageJ version 1.54g). Capsule thickness was calculated using the formula: capsule thickness = (total diameter – cell body diameter)/2.

To assess melanin production, biomass from strains grown on solid YPD medium was inoculated into 5 mL of liquid YPD, and the cultures were grown to saturation at 30°C in a roller drum rotating at 70 rpm. 10 µL of each saturated culture was spotted onto Niger seed agar (7% Niger seed, 0.1% dextrose), and the plates were incubated at 30°C or 37°C for at least 1 week prior to imaging.

**RNA-seq and differential expression analysis.**

RNA was extracted from overnight YPD liquid cultures with the mirVana miRNA Isolation Kit (Thermo Fisher Scientific, catalog# AM1561) according to the manufacturer's instructions. Total RNA-seq was performed for KN99**a** *ago1*Δ, BLOSSOM-generated CnTV1B$^{+}$ (JHG514), and CnTV1B$^{0}$ (JHG538), with all three strains in the KN99**a** *ago1*Δ background and three biological replicates per strain. Libraries were sequenced on an Illumina NovaSeq X Plus platform using 2 x 150-bp paired-end sequencing at the Duke University Sequencing and Genomic Technologies Core facility. Reads were adapter- and quality-trimmed with fastp (v0.24.0) (34) and aligned using STAR (v2.7.11b; two-pass mode) (35) to a modified H99 reference genome with *MAT*α hard-masked and supplemented with *MAT***a** and the CnTV1 sequence. Gene-level counts were generated with featureCounts (Subread v2.1.1; -s 2 -p --countReadPairs) (36) for 8,322 host genes and the CnTV1 Gag and Pol coding sequences. KN99**a** *ago1*Δ replicate 1 was excluded after identification as an outlier by PCA, leaving eight libraries for downstream analyses (KN99**a** *ago1*Δ, n = 2; CnTV1B$^{+}$ and CnTV1B$^{0}$, n = 3 each). Sample relationships were visualized by PCA using variance-stabilizing-transformed counts from the 500 most variable genes.

Differential expression analysis was performed with DESeq2 (v1.50.2 in R v4.5.3) (37). Genes with ≥ 10 counts in at least three libraries were retained, leaving 7,809 genes for differential expression analysis. Differential expression was assessed using the DESeq2 Wald test, with Benjamini–Hochberg correction for multiple testing applied separately within each contrast (adjusted $p$ value < 0.05). An additional 1.5-fold change threshold (|$\log_2$ fold change| > 0.585) was applied to distinguish genes exhibiting larger transcriptional changes from those meeting the statistical significance threshold alone. For subsequent analyses, including the Venn diagram and identification of shared transcriptional responses, genes were selected based on an adjusted $p$ value < 0.05 without an additional fold-change cutoff, given the generally modest magnitude of CnTV1-associated transcriptional changes. $\text{Log}_2$ fold changes were estimated using adaptive shrinkage (lfcShrink, type = “ashr”), whereas statistical significance was determined using Wald tests on unshrunken coefficient estimates. The 1.5-fold change threshold was applied to the shrunken $\log_2$ fold-change estimates.

CnTV1-derived features were excluded before DESeq2 normalization; parallel analyses retaining viral features produced nearly identical size factors and results. Differential expression was evaluated for CnTV1B$^+$ versus KN99**a** *ago1*Δ, CnTV1B$^+$ versus CnTV1B$^0$, and CnTV1B$^0$ versus KN99**a** *ago1*Δ. Over-representation analysis of the 294 shared genes was performed using a one-sided hypergeometric test implemented in clusterProfiler (enricher function), with 4,995 annotated genes used as the background. Benjamini–Hochberg correction was applied for multiple testing, and no GO terms reached an adjusted $p$ value < 0.05 (SI Dataset 3).

Gene-set enrichment analysis was performed on genes ranked by the DESeq2 Wald statistic using clusterProfiler (v4.18.4) (38) with the fgsea adaptive multilevel permutation procedure (weighted enrichment statistic, exponent = 1) and Gene Ontology annotations obtained from UniProt, with biological process, molecular function, and cellular component terms analyzed separately (gene-set size, 5–500; Benjamini–Hochberg correction within each ontology). A total of 681 GO terms were tested (280 biological process, 226 molecular function, and 175 cellular component). GO terms with an adjusted $p$ value < 0.05 were considered significantly enriched, yielding 48, 40, and 26 significant terms (summed across the three ontologies with multiple-testing correction performed separately for each ontology) for CnTV1B+ versus KN99**a** *ago1*Δ, CnTV1B+ versus CnTV1B$^0$, and CnTV1B$^0$ versus KN99**a** *ago1*Δ, respectively. Enrichment direction and magnitude were assessed using the normalized enrichment score (NES).

GO terms showing significant enrichment in the same direction in both CnTV1B$^+$ versus KN99**a** *ago1*Δ and CnTV1B$^+$ versus CnTV1B$^0$ comparisons (adjusted $p$ value < 0.05), but not in CnTV1B$^0$ versus KN99**a** *ago1*Δ (adjusted $p$ value ≥ 0.05), were considered CnTV1-responsive pathways for which statistically significant enrichment was no longer observed following viral curing. Twenty-three GO terms met these criteria. For visualization, five terms exhibiting residual enrichment trends in the CnTV1B$^0$ versus KN99**a** *ago1*Δ comparison (|NES| ≥ 1.3) were excluded. The remaining 18 GO terms are presented in Fig. 6F.

**Murine infection assay**

The *C. neoformans* inoculum was prepared by culturing the cells overnight in 5 mL of liquid YPD at 30°C in a roller drum rotating at 70 rpm. The cultures were centrifuged at 3,200 rpm for 5 min to harvest the cells and washed twice with sterile phosphate-buffered saline (PBS). The cells were resuspended in sterile PBS, and the cell density was determined with a hemocytometer and adjusted to 4 x $10^6$ cells/mL using sterile PBS. Four- to five-week-old A/J mice (Jackson Laboratory, USA) were intranasally infected in groups of 10 (5 male and 5 female per strain tested). All groups were evaluated in a single infection experiment. The mice were anesthetized with isoflurane via a calibrated vaporizer and intranasally instilled with 25 µL of inoculum (1 x $10^5$ cells). Survival was monitored daily, and mice were euthanized by $CO_2$ exposure upon reaching predefined humane endpoints, including weight loss exceeding 15% of the original body weight, reduced mobility, lack of grooming, social isolation, or a hunched posture. Survival differences between groups were assessed using the Gehan-Breslow-Wilcoxon test. Median survival times and their 95% confidence intervals were determined by Kaplan–Meier analysis using GraphPad Prism (v11.1.0). All animal procedures were performed in accordance with an approved IACUC protocol (protocol number A098-22-05-25).

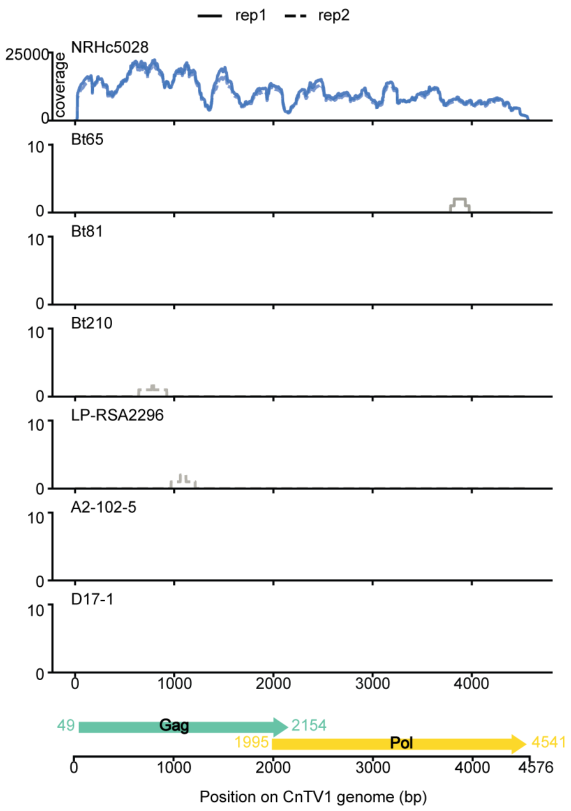


**Figure S1. NRHc5028 is infected with the novel mycovirus Cryptococcus neoformans totivirus 1 (CnTV1).** Total RNA sequencing was performed on seven previously identified RNAi-deficient isolates: NRHc5028, Bt65, Bt81, Bt210, LP-RSA2296, A2-102-5, and D17-1. Reads were mapped to the corresponding genome assembly, and unmapped reads were assembled *de novo*. Read coverage across the CnTV1 genome was detected exclusively in NRHc5028. No reads mapping to CnTV1 were detected in the other isolates. The 4,576 bp dsRNA CnTV1 genome is illustrated at the bottom and contains two untranslated regions (UTRs) and two partially overlapping open reading frames (ORFs) encoding the capsid protein (Gag) and RNA-dependent RNA polymerase (Pol). rep1 and rep2 indicate sequencing results from biological replicates.

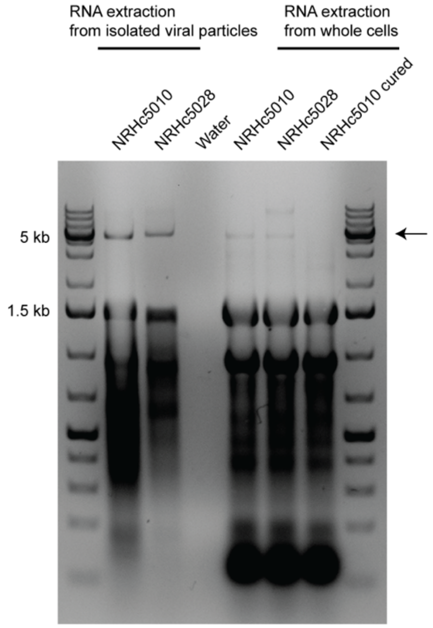


**Figure S2. CnTV1 RNA is enriched in viral particle extracts relative to whole-cell extracts.** Virus-like particles were isolated by ultracentrifugation through a sucrose cushion. CnTV1 RNA (~4.6 kb) in viral particle extracts and whole-cell extracts was detected by total RNA gel electrophoresis, revealing enrichment of the ~4.6 kb RNA species in viral particle extracts. The NRHc5010 cured strain and water served as negative controls. The arrow indicates the ~4.6 kb CnTV1 genome.

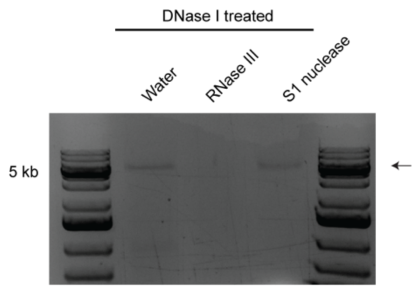


**Figure S3. CnTV1 has a dsRNA genome.** Purified viral nucleic acid was treated with DNase I and subsequently treated with water as a control, RNase III, or S1 nuclease. DNase I, RNase III, and S1 nuclease preferentially digest DNA, dsRNA, and single-stranded nucleic acids, respectively. Digests were analyzed by gel electrophoresis, revealing that the CnTV1 nucleic acid was resistant to DNase I and S1 nuclease treatment but sensitive to RNase III treatment, consistent with a dsRNA genome. The arrow indicates the ~4.6 kb CnTV1 genome.

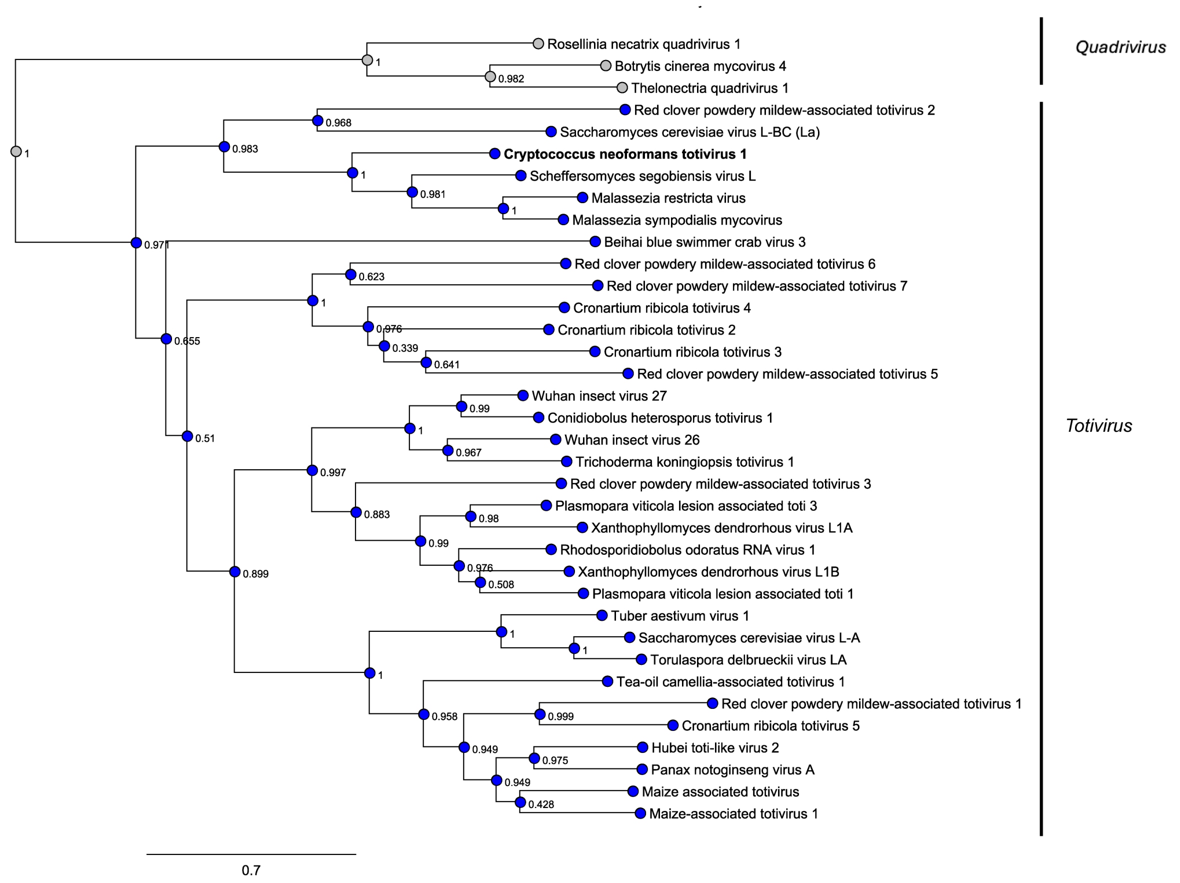


**Figure S4. Complete phylogeny of *Totivirus*.** Numbers at nodes indicate Shimodaira–Hasegawa branch support values, and the unit for the branch length is amino acid substitutions per site. This tree represents the complete phylogeny corresponding to the simplified version shown in Fig. 1D.

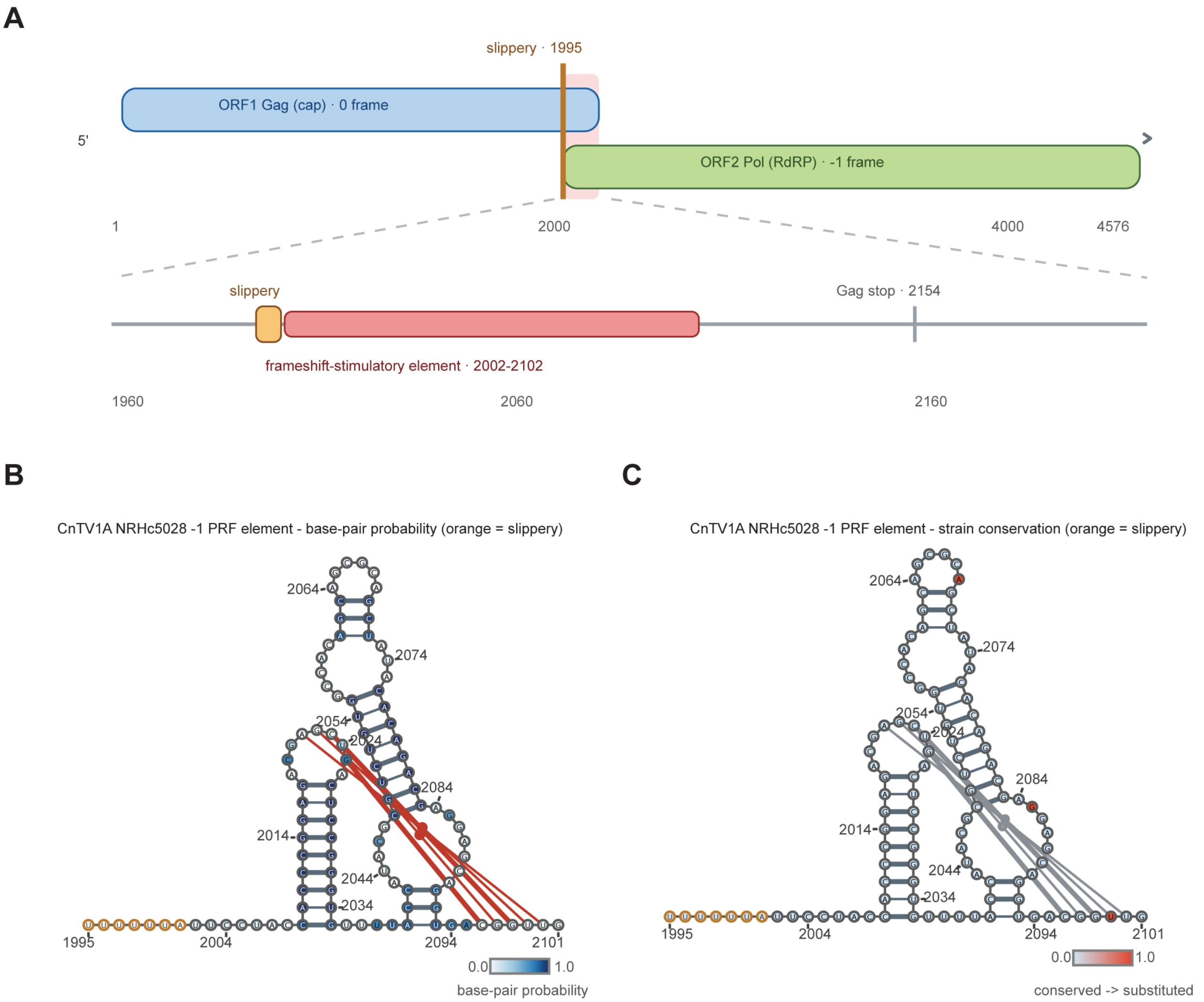


**Figure S5. Genomic organization and −1 programmed ribosomal frameshifting element of CnTV1.** (A) Genomic organization of CnTV1A from strain NRHc5028. ORF1 (Gag/cap, zero frame) and ORF2 (Pol/RdRp, −1 frame) overlap across the frameshift region. The slippery site (1995-2000) and the frameshift-stimulatory element (~2002-2102) are expanded in the lower track. (B) Secondary structure of the CnTV1A −1 PRF element from NRHc5028 rendered in VARNA. Nucleotides are colored by partition-function base-pair probability (blue scale); the slippery heptamer is shown in orange at the 5′ end; red arcs denote the pseudoknot (P2) pairs, which the nested partition function cannot score. Numbering is in genome coordinates. (C) Cross-strain (NRHc5028 and NRHc5010) sequence conservation mapped onto the −1 PRF element. Conserved positions are light blue, and substitutions are red; the slippery heptamer is orange. The substitution within the pseudoknot helix is a pairing-preserving G:C↔G:U change. Grey arcs denote the pseudoknot pairs.

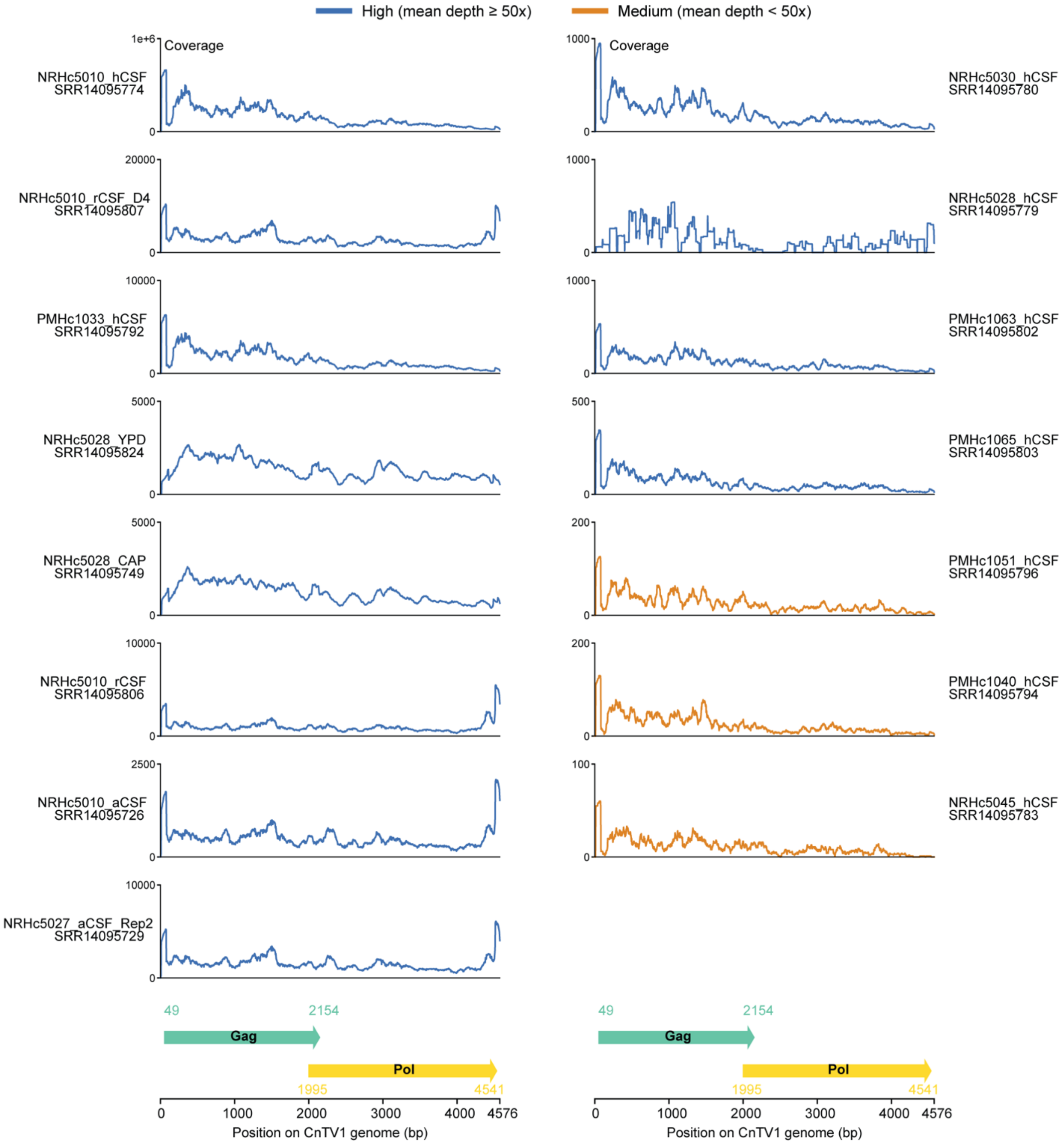


**Figure S6. Identification of CnTV1 sequences in publicly available RNA-seq datasets**. Per-base read coverage depth across the CnTV1 genome is shown for each of 15 independently remapped public libraries (GSE171092). Samples are grouped according to mean sequencing depth across the CnTV1 genome, with high-coverage samples (mean depth ≥50x) shown in blue and medium-coverage samples (mean depth <50x) shown in orange. Sample names, culture conditions, and SRA accession numbers are indicated for each sequencing experiment. Coverage depth is plotted against the position along the 4,576-bp CnTV1 genome. The genome organization of CnTV1 is shown below, with the Gag and Pol coding regions and their nucleotide coordinates indicated. Note: Read coverage is shown at different y-axis scales to visualize the very low-level CnTV1-mapped reads detected in several isolates.

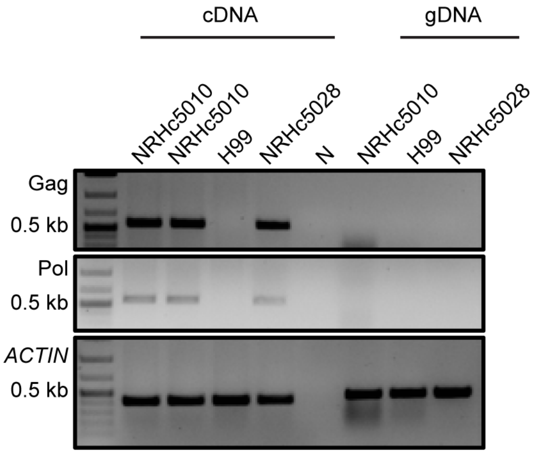


**Figure S7. NRHc5010 is infected with CnTV1.** CnTV1 was detected by targeted RT-PCR using complementary DNA (cDNA) as the template but was not detected by PCR using genomic DNA (gDNA). *ACTIN* served as a control. Water (N) and the reference strain H99 served as negative controls, whereas the CnTV1-infected NRHc5028 strain served as a positive control for virus presence. Lanes 1 and 2 represent biological replicates of NRHc5010.

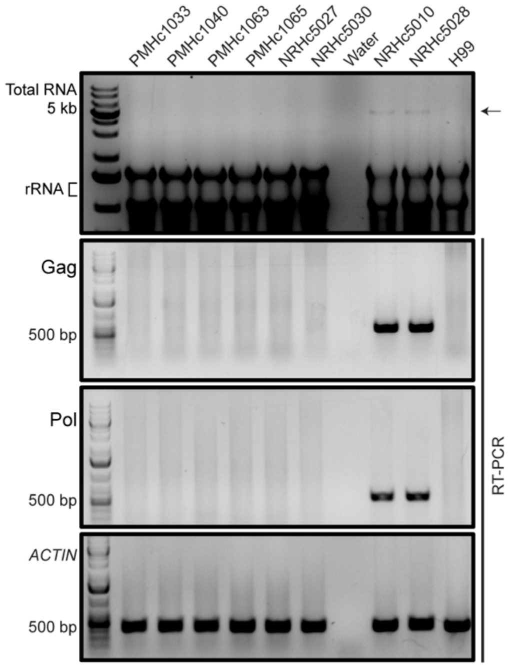


**Figure S8. CnTV1 was not detected in other isolates with lower levels of CnTV1-mapping RNA-seq reads.** Isolates with detectable CnTV1-mapping reads in at least one RNA-seq library were screened for CnTV1 by total RNA gel electrophoresis and targeted RT-PCR. CnTV1 was detected only in NRHc5010 and NRHc5028. *ACTIN* served as a PCR control. Water and the reference strain H99 served as negative controls, whereas the CnTV1-infected strains NRHc5010 and NRHc5028 served as positive controls for virus presence. The arrow indicates the ~4.6 kb CnTV1 genome.

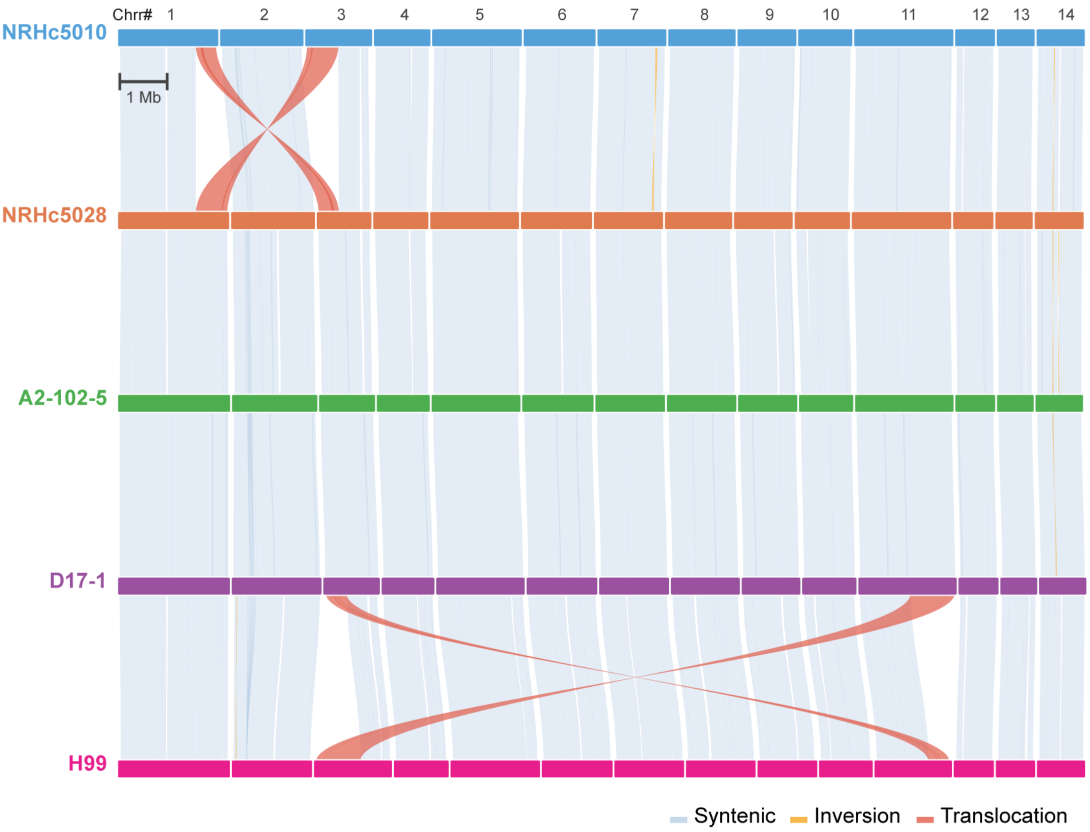


**Figure S9. Whole-genome synteny comparison among RNAi-deficient *C. neoformans* isolates and the H99 reference strain.** Chromosome-scale genome alignments are shown for NRHc5010, NRHc5028, A2-102-5, D17-1, and H99. Chromosomes are arranged from Chr1 to Chr14, and connecting ribbons indicate syntenic relationships between adjacent genomes. Light blue ribbons denote collinear syntenic regions, whereas yellow and red ribbons indicate inversions and translocations, respectively. While extensive genome-wide synteny is retained among the isolates, strain NRHc5010 contains a private reciprocal translocation between chromosomes 1 and 3, and the reference strain H99 contains a private reciprocal translocation between chromosomes 3 and 11. The scale bar represents 1 Mb.

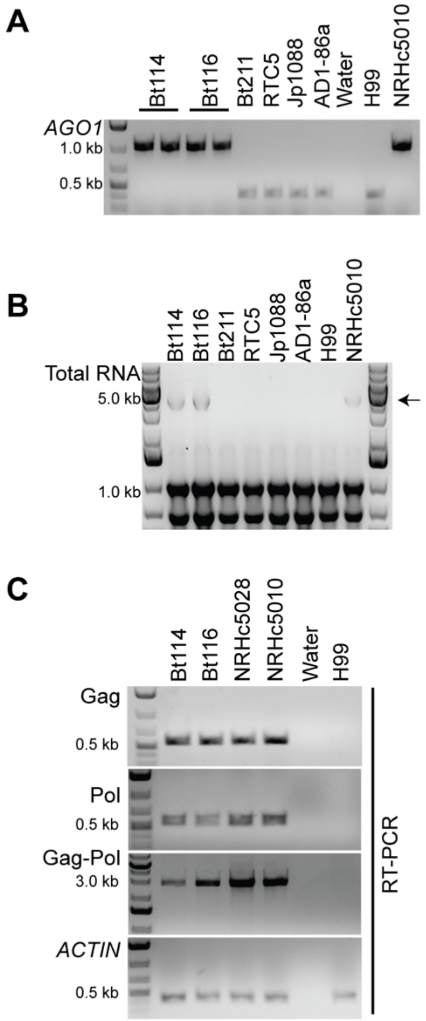


**FigS10. Identification of CnTV1 infection in Bt114 and Bt116 carrying the *AGO1* NUMT insertion.** (A) PCR analysis of the *AGO1* locus across the indicated *C. neoformans* isolates. NRHc5010 was included as a positive control for the *AGO1* NUMT insertion. (B) Agarose gel electrophoresis of total RNA extracted from the indicated isolates, revealing the presence of an RNA species similar in size to the CnTV1 genome in Bt114 and Bt116. H99 and NRHc5010 served as negative and positive controls for CnTV1 presence, respectively. The arrow indicates the ~4.6 kb CnTV1 genome. (C) RT-PCR detection of CnTV1 using primers targeting the Gag, Pol, and Gag-Pol regions in Bt114, Bt116, NRHc5028, and NRHc5010. *ACTIN* served as a control. Water and H99 served as negative controls.

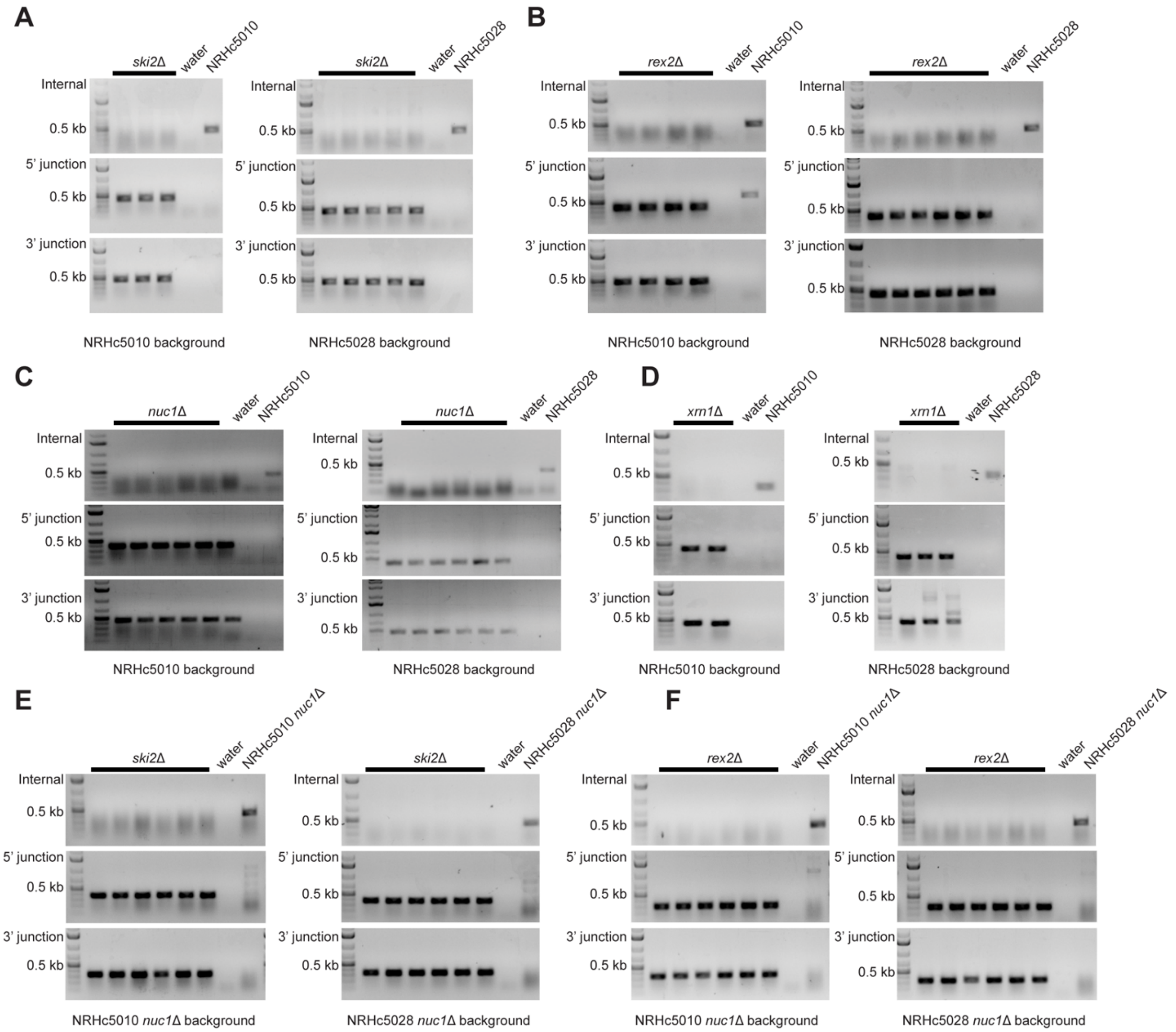


**Figure S11. Validation of deletion mutants in NRHc5010 and NRHc5028.** (A) Validation of *ski2*Δ mutants in NRHc5010 (left) and NRHc5028 (right). (B) Validation of *rex2*Δ mutants in NRHc5010 (left) and NRHc5028 (right). (C) Validation of *nuc1*Δ mutants in NRHc5010 (left) and NRHc5028 (right). (D) Validation of *xrn1*Δ mutants in NRHc5010 (left) and NRHc5028 (right). (E) Validation of *ski2*Δ mutants in NRHc5010 *nuc1*Δ (left) and NRHc5028 *nuc1*Δ (right). (F) Validation of *rex2*Δ mutants in NRHc5010 *nuc1*Δ (left) and NRHc5028 *nuc1*Δ (right). (A-F) Deletion mutants were validated by genotyping PCR, confirming loss of the open reading frame (ORF) by internal PCR and successful integration of the dominant selectable marker at the endogenous locus by 5' and 3' junction PCRs. Water and genomic DNA (gDNA) from the corresponding progenitor strain used for transformation served as controls.

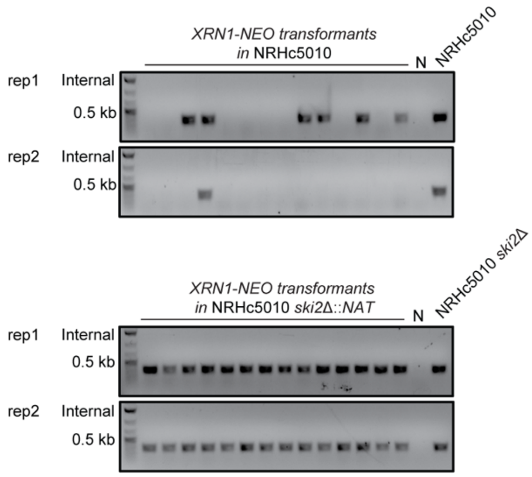


**Figure S12. NRHc5010 *ski2*Δ *xrn1*Δ mutants could not be recovered by CRISPR-Cas9-mediated gene editing.** *xrn1*Δ mutants were successfully recovered in the wild-type NRHc5010 background (top) but not in the NRHc5010 *ski2*Δ background (bottom), consistent with a negative genetic interaction between *SKI2* and *XRN1*. Transformants were screened for loss of the *XRN1* open reading frame (ORF) by internal PCR. Water (N) and genomic DNA (gDNA) from the corresponding progenitor strain used for transformation served as controls. rep1 and rep2 indicate results from independent transformations.

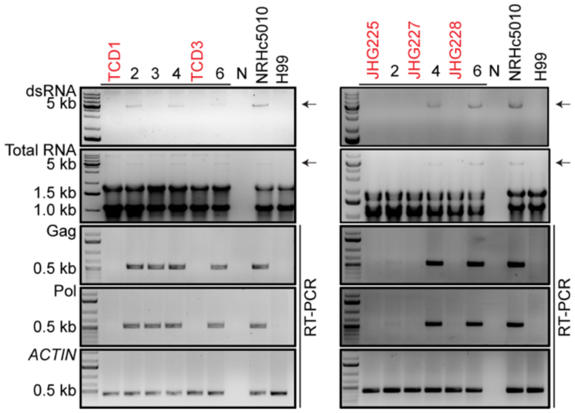


**Figure S13. NRHc5010 was chemically cured of CnTV1.** The CnTV1B-cured strains TCD1 and TCD3 were generated by passaging NRHc5010 every 48 h in liquid YPD supplemented with 50 ng/mL cycloheximide at 37°C for a total of 5 passages (left). The CnTV1B-cured strains JHG225, JHG227, and JHG228 were generated by passaging NRHc5010 every 48 h in liquid YPD supplemented with 100 ng/mL cycloheximide and 3 µg/mL ribavirin at 37°C for a total of five passages (right). CnTV1 was assessed by dsRNA-enriched and total RNA gel electrophoresis and by targeted RT-PCR. *ACTIN* served as a PCR control. Water (N) and H99 served as negative controls, whereas unpassaged NRHc5010 served as a positive control for virus presence. Arrows indicate the ~4.6 kb CnTV1 genome.

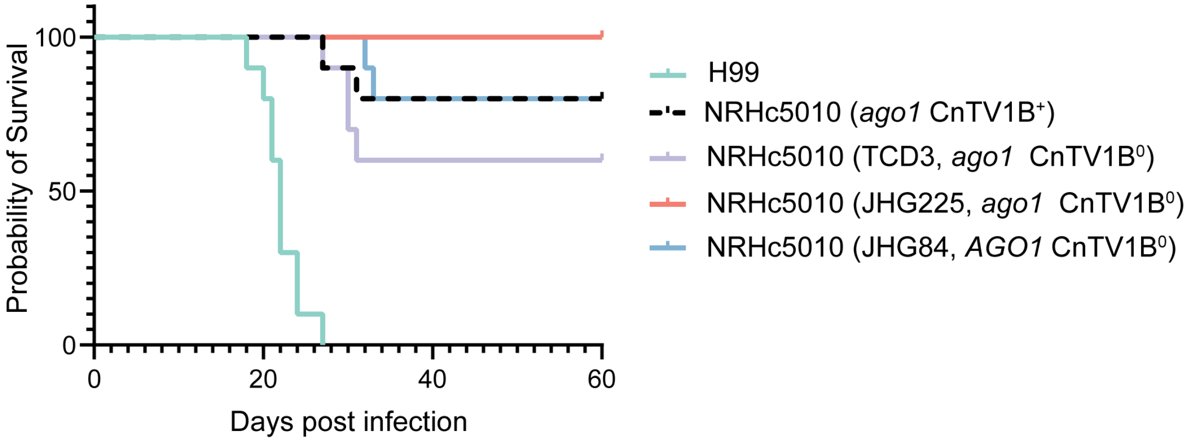


**Figure S14. NRHc5010 and its CnTV1-cured derivatives exhibit low virulence in a murine inhalation model of infection**. The CnTV1B-infected NRHc5010 strain, chemically cured derivatives TCD3 and JHG225, and genetically cured JHG84 (*AGO1*) strain were used to infect equal numbers of male and female A/J mice (n = 10 per group). Mice were intranasally inoculated with $10^5$ cells. Survival differences between groups were assessed using the Gehan-Breslow-Wilcoxon test. No significant differences in survival were observed between NRHc5010 and its CnTV1B-cured derivatives. Notably, all mice infected with the reference strain H99 succumbed to infection by 27 days post-infection (dpi), whereas most mice infected with NRHc5010 or its CnTV1B-cured derivatives survived until the experiment was terminated at 60 dpi. CnTV1B$^+$, CnTV1B-infected; CnTV1B$^0$, CnTV1B-cured.

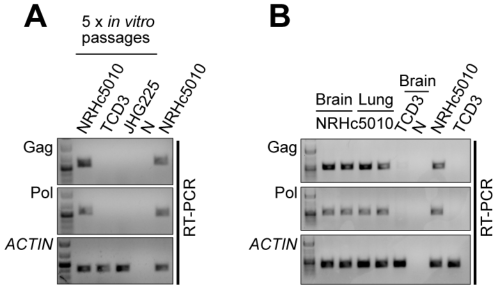


**Figure S15. CnTV1 infection status of NRHc5010 and its cured derivatives is stable *in vitro* and *in vivo*.** (A) The CnTV1B-infected NRHc5010 strain and its chemically cured derivatives, TCD3 and JHG225, were serially passaged five times *in vitro* and subsequently assessed for the presence or absence of CnTV1. (B) NRHc5010, TCD3, and JHG225 isolates recovered from the brains or lungs of infected mice were assessed for the presence or absence of CnTV1. (A, B) CnTV1 was detected by targeted RT-PCR. *ACTIN* served as a PCR control. Water (N) served as a negative control. The unpassaged CnTV1-infected NRHc5010 strain served as a positive control.

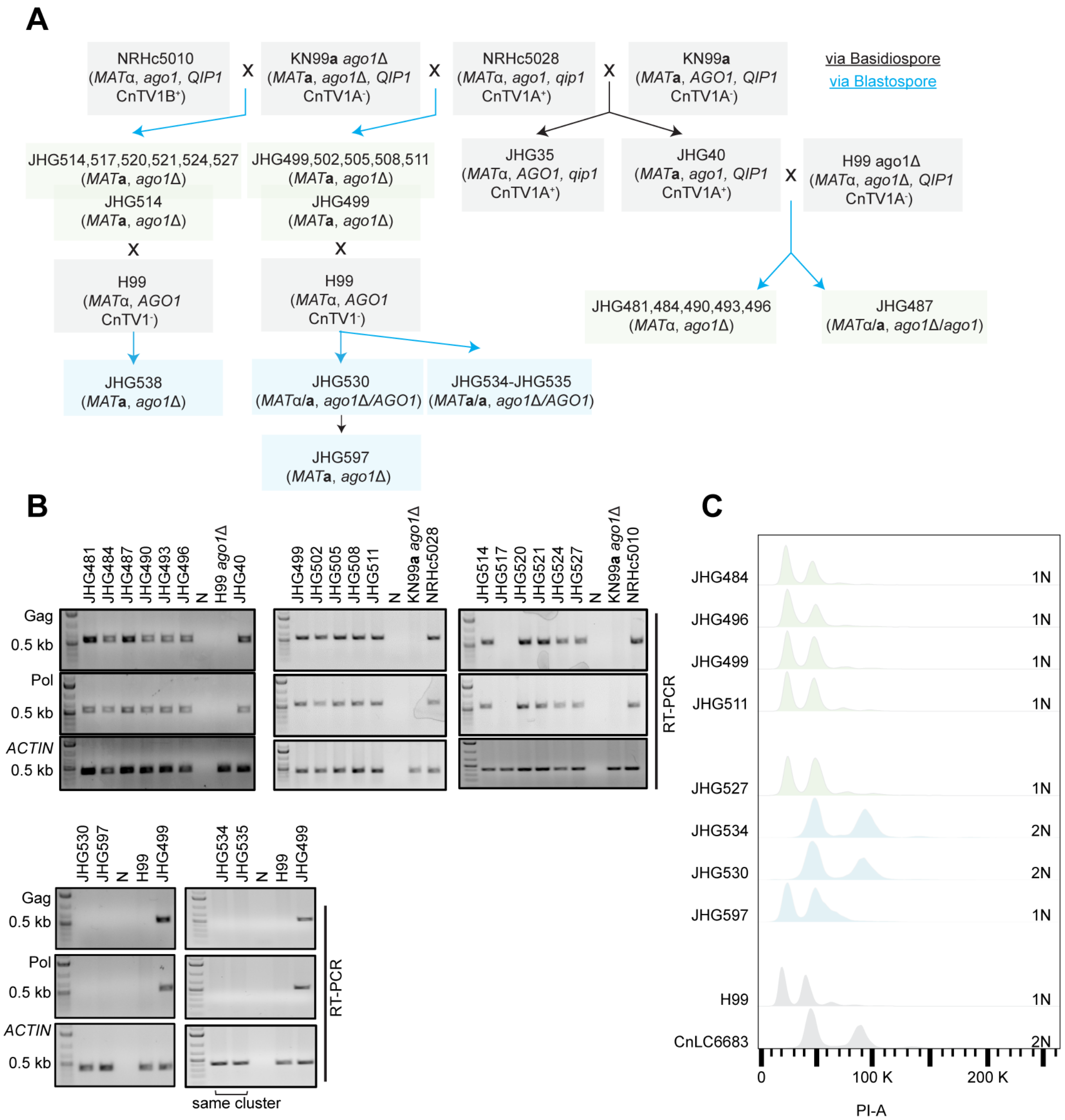


**Figure S16. Strain genealogy, CnTV1 detection, and ploidy analysis of strains generated using the BLOSSOM method.** (A) Strain genealogy showing the generation of the indicated progeny and derivative strains using the BLOSSOM method. Parental genotypes, mating types, and CnTV1 infection status are indicated. Blue and black arrows denote progeny isolated via blastospore and basidiospore dissection, respectively. (B) CnTV1 infection status of the indicated strains was assessed by RT-PCR using primers targeting the viral Gag and Pol regions. *ACTIN* served as a control. Strains derived from the same blastospore cluster are indicated. (C) Ploidy analysis of representative dissected blastospores by fluorescence-activated cell sorting (FACS). Most analyzed blastospores exhibited haploid (1N) profiles, whereas JHG534 and JHG530 exhibited diploid (2N) profiles. H99 and CnLC6683 served as haploid and diploid controls, respectively.

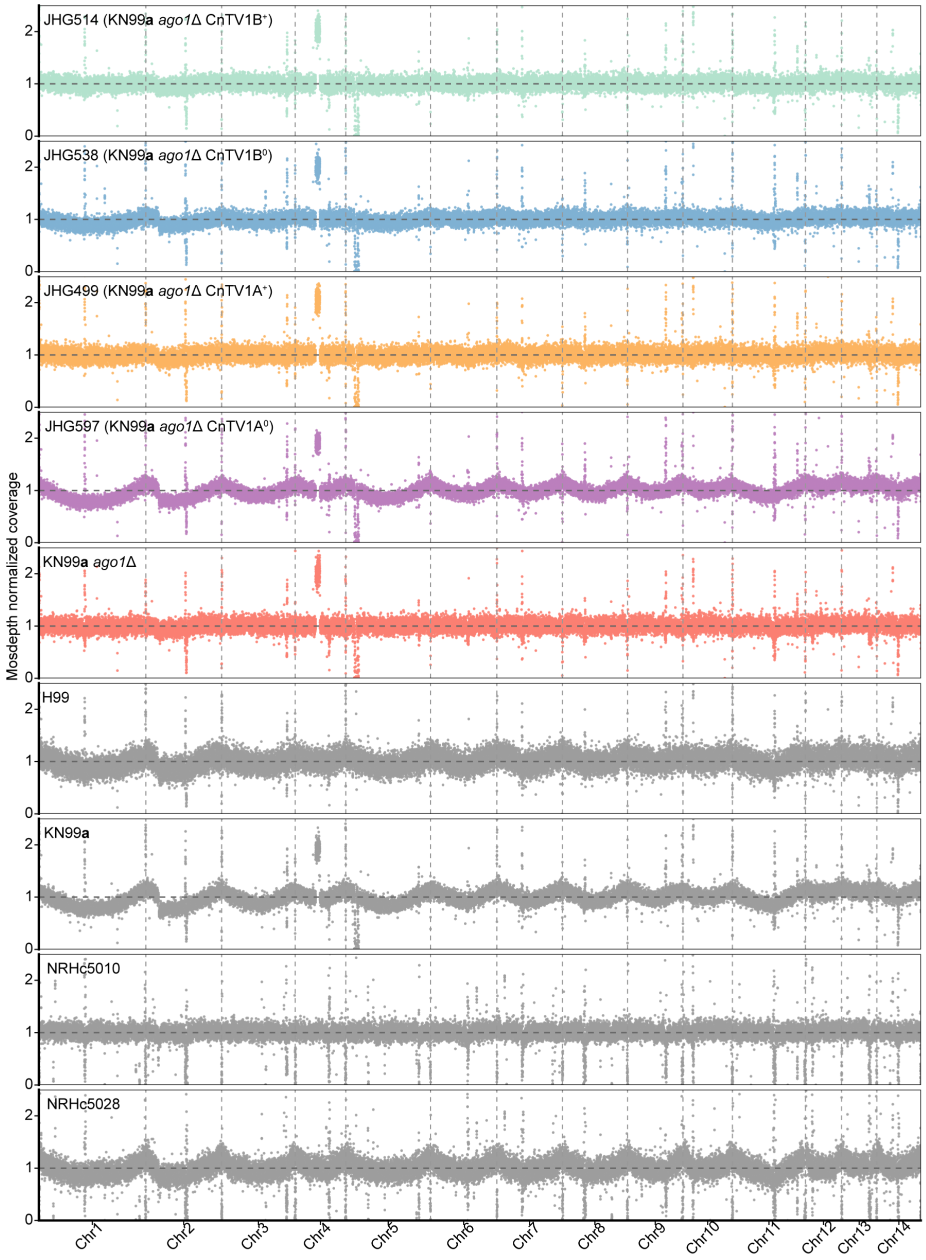


**Figure S17. Isogenic CnTV1-infected and -cured strain pairs generated by BLOSSOM are euploid.** CnTV1-infected (JHG514, CnTV1B$^{+}$; JHG499, CnTV1A$^{+}$) and CnTV1-cured (JHG538, CnTV1B$^{0}$; JHG597, CnTV1A$^{0}$) strains were generated in the KN99**a** *ago1*Δ background via BLOSSOM. The generated strains and their progenitors were subjected to whole-genome sequencing. Sequencing reads were mapped to the H99 reference genome, and read depth was calculated using mosdepth and normalized to the genome-wide median coverage for each strain. Normalized read depth is plotted across all 14 chromosomes, with chromosome boundaries indicated by vertical lines. All strains showed comparable chromosome-wide normalized read depth profiles, with no evidence of aneuploidy.

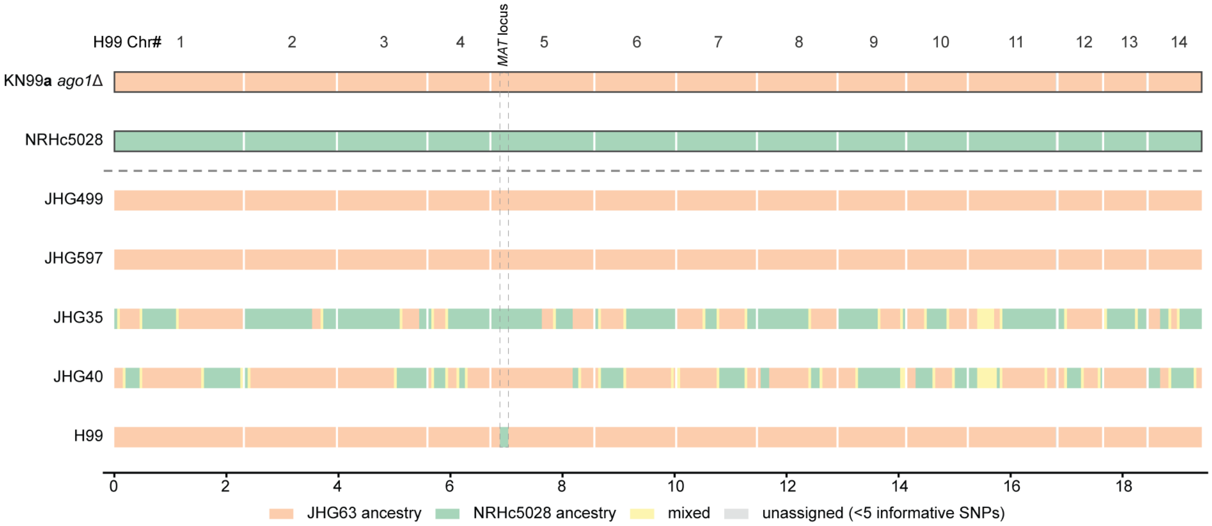


**Figure S18. Genome-wide ancestry analysis of BLOSSOM-derived progeny.** Ancestry was assigned using informative SNPs that distinguish the KN99**a** *ago1*Δ and NRHc5028 parental backgrounds. JHG499 and JHG597, generated using BLOSSOM, showed predominantly KN99**a** *ago1*Δ ancestry across the nuclear genome. In contrast, JHG35 and JHG40, meiotic progeny derived from a cross between NRHc5028 and KN99**a**, exhibited extensive recombinantion across their genomes. H99 is shown for comparison. Colors indicate JHG63 ancestry, NRHc5028 ancestry, mixed ancestry, and regions with insufficient informative SNPs for ancestry assignment. The *MAT* locus is indicated by dashed vertical lines.

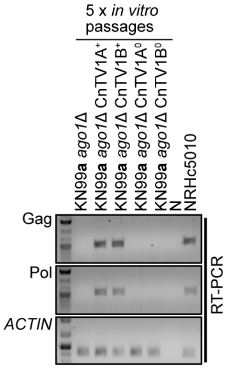


**Figure S19. CnTV1 infection status is stable in BLOSSOM-generated strains during *in vitro* passage.** CnTV1-infected (JHG499, CnTV1A$^{+}$; JHG514, CnTV1B$^{+}$) and -cured (JHG597, CnTV1A$^{o}$; JHG538, CnTV1B$^{o}$) strains generated by BLOSSOM in the KN99**a** *ago1*Δ background were serially passaged five times *in vitro* on YPD medium. CnTV1 was detected by RT-PCR using primers targeting the viral Gag and Pol regions. *ACTIN* served as a control. NRHc5010 was included as a CnTV1-positive control.

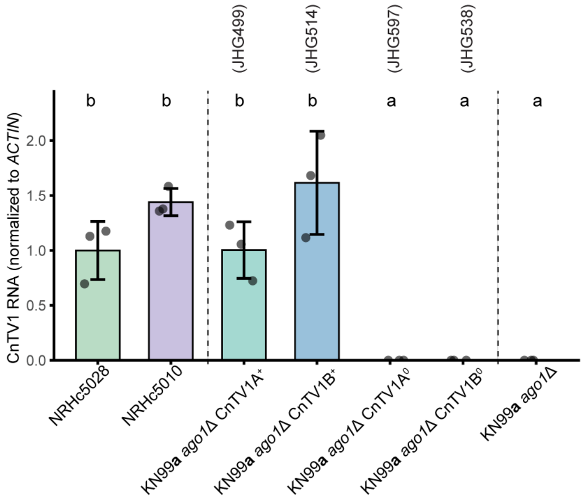


**Figure S20. CnTV1 RNA levels are similar among infected strains and undetectable in cured strains.** CnTV1A and CnTV1B RNA levels are similar in the natural isolates (NRHc5028 and NRHc5010) and their corresponding infected strains (CnTV1A$^{+}$ and CnTV1B$^{+}$) in the KN99**a** *ago1*Δ background. KN99**a** *ago1*Δ and the CnTV1-cured strains (CnTV1A$^{0}$ and CnTV1B$^{0}$) do not contain detectable CnTV1 RNA. CnTV1 RNA levels were quantified by RT-qPCR with the $2^{-\Delta\Delta Ct}$ method and normalized to the Actin-encoding gene *ACT1*. Statistical analyses were performed on ΔCt values using a one-way ANOVA followed by Tukey's honestly significant difference (HSD) post hoc test. Different letters at the top of the plot indicate significantly different RNA levels.

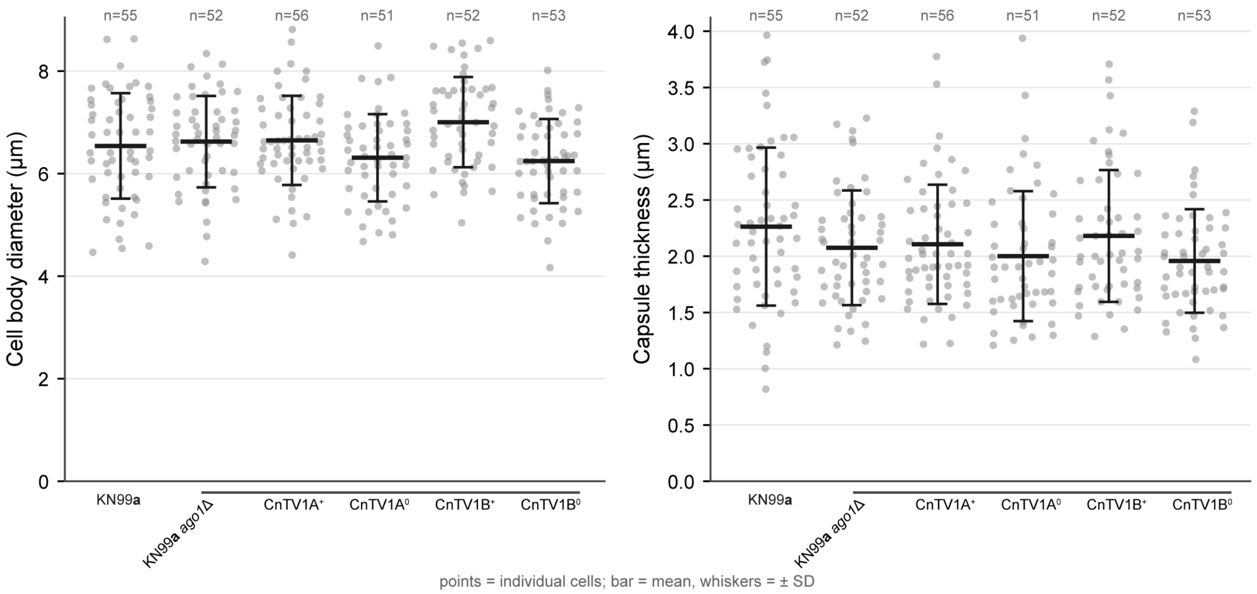


**Figure S21. Isogenic CnTV1-infected and -cured strains exhibit similar cell and capsule size.** Cells were grown in liquid RPMI at 37°C for 3 days to induce capsule production. At least 50 cells per strain were measured with Fiji/ImageJ. Cell body diameter and capsule thickness are shown on the left and right, respectively. Capsule thickness was calculated by subtracting the cell body diameter from the total cell diameter and dividing by 2. Each point represents a single cell. Horizontal bars indicate strain means, and error bars represent standard deviations. CnTV1A$^{+}$ and CnTV1B$^{+}$ strains are CnTV1-infected, while CnTV1A$^{0}$ and CnTV1B$^{0}$ are the isogenic cured strains. n indicates the number of cells measured.

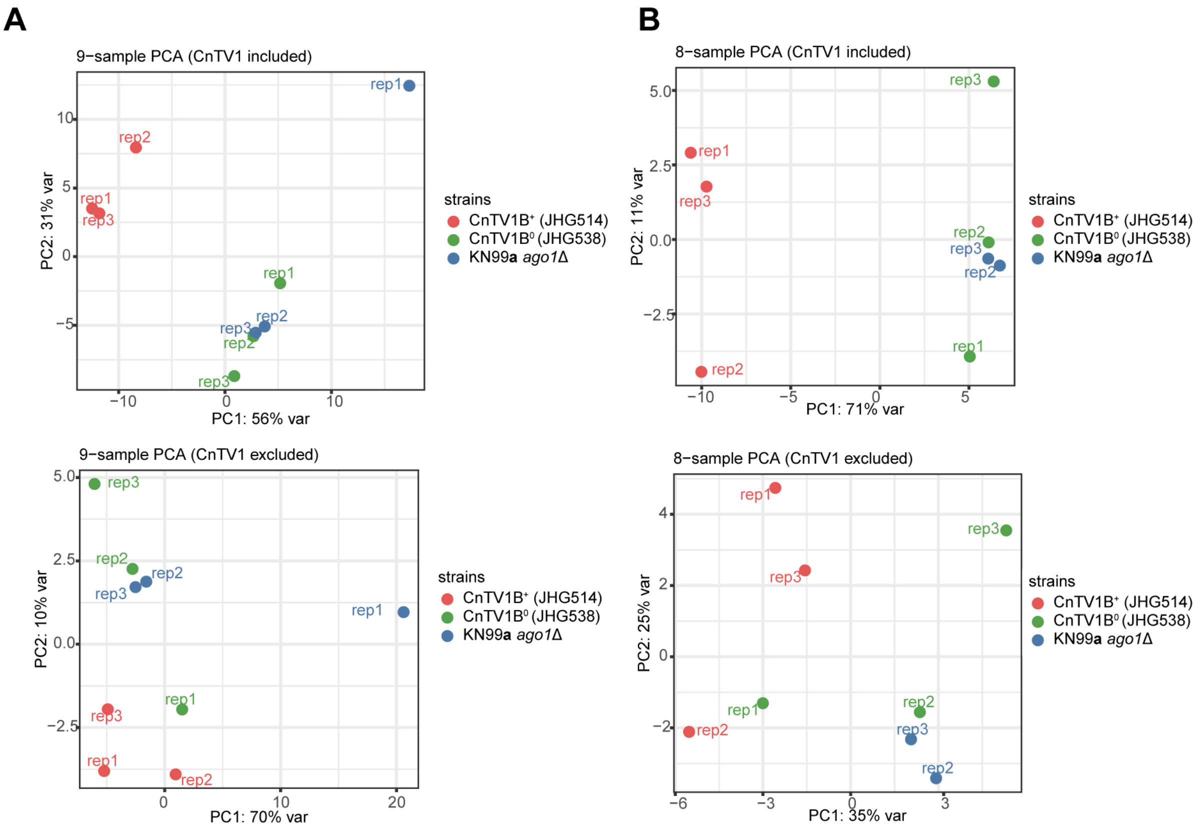


**Figure S22. Principal component analysis of host transcriptomes with and without CnTV1-derived reads.** (A) PCA of all nine RNA-seq samples: three biological replicates each of CnTV1B-infected JHG514, CnTV1-cured JHG538, and the isogenic virus-free parental strain KN99**a** *ago1*Δ. Counts were filtered to retain genes with ≥10 counts in ≥3 samples, variance-stabilized using DESeq2 (VST, blind = TRUE), and PCA was performed on the 500 most variable genes. CnTV1-derived reads were either included (top) or excluded (bottom) from the analysis. (B) The same analysis after removing one outlier KN99**a** *ago1*Δ replicate (rep1), with CnTV1-derived reads included (top) or excluded (bottom). Percentages on the axes indicate the variance explained by each principal component.

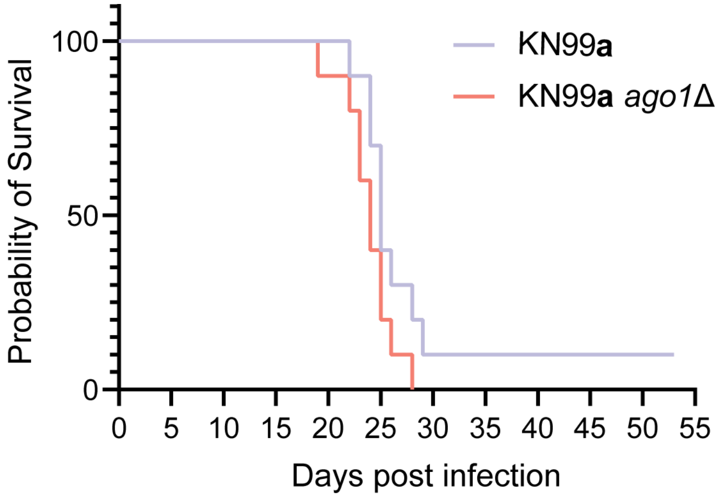


**Figure S23. Loss of *AGO1* does not significantly alter virulence in *C. neoformans*.** Survival of A/J mice infected with wild-type KN99**a** or KN99**a** *ago1*Δ in an inhalation model of cryptococcosis. The KN99**a** *ago1*Δ group is the same group shown in Fig. 6G, and all virulence assays shown here and in Fig. 6G were performed concurrently as part of the same experiment. The data are displayed separately here to facilitate direct comparison between KN99**a** and KN99**a** *ago1*Δ. No significant difference in survival was observed between the two strains. Statistical significance was assessed using the Gehan–Breslow–Wilcoxon test.

**Table S1. Strains used in this study.**

| Strain name/designation | Stock number | Genotype | Note | Source |
|---|---|---|---|---|
| H99 | JOHE4413 | *MAT*α | *C. neoformans* natural isolate, VNI lineage | (39) |
| Bt210 | JOHE5170 | *MAT*α | *C. neoformans* natural isolate, VNI lineage | (1, 40) |
| LP-RSA2296 | JOHE22081 | *MAT*α | *C. neoformans* natural isolate, VNI lineage | (1, 40) |
| A2-102-5 | JOHE22038 | *MAT*α | *C. neoformans* natural isolate, VNI lineage | (1, 40) |
| D17-1 | JOHE22039 | *MAT*α | *C. neoformans* natural isolate, VNI lineage | (1, 40) |
| NRHc5028.ENR.STOR | JOHE22792 | *MAT*α | *C. neoformans* natural isolate, VNI lineage, CnTV1+ | (1, 40) |
| Bt65 | JOHE5067 | *MAT***a** | *C. neoformans* natural isolate, VNBII lineage | (1, 40) |
| Bt81 | JOHE5082 | *MAT***a** | *C. neoformans* natural isolate, VNBII lineage | (1, 40) |
| H99 *ago1*Δ | JOHE4933 | *MAT*α *ago1*Δ::*NAT* | H99 *ago1*Δ, RNAi mutant | (41) |
| H99 *rdp1*Δ | JOHE6840 | *MAT*α *rdp1*Δ::*NEO* | H99 *rdp1*Δ, RNAi mutant | (42) |
| NRHc5010.ENR | JOHE22199 | *MAT*α | *C. neoformans* natural isolate, VNI lineage, CnTV1+ | (40) |
| Bt114 | JOHE5113 | *MAT*α | *C. neoformans* natural isolate, VNI lineage, CnTV1+ | (40) |
| Bt116 | JOHE5115 | *MAT*α | *C. neoformans* natural isolate, VNI lineage, CnTV1+ | (40) |
| PMHc1033.ENR | JOHE22002 | *MAT*α | *C. neoformans* natural isolate, VNBII lineage | (40) |
| PMHc1040.ENR.STOR | JOHE22192 | *MAT*α | *C. neoformans* natural isolate, VNI lineage | (40) |
| PMHc1063.ENR.STOR | JOHE25735 | *MAT***a** | *C. neoformans* natural isolate, VNBII lineage | (11) |
| PMHc1065.ENR.STOR | JOHE25736 | *MAT*α | *C. neoformans* natural isolate, VNI lineage | (11) |
| NRHc5027.ENR.CLIN1 | JOHE22213 | *MAT*α | *C. neoformans* natural isolate, VNBI lineage | (40) |
| NRHc5030.ENR.CLIN.ISO | JOHE22215 | *MAT***a** | *C. neoformans* natural isolate, VNBI lineage | (40) |
| Bt211 | JOHE5171 | *MAT*α | *C. neoformans* natural isolate, VNI lineage | (40) |
| RTC5 | JOHE22082 | *MAT*α | *C. neoformans* natural isolate, VNI lineage | (40) |
| Jp1088 | JOHE22285 | *MAT*α | *C. neoformans* natural isolate, VNI lineage | (40) |
| AD1-86a | JOHE22059 | *MAT*α | *C. neoformans* natural isolate, VNI lineage | (40) |
| KN99**a** | JOHE3259 | *MAT***a** | *C. neoformans* lab isolate | (43) |
| CnLC6683 | JOHE21994 | *MAT*α/**a** | *C. neoformans* lab isolate, diploid | (44) |
| H99 *cap59*Δ | JOHE3127 | *MAT*α *cap59*Δ::*HYG* | *C. neoformans* acapsular strain, H99 *cap59*Δ | (45) |
| MCD16 | JOHE5905 | *MAT*α *lac1*Δ::*URA5* | *C. neoformans* strain deficient in melanin production, H99 *lac1*Δ | (46) |
| KN99**a** *ago1*Δ (JHG63) | JOHE25618 | *MAT***a** *ago1*Δ::*NAT* | RNAi mutant, generated by crossing H99 *ago1*Δ x KN99**a** | This study |
| KN99**a** *qip1*Δ (JHG75) | JOHE25619 | *MAT***a** *qip1*Δ::*NEO* | RNAi mutant, generated by CRISPR-Cas9 editing | This study |

| | | | | |
|---|---|---|---|---|
| JHG25 | JOHE25620 | *MAT*α *qip1 AGO1* | F1 progeny from NRHc5028 x KN99**a**; CnTV1- | This study |
| JHG26 | JOHE25621 | *MAT*α *qip1 ago1* | F1 progeny from NRHc5028 x KN99**a**; CnTV1- | This study |
| JHG27 | JOHE25622 | *MAT***a** *QIP1 AGO1* | F1 progeny from NRHc5028 x KN99**a**; CnTV1- | This study |
| JHG28 | JOHE25623 | *MAT*α *qip1 ago1* | F1 progeny from NRHc5028 x KN99**a**; CnTV1- | This study |
| JHG29 | JOHE25624 | *MAT*α *qip1 ago1* | F1 progeny from NRHc5028 x KN99**a**; CnTV1- | This study |
| JHG30 | JOHE25625 | *MAT***a** *QIP1 AGO1* | F1 progeny from NRHc5028 x KN99**a**; CnTV1- | This study |
| JHG31 | JOHE25626 | *MAT*α *qip1 ago1* | F1 progeny from NRHc5028 x KN99**a**; CnTV1- | This study |
| JHG32 | JOHE25627 | *MAT***a** *QIP1 AGO1* | F1 progeny from NRHc5028 x KN99**a**; CnTV1- | This study |
| JHG33 | JOHE25628 | *MAT***a** *QIP1 AGO1* | F1 progeny from NRHc5028 x KN99**a**; CnTV1- | This study |
| JHG34 | JOHE25629 | *MAT***a** *QIP1 AGO1* | F1 progeny from NRHc5028 x KN99**a**; CnTV1- | This study |
| JHG35 | JOHE25630 | *MAT*α *qip1 AGO1* | F1 progeny from NRHc5028 x KN99**a**; CnTV1+ | This study |
| JHG36 | JOHE25631 | *MAT*α *qip1 AGO1* | F1 progeny from NRHc5028 x KN99**a**; CnTV1- | This study |
| JHG37 | JOHE25632 | *MAT*α *QIP1 AGO1* | F1 progeny from NRHc5028 x KN99**a**; CnTV1- | This study |
| JHG38 | JOHE25633 | *MAT*α *qip1 AGO1* | F1 progeny from NRHc5028 x KN99**a**; CnTV1- | This study |
| JHG39 | JOHE25634 | *MAT***a** *QIP1 AGO1* | F1 progeny from NRHc5028 x KN99**a**; CnTV1- | This study |
| JHG40 | JOHE25635 | *MAT***a** *QIP1 ago1* | F1 progeny from NRHc5028 x KN99**a**; CnTV1+ | This study |
| JHG41 | JOHE25636 | *MAT***a** *QIP1 AGO1* | F1 progeny from NRHc5028 x KN99**a**; CnTV1- | This study |
| JHG42 | JOHE25637 | *MAT*α *qip1 AGO1* | F1 progeny from NRHc5028 x KN99**a**; CnTV1- | This study |
| JHG43 | JOHE25638 | *MAT***a** *QIP1 AGO1* | F1 progeny from NRHc5028 x KN99**a**; CnTV1- | This study |
| JHG44 | JOHE25639 | *MAT*α *qip1 ago1* | F1 progeny from NRHc5028 x KN99**a**; CnTV1- | This study |
| JHG45 | JOHE25640 | *MAT***a** *QIP1 AGO1* | F1 progeny from NRHc5028 x KN99**a**; CnTV1- | This study |
| JHG46 | JOHE25641 | *MAT*α *qip1 ago1* | F1 progeny from NRHc5028 x KN99**a**; CnTV1- | This study |
| JHG47 | JOHE25642 | *MAT***a** *qip1 ago1* | F1 progeny from NRHc5028 x KN99**a**; CnTV1- | This study |
| JHG48 | JOHE25643 | *MAT*α *qip1 AGO1* | F1 progeny from NRHc5028 x KN99**a**; CnTV1- | This study |
| JHG49 | JOHE25644 | *MAT***a** *QIP1 AGO1* | F1 progeny from NRHc5028 x KN99**a**; CnTV1- | This study |
| JHG50 | JOHE25645 | *MAT*α *qip1 ago1* | F1 progeny from NRHc5028 x KN99**a**; CnTV1- | This study |
| JHG51 | JOHE25646 | *MAT*α *qip1 AGO1* | F1 progeny from NRHc5028 x KN99**a**; CnTV1- | This study |
| JHG52 | JOHE25647 | *MAT***a** *QIP1 AGO1* | F1 progeny from NRHc5028 x KN99**a**; CnTV1- | This study |
| JHG53 | JOHE25648 | *MAT***a** *QIP1 ago1* | F1 progeny from NRHc5028 x KN99**a**; CnTV1- | This study |
| JHG105 | JOHE25649 | *MAT***a** *QIP1 ago1*Δ | F1 progeny from NRHc5028 x KN99**a** *ago1*Δ; CnTV1+ | This study |
| JHG106 | JOHE25650 | *MAT*α *qip1 ago1*Δ | F1 progeny from NRHc5028 x KN99**a** *ago1*Δ; CnTV1+ | This study |
| JHG107 | JOHE25651 | *MAT*α *qip1 ago1*Δ | F1 progeny from NRHc5028 x KN99**a** *ago1*Δ; CnTV1+ | This study |

| | | | | |
|---|---|---|---|---|
| JHG108 | JOHE25652 | *MAT***a** *QIP1 ago1Δ* | F1 progeny from NRHc5028 x KN99**a** *ago1*Δ; CnTV1+ | This study |
| JHG109 | JOHE25653 | *MATα QIP1 ago1* | F1 progeny from NRHc5028 x KN99**a** *ago1*Δ; CnTV1+ | This study |
| JHG110 | JOHE25654 | *MATα QIP1 ago1* | F1 progeny from NRHc5028 x KN99**a** *ago1*Δ; CnTV1+ | This study |
| JHG111 | JOHE25655 | *MAT***a** *QIP1 ago1* | F1 progeny from NRHc5028 x KN99**a** *ago1*Δ; CnTV1+ | This study |
| JHG112 | JOHE25656 | *MATα qip1 ago1Δ* | F1 progeny from NRHc5028 x KN99**a** *ago1*Δ; CnTV1+ | This study |
| JHG113 | JOHE25657 | *MATα QIP1 ago1* | F1 progeny from NRHc5028 x KN99**a** *ago1*Δ; CnTV1+ | This study |
| JHG114 | JOHE25658 | *MAT***a** *qip1 ago1Δ* | F1 progeny from NRHc5028 x KN99**a** *ago1*Δ; CnTV1+ | This study |
| JHG115 | JOHE25659 | *MAT***a** *QIP1 ago1* | F1 progeny from NRHc5028 x KN99**a** *ago1*Δ; CnTV1+ | This study |
| JHG116 | JOHE25660 | *MATα qip1 ago1* | F1 progeny from NRHc5028 x KN99**a** *ago1*Δ; CnTV1+ | This study |
| JHG117 | JOHE25661 | *MATα QIP1 ago1Δ* | F1 progeny from NRHc5028 x KN99**a** *ago1*Δ; CnTV1+ | This study |
| JHG137 | JOHE25662 | *MAT***a** *qip1Δ AGO1* | F1 progeny from NRHc5028 x KN99**a** *qip1*Δ; CnTV1+ | This study |
| JHG138 | JOHE25663 | *MATα qip1 ago1* | F1 progeny from NRHc5028 x KN99**a** *qip1*Δ; CnTV1+ | This study |
| JHG139 | JOHE25664 | *MATα qip1 ago1* | F1 progeny from NRHc5028 x KN99**a** *qip1*Δ; CnTV1+ | This study |
| JHG140 | JOHE25665 | *MATα qip1 AGO1* | F1 progeny from NRHc5028 x KN99**a** *qip1*Δ; CnTV1+ | This study |
| JHG141 | JOHE25666 | *MATα qip1 ago1* | F1 progeny from NRHc5028 x KN99**a** *qip1*Δ; CnTV1+ | This study |
| JHG142 | JOHE25667 | *MATα qip1 ago1* | F1 progeny from NRHc5028 x KN99**a** *qip1*Δ; CnTV1+ | This study |
| JHG143 | JOHE25668 | *MATα qip1 ago1* | F1 progeny from NRHc5028 x KN99**a** *qip1*Δ; CnTV1+ | This study |
| JHG144 | JOHE25669 | *MAT***a** *qip1Δ ago1* | F1 progeny from NRHc5028 x KN99**a** *qip1*Δ; CnTV1+ | This study |
| JHG145 | JOHE25670 | *MATα qip1Δ AGO1* | F1 progeny from NRHc5028 x KN99**a** *qip1*Δ; CnTV1+ | This study |
| JHG146 | JOHE25671 | *MATα qip1Δ AGO1* | F1 progeny from NRHc5028 x KN99**a** *qip1*Δ; CnTV1+ | This study |
| JHG147 | JOHE25672 | *MATα qip1 ago1* | F1 progeny from NRHc5028 x KN99**a** *qip1*Δ; CnTV1+ | This study |
| JHG148 | JOHE25673 | *MATα qip1 ago1* | F1 progeny from NRHc5028 x KN99**a** *qip1*Δ; CnTV1+ | This study |
| JHG149 | JOHE25674 | *MATα qip1 ago1* | F1 progeny from NRHc5028 x KN99**a** *qip1*Δ; CnTV1+ | This study |
| JHG150 | JOHE25675 | *MATα qip1Δ ago1* | F1 progeny from NRHc5028 x KN99**a** *qip1*Δ; CnTV1+ | This study |
| JHG151 | JOHE25676 | *MAT***a** *qip1 AGO1* | F1 progeny from NRHc5028 x KN99**a** *qip1*Δ; CnTV1+ | This study |
| JHG152 | JOHE25677 | *MATα qip1 AGO1* | F1 progeny from NRHc5028 x KN99**a** *qip1*Δ; CnTV1+ | This study |
| JHG172 | JOHE25678 | *MATα QIP1 ago1* | F1 progeny from NRHc5028+*QIP1* x KN99**a** *qip1*Δ; CnTV1- | This study |
| JHG173 | JOHE25679 | *MAT***a** *qip1Δ AGO1* | F1 progeny from NRHc5028+*QIP1* x KN99**a** *qip1*Δ; CnTV1- | This study |
| JHG174 | JOHE25680 | *MAT***a** *qip1Δ ago1* | F1 progeny from NRHc5028+*QIP1* x KN99**a** *qip1*Δ; CnTV1- | This study |
| JHG175 | JOHE25681 | *MAT***a** *qip1Δ AGO1* | F1 progeny from NRHc5028+*QIP1* x KN99**a** *qip1*Δ; CnTV1- | This study |

| | | | | |
|---|---|---|---|---|
| JHG176 | JOHE25682 | *MAT***a** *qip1*Δ *AGO1* | F1 progeny from NRHc5028+*QIP1* x KN99**a** *qip1*Δ; CnTV1- | This study |
| JHG177 | JOHE25683 | *MAT***a** *qip1*Δ *AGO1* | F1 progeny from NRHc5028+*QIP1* x KN99**a** *qip1*Δ; CnTV1- | This study |
| JHG178 | JOHE25684 | *MAT***a** *qip1*Δ *AGO1* | F1 progeny from NRHc5028+*QIP1* x KN99**a** *qip1*Δ; CnTV1- | This study |
| JHG179 | JOHE25685 | *MAT***a** *qip1*Δ *AGO1* | F1 progeny from NRHc5028+*QIP1* x KN99**a** *qip1*Δ; CnTV1- | This study |
| JHG180 | JOHE25686 | *MAT*α *QIP1 ago1* | F1 progeny from NRHc5028+*QIP1* x KN99**a** *qip1*Δ; CnTV1- | This study |
| JHG181 | JOHE25687 | *MAT***a** *qip1*Δ *ago1* | F1 progeny from NRHc5028+*QIP1* x KN99**a** *qip1*Δ; CnTV1- | This study |
| JHG182 | JOHE25688 | *MAT***a** *qip1*Δ *ago1* | F1 progeny from NRHc5028+*QIP1* x KN99**a** *qip1*Δ; CnTV1- | This study |
| JHG183 | JOHE25689 | *MAT*α *qip1*Δ *ago1* | F1 progeny from NRHc5028+*QIP1* x KN99**a** *qip1*Δ; CnTV1- | This study |
| JHG184 | JOHE25690 | *MAT***a** *qip1*Δ *AGO1* | F1 progeny from NRHc5028+*QIP1* x KN99**a** *qip1*Δ; CnTV1- | This study |
| JHG185 | JOHE25691 | *MAT***a** *QIP1 ago1* | F1 progeny from NRHc5028+*QIP1* x KN99**a** *qip1*Δ; CnTV1- | This study |
| JHG186 | JOHE25692 | *MAT***a** *qip1*Δ *AGO1* | F1 progeny from NRHc5028+*QIP1* x KN99**a** *qip1*Δ; CnTV1- | This study |
| NRHc5010+*AGO1-1* (JHG84) | JOHE25693 | *MAT*α *AGO1* | *AGO1*-repaired, $CnTV1^0$, repair donor from H99 | This study |
| NRHc5010 $NEO^R$*-1* (JHG93) | JOHE25694 | *MAT*α *NEO* | RNAi-deficient, CnTV1+ | This study |
| JHG40+*AGO1* (JHG85) | JOHE25695 | *MAT***a** *AGO1* | *AGO1*-repaired, $CnTV1^0$, repair donor from H99 | This study |
| JHG35+*QIP1*-1 (JHG88) | JOHE25696 | *MAT*α *QIP1* | *QIP1*-repaired, $CnTV1^0$, repair donor from H99 | This study |
| JHG35+*QIP1*-2 (JHG89) | JOHE25697 | *MAT*α *QIP1* | *QIP1*-repaired, $CnTV1^0$, repair donor from H99 | This study |
| JHG35+*QIP1*-3 (JHG90) | JOHE25698 | *MAT*α *QIP1* | *QIP1*-repaired, $CnTV1^0$, repair donor from H99 | This study |
| NRHc5028+*QIP1* (JHG92) | JOHE25699 | *MAT*α *ago1 QIP1* | *QIP1*-repaired, RNAi-deficient, CnTV1+, repair donor from H99 | This study |
| NRHc5028 $NEO^R$ (JHG94) | JOHE25700 | *MAT*α *NEO* | RNAi-deficient, CnTV1+ | This study |
| NRHc5010+*AGO1-2* (JHG803) | JOHE25701 | *MAT*α *AGO1* | *AGO1*-repaired, $CnTV1^0$, repair donor from NRHc5010 | This study |
| NRHc5010+*AGO1-10* (JHG805) | JOHE25702 | *MAT*α *AGO1* | *AGO1*-repaired, $CnTV1^0$, repair donor from NRHc5010 | This study |
| NRHc5010 $NEO^R$*-2* (JHG806) | JOHE25703 | *MAT*α *NEO* | RNAi-deficient, CnTV1+ | This study |
| NRHc5010 *nuc1*Δ (JHG542) | JOHE25704 | *MAT*α *nuc1*Δ::*NAT* | NRHc5010 *nuc1*Δ::*NAT* | This study |
| NRHc5010 *ski2*Δ (JHG554) | JOHE25705 | *MAT*α *ski2*Δ::*NAT* | NRHc5010 *ski2*Δ::*NAT* | This study |
| NRHc5010 *rex2*Δ (JHG562) | JOHE25706 | *MAT*α *rex2*Δ::*NAT* | NRHc5010 *rex2*Δ::*NAT* | This study |
| NRHc5010 *xrn1*Δ (JHG626) | JOHE25707 | *MAT*α *xrn1*Δ::*NAT* | NRHc5010 *xrn1*Δ::*NAT* | This study |

| | | | | |
|---|---|---|---|---|
| NRHc5010 *nuc1Δ ski2Δ* (JHG602) | JOHE25708 | *MATα nuc1Δ::NAT ski2Δ::NEO* | NRHc5010 *nuc1Δ* (JHG542) *ski2Δ::NEO* | This study |
| NRHc5010 *nuc1Δ rex2Δ* (JHG608) | JOHE25709 | *MATα nuc1Δ::NAT rex2Δ::NEO* | NRHc5010 *nuc1Δ* (JHG542) *rex2Δ::NEO* | This study |
| NRHc5028 *nuc1Δ* (JHG548) | JOHE25710 | *MATα nuc1Δ::NAT* | NRHc5028 *nuc1Δ::NAT* | This study |
| NRHc5028 *ski2Δ* (JHG558) | JOHE25711 | *MATα ski2Δ::NAT* | NRHc5028 *ski2Δ::NAT* | This study |
| NRHc5028 *rex2Δ* (JHG566) | JOHE25712 | *MATα rex2Δ::NAT* | NRHc5028 *rex2Δ::NAT* | This study |
| NRHc5028 *xrn1Δ* (JHG628) | JOHE25713 | *MATα xrn1Δ::NAT* | NRHc5028 *xrn1Δ::NAT* | This study |
| NRHc5028 *nuc1Δ ski2Δ* (JHG614) | JOHE25714 | *MATα nuc1Δ::NAT ski2Δ::NEO* | NRHc5028 *nuc1Δ* (JHG548) *ski2Δ::NEO* | This study |
| NRHc5028 *nuc1Δ rex2Δ* (JHG620) | JOHE25715 | *MATα nuc1Δ::NAT rex2Δ::NEO* | NRHc5028 *nuc1Δ* (JHG548) *rex2Δ::NEO* | This study |
| KN99α *ski2Δ* (JHG783) | JOHE25716 | *MATα ski2Δ::NAT* | KN99α *ski2Δ::NAT* | (47) |
| KN99**a** *xrn1Δ* (JHG695) | JOHE25717 | *MAT***a** *xrn1Δ::NEO* | KN99**a** *xrn1Δ::NEO* | This study |
| KN99**a** *ski2Δ* (JHG728) | JOHE25718 | *MAT***a** *ski2Δ::NAT* | KN99**a** *ski2Δ::NAT*, F1 progeny from JHG783 x JHG695 | This study |
| NRHc5010 *ski2Δ+AGO1-1* (JHG807) | JOHE25719 | *MATα ski2Δ::NAT AGO1* | *AGO1*-repaired in JHG554, CnTV1$^0$ | This study |
| NRHc5010 *ski2Δ+AGO1-21* (JHG808) | JOHE25720 | *MATα ski2Δ::NAT AGO1* | *AGO1*-repaired in JHG554, CnTV1$^0$ | This study |
| NRHc5010 *ski2Δ+AGO1-28* (JHG809) | JOHE25721 | *MATα ski2Δ::NAT AGO1* | *AGO1*-repaired in JHG554, CnTV1$^0$ | This study |
| NRHc5010 *ski2Δ NEO$^R$* (JHG810) | JOHE25722 | *MATα ski2Δ::NAT NEO* | RNAi-deficient, CnTV1+ | This study |
| NRHc5010 *xrn1Δ+AGO1* (JHG692) | JOHE25723 | *MATα xrn1Δ::NAT AGO1* | *AGO1*-repaired in JHG626, CnTV1$^0$ | This study |
| TCD1 | JOHE25724 | *MATα* | Chemically cured, CnTV1$^0$ derivative of NRHc5010 | This study |
| TCD3 | JOHE25725 | *MATα* | Chemically cured, CnTV1$^0$ derivative of NRHc5010 | This study |
| JHG225 | JOHE25726 | *MATα* | Chemically cured, CnTV1$^0$ derivative of NRHc5010 | This study |
| JHG227 | JOHE25727 | *MATα* | Chemically cured, CnTV1$^0$ derivative of NRHc5010 | This study |
| JHG228 | JOHE25728 | *MATα* | Chemically cured, CnTV1$^0$ derivative of NRHc5010 | This study |
| JHG499 | JOHE25729 | *MAT***a** *ago1Δ::NAT* | BLOSSOM-generated CnTV1A+ derivative of KN99**a** *ago1Δ* | This study |
| JHG530 | JOHE25730 | *MATα/***a** *ago1Δ/AGO1* | BLOSSOM-generated CnTV1A$^0$ derivative of JHG499 | This study |
| JHG597 | JOHE25731 | *MAT***a** *ago1Δ::NAT* | CnTV1A$^0$ haploid progeny derived from selfing JHG530 | This study |
| JHG514 | JOHE25732 | *MAT***a** *ago1Δ::NAT* | BLOSSOM-generated CnTV1B+ derivative of KN99**a** *ago1Δ* | This study |
| JHG538 | JOHE25733 | *MAT***a** *ago1Δ::NAT* | BLOSSOM-generated CnTV1B$^0$ derivative of JHG514 | This study |
| JHG539 | JOHE25734 | *MAT***a** *ago1Δ::NAT* | BLOSSOM-generated CnTV1B$^0$ derivative of JHG514 | This study |

**Table S2. Primers used in this study.**

| Primer name | Primer Sequence (5'-3') | Purpose |
|---|---|---|
| JOHE52652/JH7 | GGTGTCATGGTCGGTATGGG | *CnACTIN* genotyping F |
| JOHE52653/JH8 | *GATACGGAGGATAGCGTGGG* | *CnACTIN genotyping R* |
| JOHE39201 | CTAACTCTACTACACCTCACGGCA | Cn*STE20*α *F*, *Genotyping MAT*α locus |
| JOHE39202 | CGCACTGCAAAATAGATAAGTCTG | Cn*STE20*α *R*, *Genotyping MAT*α locus |
| JOHE39203 | GGCTGCAATCACAGCACCTTAC | Cn*STE20***a** *F*, *Genotyping MAT***a** locus |
| JOHE39204 | CTTCATGACATCACTCCCCTAT | Cn*STE20**a** R*, *Genotyping MAT***a** locus |
| JOHE53664/JH201 | TTTACAGGTTACGGATCGTC | Cn*Ago1*_test3F, distinguish NUMT insertion |
| JOHE53665/JH202 | TACTGGTCACAGTGAGGTGG | Cn*Ago1*_test3R, distinguish NUMT insertion |
| JOHE52712/JH36 | TGTAAAACGACGGCCAGTG | M13F, to amplify Cas9 |
| JOHE52713/JH37 | GCGGATAACAATTTCACACAGG | M13R, to amplify Cas9 |
| JOHE52830/JH70 | AATTGGAGCTCCACCGCG | *CnCas9*_PCnU6/F, gRNA assembly |
| JOHE52831/JH71 | GGGAACAAAAGCTGGTACCG | *CnCas9*_sgRNA/R, gRNA assembly |
| JOHE52832/JH72 | GAGTGCTGTGGTGAAAGAGATGTTTTAGAGCTAGAAATAGCAAGTT | *CnSH1*_sgRNAF, gRNA assembly |
| JOHE52833/JH73 | ATCTCTTTCACCACAGCACTCAACAGTATACCCTGCCGGTG | *CnSH1*_PCnU6-R, gRNA assembly |
| JOHE53319/JH184 | TACCTGGGACTACTTCAAATC | CnTV1 Gag RTPCR_F |
| JOHE53320/JH185 | CCCTGCGACTCTTAGAATAG | CnTV1 Gag RTPCR_R |
| JOHE53321/JH186 | TGTGATGTCACCAGCTTCCT | CnTV1 Pol RTPCR_F |
| JOHE53322/JH187 | ATAGTGCCGAGATTCGTTTT | CnTV1 Pol RTPCR_R |
| JOHE53351/JH191 | GCTGAGCTCAGAAAGAACAAGGTTTTAGAGCTAGAAATAGCAAGTT | Cn*QIP1*_swapg1_sgRNAF, gRNA assembly |
| JOHE53352/JH192 | CTTGTTCTTTCTGAGCTCAGCAACAGTATACCCTGCCGGTG | Cn*QIP1*_swapg1_PCnu6_R, gRNA assembly |
| JOHE54349/JH209 | TAACCAGGCGTCAGTCACCA | Cn*QIP1*swapF, generate repair donor from H99 |
| JOHE54350/JH210 | TAGGGCTCACAAAGGCATCA | Cn*QIP1*swapR, generate repair donor from H99 |
| JOHE54357/JH217 | AGAAGAACAGGCAGATACAGG | Cn*QIP1*_internal_F, genotyping *QIP1* locus |
| JOHE54358/JH218 | TTTCCATCCGCCATACATAG | Cn*QIP1*_internal_R, genotyping *QIP1* locus |
| JOHE54365/JH225 | GAGGGTGTCCAGGATCATAGGTTTTAGAGCTAGAAATAGCAAGTT | Cn*QIP1*_Kog1_sgRNAF, gRNA assembly |
| JOHE54366/JH226 | CTATGATCCTGGACACCCTCAACAGTATACCCTGCCGGTG | Cn*QIP1*_Kog1_PCnU6_R, gRNA assembly |
| JOHE54367/JH227 | GTGACGACCCCGACTGGGTAGGTTTTAGAGCTAGAAATAGCAAGT T | Cn*QIP1*_Kog2_sgRNAF, gRNA assembly |
| JOHE54368/JH228 | CTACCCAGTCGGGGTCGTCACAACAGTATACCCTGCCGGTG | Cn*QIP1*_Kog2_PCnU6_R, gRNA assembly |
| JOHE54528/JH235 | ATCAGCTCGATCCTGTGATTT | Cn*QIP1*_Ko5junc_neoF, amplify NEO-containing KO donor |

| | | |
|---|---|---|
| JOHE54531/JH238 | TTAGTTGCCGCTCAGGAGTT | Cn*QIP1*_Ko3junc_neoR, amplify NEO-containing KO donor |
| JOHE54532/JH239 | CAACCTCATCGTCGGCATTA | Cn*QIP1*sw1_spanF, genotyping *QIP1* locus |
| JOHE54533/JH240 | CTTGTTCCCTCCAATCTGCA | Cn*QIP1*sw1_spanR, genotyping *QIP1* locus |
| JOHE54534/JH241 | ATGCCACCACGGTTCAAAGG | Cn*QIP1*_mut1F, genotyping *QIP1* locus |
| JOHE54535/JH242 | TCTGCATCTTCCTGCCTCCT | Cn*QIP1*_mut1R, genotyping *QIP1* locus |
| JOHE54370/JH229 | TTGGTGAGACTGGTCGTTGC | Cn*AGO1*_swapF, generate repair donor from H99 |
| JOHE54371/JH230 | ATGCCCATTGACTCCTTATT | Cn*AGO1*_swapR, generate repair donor from H99 |
| JOHE54372/JH231 | GCTAGTTCGATTAGTGTACCGGTTTTAGAGCTAGAAATAGCAAGTT | Cn*AGO1*_swapg1_sgRNAF |
| JOHE54373/JH232 | CGGTACACTAATCGAACTAGCAACAGTATACCCTGCCGGTG | Cn*AGO1*_swapg1_PCnU6_R |
| JOHE54374/JH233 | GAATGTGATGCAGCAGAACCAGTTTTAGAGCTAGAAATAGCAAGTT | Cn*AGO1*_swapg2_sgRNAF |
| JOHE54375/JH234 | TGGTTCTGCTGCATCACATTCAACAGTATACCCTGCCGGTG | Cn*AGO1*_swapg2_PCnU6_R |
| JOHE54536/JH243 | CTCAGCGTCAAGTAAGTCCG | Cn*AGO1*_mut12F, genotyping *AGO1* locus |
| JOHE54537/JH244 | ATGTGATGCAGCAGAACCAT | Cn*AGO1*_mut12R, genotyping *AGO1* locus |
| JOHE54538/JH245 | GCCACTTCCAAACAGCACCC | Cn*AGO1*sw12_spanF, genotyping *AGO1* locus |
| JOHE54539/JH246 | CCAACGACCCACCATTCACC | Cn*AGO1*sw12_spanR, genotyping *AGO1* locus |
| JOHE56190/CL48 | TACATTCCCTCTTGATCAGAGCCAGCCTTGATTTGTCCAGTCACTC TCCACTCCCAAAAAAAAACGACGGCCAGTGAATT | Cn*NUC1*_donorF, amplify NAT-containing KO donor |
| JOHE56191/CL49 | AGCACAACAGCGACGGAAACAACATGAGTAAAATCCAGAGCCCAT AGATATCATGTACATAGCTATGACCATGATTACGC | Cn*NUC1*_donorR, amplify NAT-containing KO donor |
| JOHE56192/CL50 | GCTCTGTGGACATGGGGATG | Cn*NUC1*_5_F, genotyping *NUC1* 5' junction |
| JOHE56193/CL51 | CCAGCTCACCTCCCGCAG | NAT_5_R, 5′ junction genotyping of NAT insertion |
| JOHE56194/CL52 | GCCGTTGAACCCTCAGGATC | NAT_3_F, 3′ junction genotyping of NAT insertion |
| JOHE56195/CL53 | GGCATCGTGAGGTGTAGGG | Cn*NUC1*_3_R, genotyping *NUC1* 3'junction |
| JOHE56196/CL54 | CAAGGAGGATGACGGTATTCCC | Cn*NUC1*_internal_F, genotyping *NUC1* locus |
| JOHE56197/CL55 | CAACCTTCGGCAAAAGTCCTC | Cn*NUC1*_ internal_R, genotyping *NUC1* locus |
| JOHE56198/CL56 | ATTAATCAATTTGCATTCTTGATTATACAATAAGTAAGACAGTAGTG TATACTAACAGCAAAAACGACGGCCAGTGAATT | Cn*XRN1*_donorF, amplify NAT-containing KO donor |
| JOHE56199/CL57 | CTACACCTCGGAGCTCCAGAAGATCTAAAACATAGATTTCCTCGTT TTGATGAAACCTTTAGCTATGACCATGATTACGC | Cn*XRN1*_donorR, amplify NAT-containing KO donor |
| JOHE56200/CL58 | GCTTCGAGCGGGTGTAGG | Cn*XRN1*_5_F, genotyping *XRN1* 5' junction |
| JOHE56201/CL59 | GGTGGTGAATGTCGTCTGC | Cn*XRN1*_3_R, genotyping *XRN1* 3' junction |
| JOHE56248/JH285 | TGGCTGACACTGGATGCGTTAC | Cn*XRN1*_internal2F, genotyping *XRN1* locus |

| | | |
|---|---|---|
| JOHE56249/JH286 | GGCTGATTCTGCTGCTTCTC | Cn*XRN1*_internal2R, genotyping *XRN1* locus |
| JOHE56204/CL62 | GGGCTATATCAATATTACTCCGACTTTTTTGGTATAGCCATTGAAG CCAGGGTGACAGAAAAAACGACGGCCAGTGAATT | Cn*SKI2*_donor_F, amplify NAT-containing KO donor |
| JOHE56205/CL63 | CAGTGACGACTGAAACTTTCAAGTTGCTTGCATACAATGATGGTAT GAATATGAATGCGCAGCTATGACCATGATTACGC | Cn*SKI2*_donor_R, amplify NAT-containing KO donor |
| JOHE56206/CL64 | GGGGCTATGGTACTGTCTGAGG | Cn*SKI2*_5_F, genotyping *SKI2* 5' junction |
| JOHE56207/CL65 | GCGGCTTCCTCTTACGTCC | Cn*SKI2*_3_R, genotyping *SKI2* 3' junction |
| JOHE56208/CL66 | GGTCGAGGAGGAGCGTCAATG | Cn*SKI2*_internal_F, genotyping *SKI2* locus |
| JOHE56209/CL67 | CCTCGCCTGCATACAAAAGTCAG | Cn*SKI2*_internal_R, genotyping *SKI2* locus |
| JOHE56210/CL68 | CCACCGTACTTCCATCATCAGCCATCAACTTTATCAATAGTCCTTCT GGACTTTTCAAATAAAACGACGGCCAGTGAATT | Cn*REX2*_F, amplify NAT-containing KO donor |
| JOHE56211/CL69 | ATATCCTTACAATCAATCAATGCATGATAAATGTGCAATTCAATTAC ATATCCTTCAGAAAGCTATGACCATGATTACGC | Cn*REX2*_R, amplify NAT-containing KO donor |
| JOHE56212/CL70 | CGAAATTGACGTGGCCGGC | Cn*REX2*_5_F, genotyping *REX2* 5' junction |
| JOHE56213/CL71 | GCTGAGCAACTCCATCCATTTGC | Cn*REX2*_3_R, genotyping *REX2* 3' junction |
| JOHE56214/CL72 | CGCTCATGCACTGTTGTCCAC | Cn*REX2*_INT_F, genotyping *REX2* locus |
| JOHE56215/CL73 | CCTTTGGGTACCAACGCCTAC | Cn*REX2*_INT_R, genotyping *REX2* locus |
| JOHE56216/CL74 | GTCCCTGACCAGGCAGATTTGGTTTTAGAGCTAGAAATAG | Cn*NUC1*_g1234_F, gRNA assembly |
| JOHE56217/CL75 | CAAATCTGCCTGGTCAGGGACAACAGTATACCCTGCCGGT | Cn*NUC1*_g1234_R, gRNA assembly |
| JOHE56218/CL76 | GCAGGCCTCACACTCGGTGTGTTTTAGAGCTAGAAATAGC | Cn*NUC1*_g34_F, gRNA assembly |
| JOHE56219/CL77 | ACACCGAGTGTGAGGCCTGCAACAGTATACCCTGCCGGTG | Cn*NUC1*_g34_R, gRNA assembly |
| JOHE56220/CL78 | GGTATCTCATTGGTTGGCGGGTTTTAGAGCTAGAAATAGC | Cn*XRN1*_g5141_F, gRNA assembly |
| JOHE56221/CL79 | CCGCCAACCAATGAGATACCAACAGTATACCCTGCCGGTG | Cn*XRN1*_g5141_R, gRNA assembly |
| JOHE56222/CL80 | GTTGGGTGTGATCAACTGAGGTTTTAGAGCTAGAAATAGC | Cn*XRN1*_g181_F, gRNA assembly |
| JOHE56223/CL81 | CTCAGTTGATCACACCCAACAACAGTATACCCTGCCGGTG | Cn*XRN1*_g181_R, gRNA assembly |
| JOHE56224/CL82 | AGACAAGGTGGGATGGGCAGGTTTTAGAGCTAGAAATAGC | Cn*SKI2*_g433_F, gRNA assembly |
| JOHE56225/CL83 | CTGCCCATCCCACCTTGTCTAACAGTATACCCTGCCGGTG | Cn*SKI2*_g433_R, gRNA assembly |
| JOHE56226/CL84 | GTTGGGCTGCCAGTTATGCGGTTTTAGAGCTAGAAATAGC | Cn*SKI2*_g3969_F, gRNA assembly |
| JOHE56227/CL85 | CGCATAACTGGCAGCCCAACAACAGTATACCCTGCCGGTG | Cn*SKI2*_g3969_R, gRNA assembly |
| JOHE56228/CL86 | GCGACGATGGTCCTCTCGTTTGTTTTAGAGCTAGAAATAG | Cn*REX2*_g24_F, gRNA assembly |
| JOHE56229/CL87 | AAACGAGAGGACCATCGTCGCAACAGTATACCCTGCCGGT | Cn*REX2*_g24_R, gRNA assembly |
| JOHE56230/CL88 | GTGACGTGGATGGACGACCAAGTTTTAGAGCTAGAAATAG | Cn*REX2*_g898_F, gRNA assembly |
| JOHE56231/CL89 | TTGGTCGTCCATCCACGTCACAACAGTATACCCTGCCGGT | Cn*REX2*_g898_R, gRNA assembly |
| JOHE56269/CL100 | GGGCTATATCAATATTACTCCGACTTTTTTGGTATAGCCATTGAAG CCAGGGTGACAGAAGCTGCGAGGATGTGAGCTGG | Cn*SKI2*_NEO_F, amplify NEO-containing KO donor |

| | | |
|---|---|---|
| JOHE56270/CL111 | CAGTGACGACTGAAACTTTCAAGTTGCTTGCATACAATGATGGTAT GAATATGAATGCGCGGTTTATCTGTATTAACACG | Cn*SKI2*_NEO_R, amplify NEO-containing KO donor |
| JOHE56271/CL112 | GCCACTCGAATCCTGCATGC | NEO_5_R, 5′ junction genotyping of NEO insertion |
| JOHE56272/CL113 | GGCTCCTTGTCTCTGAAACCAG | NEO_3_F, 3′ junction genotyping of NEO insertion |
| JOHE56273/CL114 | CCACCGTACTTCCATCATCAGCCATCAACTTTATCAATAGTCCTTCT GGACTTTTCAAATGCTGCGAGGATGTGAGCTGG | Cn*REX2*_NEO_F, amplify NEO-containing KO donor |
| JOHE56274/CL115 | ATATCCTTACAATCAATCAATGCATGATAAATGTGCAATTCAATTAC ATATCCTTCAGAAGGTTTATCTGTATTAACACG | Cn*REX2*_NEO_R, amplify NEO-containing KO donor |
| JOHE56250/JH287 | CAAGGCGGCGAACTCTATCA | CnTV1_Pol_qPCRF, qPCR |
| JOHE56251/JH288 | TCGGGCACAAACTCTGTATCTG | CnTV1_Pol_qPCRR, qPCR |
| JOHE52657/JH12 | ATCGTTCTTGACTCTGGTGACGGT | Cn*ACT1* qPCRF, qPCR |
| JOHE52658/JH13 | AAGTGGTGAAGAGGTAACCACGCT | Cn*ACT1* qPCRR, qPCR |
| JOHE56414/JH308 | ATTGGGAGGCTGTGGTAAAG | Cn*XRN1*_qRT1F, qPCR |
| JOHE56415/JH309 | CAGGGTCGTAAACAAATTGAG | Cn*XRN1*_qRT1R, qPCR |
| JOHE56590/JH310 | CAGTATGTTTAGCAGTACCCGTAA | *CNAG_07650*-qRT1F, qPCR |
| JOHE56591/JH311 | GACCTCTTGAATCTCGCCTC | *CNAG_07650*-qRT1R, qPCR |
| JOHE56949/JH426 | GATTACGCCAAGCTTTTTGGTGATTGGCCGGCATACGT | CnTV1_GSP2, RACE PCR |
| JOHE56950/JH427 | GATTACGCCAAGCTTGCCGCCTTGGCAAAGGCGTCTTT | CnTV1_GSP1, RACE PCR |
| JOHE56951/JH428 | GATTACGCCAAGCTTACCGCGTGCTATTGAGGCGAATG | CnTV1_NGSP2, RACE PCR |
| JOHE56952/JH429 | GATTACGCCAAGCTTGACGCCGGGCTTTATTGGGCTAA | CnTV1_NGSP1, RACE PCR |
| JOHE56969/JH430 | GATTACGCCAAGCTTTTTGCTTGGCAGCTCCGCTCTCAG | CnTV1_GSP_B1, RACE PCR |
| JOHE56970/JH431 | GATTACGCCAAGCTTGGCGATGACAGGGGCGGCAATAGC | CnTV1_GSP_B2, RACE PCR |
| JOHE56971/JH432 | GATTACGCCAAGCTTATCCGGATCTGCCGTGTACGCCAT | CnTV1_GSP_C1, RACE PCR |
| JOHE56972/JH433 | GATTACGCCAAGCTTGAGAGGGCTGAAGCGTGCGTACGA | CnTV1_GSP_C2, RACE PCR |
| JOHE57937/JH436 | CGTAGGCTGGATACTAAGAG | Cn*AGO1*_swap1F, generate repair donor from NRHc5010 |
| JOHE57938/JH437 | GTCTTGATGACGGGGTTGATAATGTCATAGTGGCTGATAAAT | Cn*AGO1*_swap1R, generate repair donor from NRHc5010 |
| JOHE57939/JH438 | ATCAACCCCGTCATCAAGAC | Cn*AGO1*_swap2F, generate repair donor from NRHc5010 |
| JOHE57940/JH439 | CAGCCACTGTAGTCAAGATTTAGA | Cn*AGO1*_swap2R, generate repair donor from NRHc5010 |

**Legends for Datasets S1 to S4**

**Dataset S1.** Pairwise amino acid identities of CnTV1 Gag and Pol proteins determined by BLASTP.

**Dataset S2.** Candidate variants in five CnTV1-free chemically cured derivatives of NRHc5010.

**Dataset S3.** GO term enrichment analysis of the 294 shared differentially expressed genes between CnTV1B$^+$ versus KN99**a** *ago1*Δ and CnTV1B$^+$ versus CnTV1B$^0$.

**Dataset S4.** Metadata for sequences used in the CnTV1 phylogenetic analysis.
